# A Survey and Systematic Assessment of Computational Methods for Drug Response Prediction

**DOI:** 10.1101/697896

**Authors:** Jinyu Chen, Louxin Zhang

## Abstract

Drug response prediction arises from both basic and clinical research of personalized therapy, as well as drug discovery for cancer and other diseases. With gene expression profiles and other omics data being available for over 1000 cancer cell lines and tissues, different machine learning approaches have been applied to solve drug response prediction problems. These methods appear in a body of literature and have been evaluated on different datasets with only one or two accuracy metrics. We systematically assessed 17 representative methods for drug response prediction, which have been developed in the past five years, on four large public datasets in nine metrics. This study provides insights and lessons for future research into drug response prediction.

## 1 Introduction

Cancer is still an incurable disease. Recent cancer genome studies have suggested that each cancer patient owns a unique profile of gene mutations and that there is no silver bullet for defeating all the cancers. Therefore, how to use the genomic profiles of cancer patients to design a personalized treatment is vital for effective control of disease progression [1, 2]. This requires accurate prediction of the patients’ response to target-specific drugs and therapies.

Given the paucity of patients’ molecular profiles and their responses to drugs, large-scale drug screening experiments on cultured and genome-wide profiled cancer cell lines have been used to study drug response [3, 4]. The US National Cancer Institute (NCI) 60 human tumor cell lines screen database provides drug screening data for thousands of drugs in 60 tumor cell lines, for which mRNA, microRNA expression, DNA methylation and mutation profiles are also available [3, 5]. In 2012, another two large-scale pharmacogenomic datasets were made publicly available: the Cancer Cell Line Encyclopedia (CCLE) [6] and the Genomics of Drug Sensitivity in Cancer (GDSC) [7]. The CCLE contains drug sensitivity data from over 11,000 experiments involving 24 drugs and 504 cell lines and both the gene mutational and expression profiles for about 1000 cell lines. The GDSC includes the data generated from about 200,000 drug response experiments that tested 266 drugs on nearly 1000 human cancer cell lines.

With a large amount of drug response data for cell lines being available, various machine learning approaches have been applied to design computational methods for drug response prediction in recent years. These prediction methods, their interpretability and the importance of the integration of multiple omics profiles for performance improvement are well documented in two survey articles [8, 9]. Additionally, key methods developed before 2014 were evaluated in [10] and [11]. Since 2015, a number of new approaches have been developed for drug response prediction, including regression methods with or without kernels, [12–16], Bayesian inference methods [17–20], matrix-factorization-based methods [21–23] and other miscellaneous (deep) machine learning methods [24–28].

In this study, we aim to systematically assess 17 representative computational methods that have been developed since 2015 (Table 1) and the classic regression approaches on four public datasets: GDSC, CCLE, NCI-60 and the Cancer Therapeutics Response Portal (CTRP) dataset [29]. These methods have often been evaluated by the developers on the datasets that were obtained from the aforementioned datasets in one or two metrics. It remains therefore unclear which are among the best methods. We evaluated these state-of-the-art methods on the same datasets in nine metrics. On the basis of these analyses, we draw lessons and insights that could hopefully benefit future research into drug response prediction.

**Table 1.**
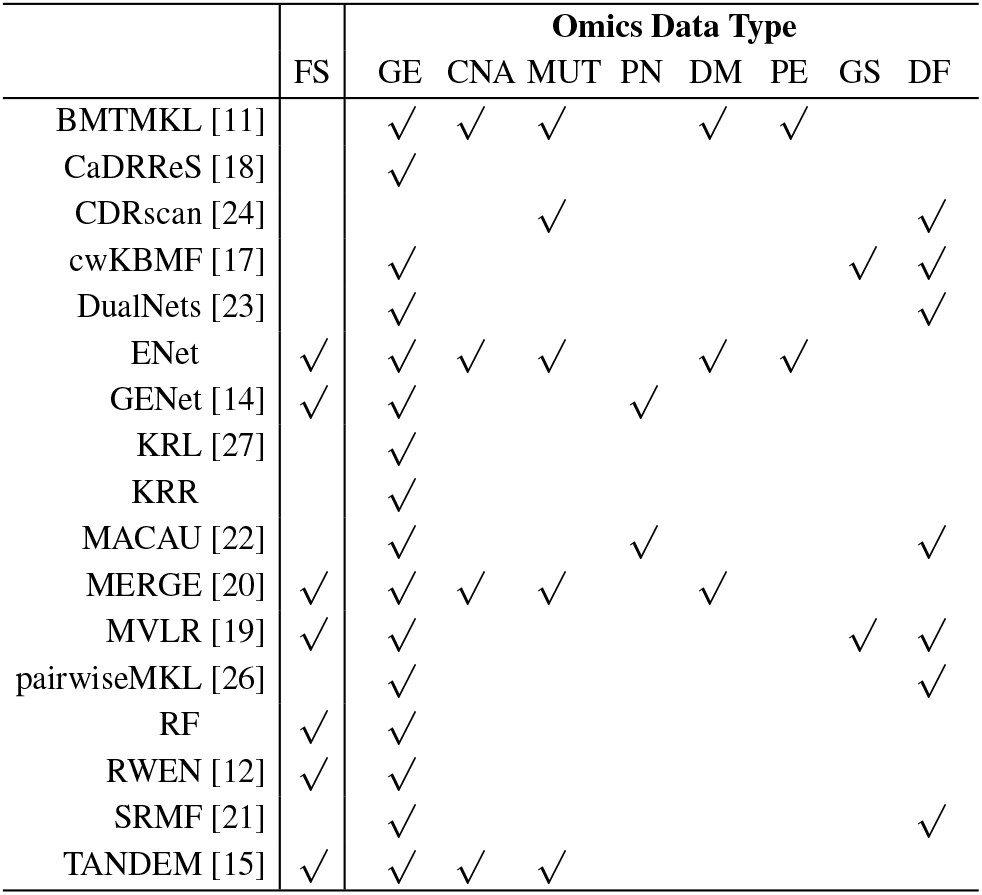
Summary of key types of omics data that can be integrated in the 17 assessed methods (Section 2.4–2.7) in addition to drug response data. GE: Gene expression; CNA: Copy number aberration; MUT: Mutational profile; PN: Cellular protein network; DM: DNA methylation; PE: Protein expression; GS: Gene similarity profile; DF: Drug features. FS: Feature selection, indicating whether a method has a feature selection function or not.

|  | FS | Omics Data Type |  |  |  |  |  |  |  |
| --- | --- | --- | --- | --- | --- | --- | --- | --- | --- |
|  |  | GE | CNA | MUT | PN | DM | PE | GS | DF |
| BMTMKL [11] |  | ✓ | ✓ | ✓ |  | ✓ | ✓ |  |  |
| CaDRReS [18] |  | ✓ |  |  |  |  |  |  |  |
| CDRscan [24] |  |  |  | ✓ |  |  |  |  | ✓ |
| cwKBMF [17] |  | ✓ |  |  |  |  |  | ✓ | ✓ |
| DualNets [23] |  | ✓ |  |  |  |  |  |  | ✓ |
| ENet | ✓ | ✓ | ✓ | ✓ |  | ✓ | ✓ |  |  |
| GENet [14] | ✓ | ✓ |  |  | ✓ |  |  |  |  |
| KRL [27] |  | ✓ |  |  |  |  |  |  |  |
| KRR |  | ✓ |  |  |  |  |  |  |  |
| MACAU [22] |  | ✓ |  |  | ✓ |  |  |  | ✓ |
| MERGE [20] | ✓ | ✓ | ✓ | ✓ |  | ✓ |  |  |  |
| MVLR [19] | ✓ | ✓ |  |  |  |  |  | ✓ | ✓ |
| pairwiseMKL [26] |  | ✓ |  |  |  |  |  |  | ✓ |
| RF | ✓ | ✓ |  |  |  |  |  |  |  |
| RWEN [12] | ✓ | ✓ |  |  |  |  |  |  |  |
| SRMF [21] |  | ✓ |  |  |  |  |  |  | ✓ |
| TANDEM [15] | ✓ | ✓ | ✓ | ✓ |  |  |  |  |  |

## 2 Brief overview of drug response prediction and machine learning methods

### 2.1 The problems of drug response prediction

Drug response in cell lines are often measured in terms of either the half maximal inhibitory concentration (IC_50_ or GI_50_). Both IC_50_ and GI_50_ represent how much of a particular drug is needed to inhibit a cell activity and the cell growth by 50%, respectively. Another quantitative measure for drug response is the area under the concentration response curve (AUC) [30]. For IC_50_, GI_50_ or AUC, the lower the value is, the more effective is the drug. The response to *m* drugs in *n* cell lines are given as an *n × m* real matrix, in which the (*i, j*)-th entry is the IC_50_, GI_50_ or AUC value of the *j*-th drug in the *i*-th cell line.

Drug response prediction is a supervised learning problem: given a training drug response dataset *R*_*n×m*_ with *n* cell lines and *m* drugs, a response function *r*(*x, y*) is learned from *R*_*n×m*_ and then used to compute the response to (i) a known drug (in the training dataset) in a new cell line, (ii) a new drug in a known cell line or (iii) a new drug in a new cell line. In this study, we assess different methods for the Type-i drug response prediction. The Type-iii drug response prediction is believed to be more challenging than the first two.

### 2.2 Integration of omics data

The rationale for the predictability of drug response is that the response to drugs that have similar chemical properties in a cell line are highly related and cell lines that have similar genomic profiles also have similar drug response to a drug. Good methods for drug response prediction rely therefore on extra omics data other than the drug response matrix. The omics data that are frequently used for the purpose include:

- Gene expression profiles, which characterize the expression level of a set of genes in a cell line. Currently, both microarray and RNA-Seq gene expression datasets are available in the aforementioned datasets.
- MicroRNA expression profiles. MicroRNAs are non-coding RNA molecules consisting of about 20 nucleotides. They function in RNA silencing and post-transcriptional regulation of gene expression.
- Copy number alteration/aberration (CNA) profiles. Copy number alterations are changes in copy number for genes in a (tumor) cell line.
- Mutational profiles. These refer to a list of mutation types (e.g., base substitutions, short insertions and deletions) and mutation counts in a (tumor) cell line.
- DNA methylation profiles. DNA methylation is tissue-specific and dynamic. The DNA methylation profiles characterize the patterns of aberrant DNA methylation occurring in a (tumor) cell line.
- Gene similarity profiles. These comprise curated data characterizing functionally and structurally similarities of a set of genes, which are often computed from genomic and/or proteomic profiles.
- Cellular protein networks. These could be cell-specific protein–protein interaction networks, gene regulatory networks, or signaling pathways.
- Protein expression profiles. These characterize the expression levels of proteins in a cell line. Protein expression levels can be measured on the reverse phase protein lysate microarray or the global mass spectrometry platform.
- Drug features. There are two main types of drug features. One is information on the drug’s primary targets or target pathways, which are very useful for drug classification. Another is the molecular structural information. Each drug can be characterized by different 1-dimensional and 2-dimensional structural properties, such as the number of atoms, the presence of chemical substructures, etc.

By concatenating different omics datasets and encoding non-numeric entries into numbers, we can represent multiple omics profiles of *n* cell lines into a real-valued matrix *C*_*n×s*_, in which each column corresponds to a feature in one of the omics profiles. Similarly, we can also represent multiple profiles of *m* drugs by a real-valued matrix *D* of *m* rows and integrate the relationships among *k* genes such as a gene similarity profile and a cellular protein network into a *k×k* matrix *G*. Thus, mathematically, the training dataset for drug response prediction includes a cell line– drug response matrix *R*_*n×m*_ and at most three auxiliary profile matrices: *C*_*n×s*_, *D* and *G*.

Table 1 summarizes which omics data are used by each of the methods to be evaluated in this work.

### 2.3 Single-task vs multi-task learning

A linear regression model assumes a linear relationship between the drug response level and the genomic profile of cell lines. In other words, the linear regression approach learns a linear function *r*(*x*) that maps the profiles *x* of cell lines to the drug response levels. It often identifies good bio-markers. However, it has relatively low prediction accuracy particularly when there are limited samples for a drug, which is also confirmed by this study.

Recently, many methods have been developed to predict the response of multiple drugs in a single model [11, 17, 19–21, 26]. In this so-called multi-task learning framework [31], sharing information on different drugs (tasks) often enables us to obtain a prediction method with high accuracy. It is easy to find out which methods are developed in the multi-task learning framework.

In the rest of this section, we groups different methods into four groups depending on their underlying mathematical principle.

### 2.4 Linear regression methods and their generalizations

Different regression methods have been developed for drug response prediction. They use the drug response vector *R*_*n×*1_ of a drug *d* in the *n* cell lines to learn a linear response function *r*_*d*_(**x**) = **w** *•* **x** with the coefficient vector **w** = (*w*_1_, *w*_2_, *…, w*_*s*_), where **x** = (*x*_1_, *x*_2_, *…, x*_*s*_)^*T*^ represents the profile vector of a cell line. For a new cell line with the profile vector **c**, *r*_*d*_(**c**) is output as the response to the drug in the cell line.

A linear regression method computes the coefficient vector **w** of the function *r*_*d*_(**x**) by solving the following optimization problem:

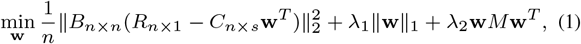

where ||*•*||_1_ and ||*•*||_2_ are, respectively, the *L*_1_-and *L*_2_-norm defined for real vectors; *M* is either the *s × s* identity matrix or an *s × s* symmetric semi-definite matrix derived from an input profile matrix; *T* denotes the transpose operator for matrices; and *B*_*n×n*_ is a diagonal matrix with *n* parametric diagonal entries *b*_*j*_. The free parameters *λ*_*i*_’s and *b*_*j*_’s are set differently for different methods.

#### Elastic net (ENet)

In this method, *M* = **I**_*s×s*_ and *B*_*n×n*_ = *I*_*n×n*_ (i.e., *b*_*i*_ = 1) in Eqn. (1) [32]. Thus, the third term of the equation simply becomes the *L*_2_-norm of the coefficient vector **w**. Additionally, the ENet uses a 5- or 10-fold cross-validation procedure to determine the values of *λ*_1_ and *λ*_2_.

The implementation of the ENet available in the R package has the time complexity *O*(*s*^2^*n* + *s*^3^), where *s* and *n* are the number of features and cell lines in the input cell line profile matrix, respectively.

#### Generalized ENet (GENet)

In this method, *B*_*n×n*_ = *I*_*n×n*_ in Eqn. (1). We assessed two versions of this model: **GENet-Lap** and **GENet-NLap** in which *M* is set to the Laplacian matrix and the normalized Laplacian matrix of a gene interaction network [14], respectively.

In [14], the authors proposed to solve the optimization problem in Eqn. (1) through cyclic coordinate descent [33] by changing one coefficient *w*_*k*_ at a time, while keeping the values of other coefficients *w*_*j*_ (*j ≠ k*) unchanged. For this computational method, the time complexity of each iteration is *O*(*nms*^3^).

#### Response-weighted ENet (RWENet)

This approach uses an iterative procedure to set (*b*_*j*_) and compute the coefficient vector **w** in Eqn. (1). Here, the drug response data 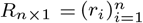 is assumed to follow a normal distribution *N* (*µ, σ*^2^) with the parameters being estimated as [12]:

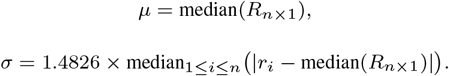

Under this assumption, the drug response value is less than ℓ (resp. larger than *γ*) with a probability of 0.05, where ℓ = *µ −* 1.65 *× σ* and *γ* = *µ* + 1.65 *× σ*.

In the RWENet method, *b*_*j*_ is initially set to 1, that is, 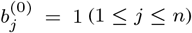. Thus in the first step, ENet is simply called to compute the coefficient vector **w**^(0)^. In the *t*-th iteration step, 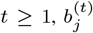 are set as follows, depending on whether or not *r*_*j*_ *≤* ℓ with the current coefficient vector **w**^(*t−*1)^:

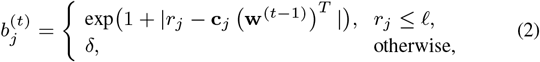

where *δ* is a small fixed constant close to 0 and **c**_*j*_ is the *j*-th row of *C*_*n×s*_. The purpose of setting (*b*_*j*_) in this way is to find a coefficient vector that has a good prediction for cell lines with low drug responses, which are sensitive to the drug. The algorithm continues until the difference between the input drug responses and the predicted values are small enough. This variant is called the RWENet-left method. The time complexity of the RWENet-left method is *O*(*ms*^3^ + *mns*^2^ + *ms*) for each iteration.

Symmetrically, the method can be modified for predicting drug responses for cell lines that are resistant to a drug through setting *r*_*j*_ ≥ γ in Eqn. (2), as well as for predicting the responses for cell lines that are either sensitive or resistant to a drug. These two variants of RWENet are called the RWENet-right and RWENet-both methods [12], respectively.

#### MERGE (mutation, expression hubs, known regulators, genomic

Lee *et al.* implemented a linear regression method by incorporating multi-omics prior information about genes for all the drugs simultaneously [20]. They used cell line profiles *C*_*n×s*_ = (*c*_*ij*_) for inference and gene profiles *D*_*s×k*_ = (*d*_*ij*_) for assigning scores for the *s* genes. Lee *et al.* inferred the regression coefficient matrix *W*_*s×m*_ = (*w*_*ij*_) by solving the following optimization problem:

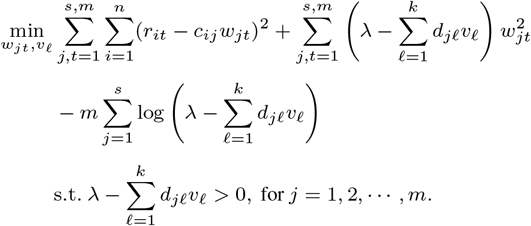

Here, *λ* is a hyperparameter to be determined on the basis of the data and can be considered as the upper bound of the gene scores 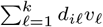. Thus, the second term is used to penalize genes with lower scores.

#### Kernel ridge regression (KRR)

This method first maps a cell line vector *v* to a vector *f* (*v*) in a high-dimensional feature space and then computes the coefficient column vector **w** in the image vector space [27, 34]. More specifically, it solves the following optimization problem:

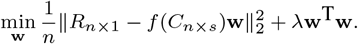

This problem can be solved as easily as solving Eqn. (1). The reason is that one only needs to know how to compute the inner product *f* (**u**) · *f* (**v**) of image vectors to solve the optimization problem. He *et al.* used the so-called Gaussian kernel function to define the inner product of the vectors *f* (**u**) and *f* (**v**) [27]:

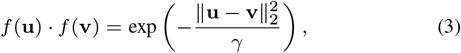

where *γ* is a free parameter.

#### Pairwise multiple kernel learning (pairwiseMKL)

When drug profiles are available, it is possible to map a cell line profile *v*_*c*_ to *f* (*v*_*c*_) in a new feature space *H*, map a drug profile *u*_*d*_ to *g*(*u*_*d*_) in another new feature space *H*′ and then find a linear regression function in the product space *H × H*′ to predict drug response [26].

Let *k*_*c*_ and *k*_*d*_ be the two kernel functions such that:

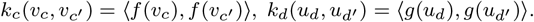

Define 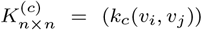, 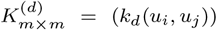 and their Kronecker product 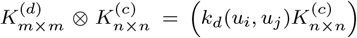. In the Kronecker product, each drug-cell line pair (*u*_*d*_, *v*_*c*_) corresponds to a row and a column. Hence, 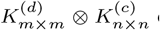 can be used to define a linear regression function in *H × H*′, by which the drug response *r*_*uv*_ to a drug *u* in a cell line *v* can be predicted as:

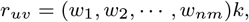

where *k* is the column vector consisting of the kernel values between each training drug-cell line pair (*u*_*d*_, *v*_*c*_) and the test pair (*u, v*). Here, the coefficient row vector **w** = (*w*_1_, *w*_2_, *…*, *w*_*nm*_) can be found by solving the following system of linear equations:

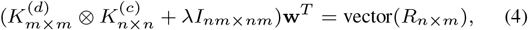

where vector 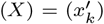 is the flat form of an *n × m* matrix *X* that is defined by 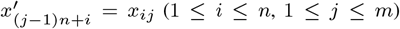 and *λ* is a hyperparameter to be determined.

Cichonska et al. incorporated a few ideas into the above kernel regression approach. First, they proposed to use both multiple kernels 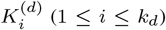 for drugs and multiple kernels 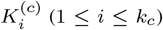 for cell lines. In this way, 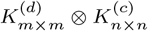 in Eqn. (4) is replaced by a weighted combination of the pairwise Kronecker products:

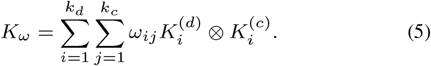

Second, the authors applied the central kernel alignment technique [35] to compute the kernel mixture weight matrix *ω* = (*ω*_*ij*_) in Eqn. (5).

Let 1_*nm*_ be the matrix of size *nm* in which each entry is 1 and 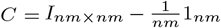, which satisfies that *C*^2^ = *C*. In the central kernel alignment approach, the kernel function *K*_*ω*_ in Eqn. (5) is normalized into 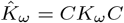. Let

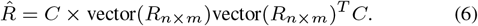

The weight matrix (*ω*_*ij*_) is computed by maximizing the similarity between 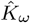 and 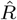:

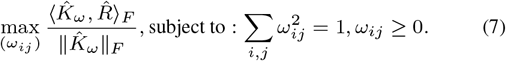

Third, Cichonska et al. proposed to apply a Gaussian response kernel to each drug response *r*_*ij*_ such that *r*_*ij*_ is replaced by a vector of size *nm* that corresponds to a histogram vector of the Gaussian distribution centered around *r*_*ij*_ ([36]). In this way, vector(*R*_*n×m*_) in Eqn. (6) is replaced by an *nm × nm* matrix.

Lastly, one disadvantage of the pairwise multiple kernel regression approach is its high time complexity. Cichonska et al. used the bilinearity of the Kronecker product of matrices to simplify the time complexity of solving the optimization problems in Eqn. (4) and (7). In their implementation of the conjugate gradient method that iteratively solves the optimization problems, each iteration takes *O*(*k*_*d*_*k*_*c*_*nm*(*n* + *m*)) time.

### 2.5 Bayesian inference methods

Bayesian inference is a statistic method in which Bayes’ theorem is used to update the probability for a model as more data becomes available. Bayesian inference methods for drug response predication often take multi-task learning approach. They attempt to predict all the responses *r*(*i, j*) (1 *≤ i ≤ n*; 1 *≤ j ≤ m*) through jointly learning a single model from the training response matrix *R*_*n×m*_ and auxiliary cell line and drug profile matrices.

#### The Bayesian multi-task multi-kernel learning method (BMTMKL)

This multi-task learning method had the best performance in the 2014 NCI-DREAM drug sensitivity prediction challenge [11].

For a matrix *X*, we use *X*[*i, \**] and *X*[*\*, j*] to denote its *i*-th row and *j*-th column, respectively. We can rewrite the response matrix *R*_*n×m*_ as follows:

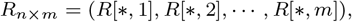

where *R*[*\*, j*] is the responses to drug *j* in the cell lines. Costello *et al.* proposed to decompose *R*[*\*, j*] into a linear combination of *p* components 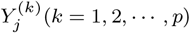 as follows [11]:

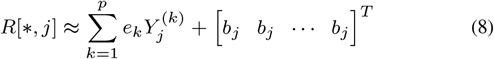

for each *j* = 1, 2, *…, m*; equivalently:

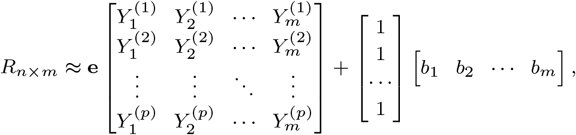

where **e** = [*e*_1_ *e*_2_ *… e*_*p*_] *∈* ℝ^1*×p*^ is common to all drugs, and both 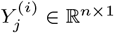 and *b*_*j*_ depend only on drug *j* for each pair of *i* and *j*.

One may consider that there are *p* different profiles of cell lines (such as the DNA methylation profiles or gene expression profiles) and 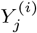 is the response component of drug *j* that is inferred from the *i*-th cell line profile for each *j* = 1, 2, *…, m* and *i* = 1, 2, *…, p*. Costello *et al.* assumed the following connection between 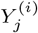 and cell line profiles 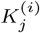:

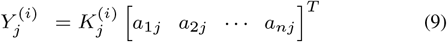

for each drug *j*, where 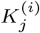 is the *n × n* similarity matrix that is computed via the Gaussian kernel function in Eqn. (3) or other similarity function.

Costello *et al.* inferred drug responses by using Bayesian inference. They assumed that each *e*_*i*_ of the parameter vector **e** follows a normal distribution 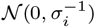 such that *σ*_*i*_ follows a Gamma distribution:

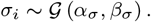

Similarly, for each drug *j* = 1, 2, *…, m*, each *b*_*j*_ is sampled from a similar distribution:

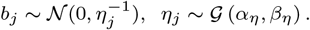

Each component of the vector (*a*_1*j*_, *a*_2*j*_, …, *a*_*nj*_) is independently sampled from the following distribution:

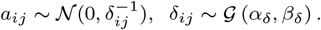

Costello *et al.* further assumed that the response components 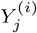 are sampled from the following multivariate normal distributions:

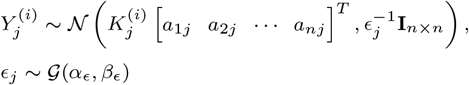

and

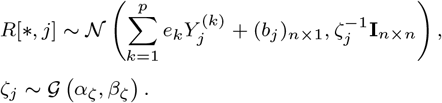

In summary, this is a very general stochastic model involving the following 10 parameters for prior Gamma distributions:

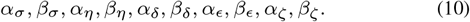

Costello *et al.* applied their method to an instance where the training dataset involved 28 drugs and 35 cell lines and the test data included 18 cell lines. They set the number of components (*p*) to 22 by considering different combinations of input cell line profiles. It is worth pointing out that they used a form that is more general than Eqn. (9) so that the responses data do not necessarily involve the same set of cell lines for different drugs.

#### Component-wise kernel Bayesian matrix factorization (cwKBMF)

This was developed on the basis of a matrix factorization that was more general than Eqn. (8)–(9) [17]. Ammad-ud-din et al. took drug profile into account through a drug similarity matrix.

#### Multi-view multi-task linear regression (MVLR)

This method approaches the linear regression model via Bayesian inference. For *m* drugs targeting one or more common pathways, a linear regression model assumes the following relationship between the drug responses *R*_*n×m*_ and cell line profiles *C*_*n×s*_:

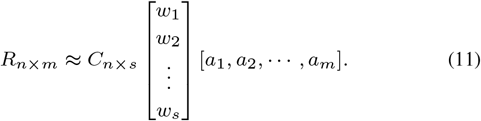

In [19], the cell line profile was divided into *p* parts, i.e.,

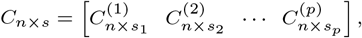

where Σ_1*≤i≤p*_ *s*_*i*_ = *s*. The rationale of this approach is that each 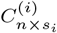 represents a data view, generated from related features; the corresponding *w*_*i*_’s form the regression coefficient vector for each data view; and *a*_*i*_ is a parameter being set for each drug *i*. Note that if *p* = 1, *s*_1_ = *s*, Eqn. (11) is equivalent to the relationship between *R*[*\*, j*] and *C*_*n×s*_ on which the linear regression model Eqn. (1) is based for each *j*.

Ammad-ud-din *et al.* inferred the parameters (*w*_*j*_) (1 *≤ j ≤ s*) and *a*_*i*_ (1 *≤ i ≤ m*) by assuming the following distributions [19]:

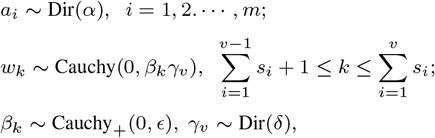

and

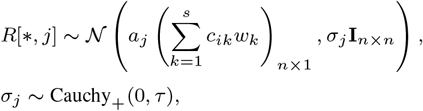

in which there are four free parameters (*α, ϵ, δ, τ*) for the Dirichlet prior

#### MACAU

This is a Bayesian matrix factorization learning method [37]. Yang et al. used this method on drug target information and perturbed pathway information for drug response prediction [22]. The drug response matrix *R*_*n×m*_ is assume to has the following factorization:

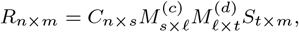

where *t* is the number of drug targets, *s* is the number of perturbed pathways, ℓ is a hyperparameter of the method. Here, 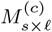 and 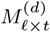 are two so-called link matrices that project the side information about cell lines and drugs into the latent matrix factors of the drug response matrix *R*_*n×m*_, respectively.

Macau uses Gibbs sampling to sample both the latent vectors and the link matrices. Having the time complexity *O*(*st*), it can support high-dimensional side information when conjugate gradient based noise injection sampler is employed [37].

### 2.6 Matrix-factorization based methods

#### Predicting cancer drug response using a recommender system (CaDRRes) as follows

This method was developed under the assumption that the drug response function can be approximated by the sum of a few low-rank matrices. More specifically, the response matrix *R*_*n×m*_ for *m* drugs and *n* cell lines has the following decomposition [18]:

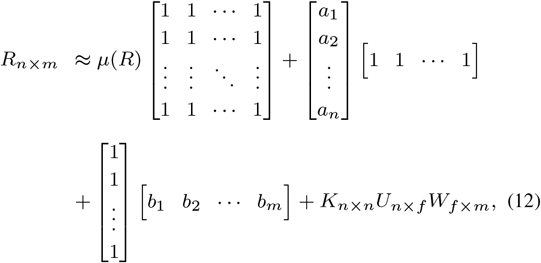

where *K* is a similarity matrix of the cell lines, *f* is a small positive integer to be determined, and *µ*(*R*) is the average value of entries in *R*_*n×m*_.

Let 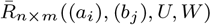 denote the right-hand side of Eqn. (12). Suphavilai *et al.* solved the following minimization problem by using the gradient descent method to predict drug responses [18]:

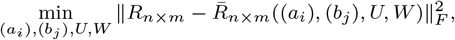

where *K* is calculated via the Pearson’s correlation of gene expression profiles for cell lines and *f* = 10.

#### DualNets

Zhang *et al.* developed this simple but efficient network interpolation method for drug response prediction [23]. They assumed that the response to a known drug *d* in a new cell line is a weighted combination of the responses of the neighboring cell lines *c*_*i*_ (*i* = 1, 2, *…, n*) to drug *d* as given below:

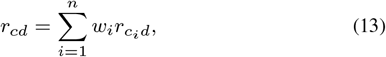

where 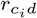 is the response level to drug *d* in cell line *c*_*i*_ and the coefficient *w*_*i*_ is defined as a negative exponential function. Let *s*(*c*, *c*_*i*_) be the gene expression correlation between the two cell lines *c* and *c*_*i*_. Then *w*_*i*_ is given as:

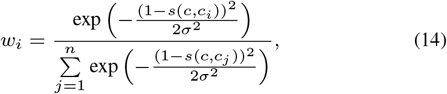

where *σ* is the decay rate parameter.

Similarly, the response value of a known cell line to a new drug could be inferred by using drug similarity.

#### The similarity-regularized matrix factorization (SRMF) method

This method simply assumes that the drug response matrix *R*_*n×m*_ is approximately a product of two low-rank matrices *U*_*n×k*_ (a compression of the cell line profile matrix) and *V*_*m×k*_ (a compression of the drug profile matrix). Wang *et al.* used a cell line similarity matrix *S*_cell_ and a drug similarity matrix *S*_drug_ to compute the two low-rank matrices [21] as follows:

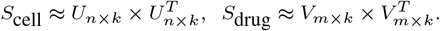

In this way, Wang *et al.* proposed to compute *U* and *V* by solving the following optimization problem:

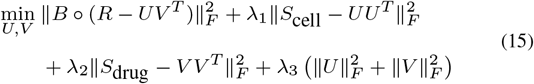

where *B*_*n×m*_ = (*b*_*ij*_) is the indicator matrix such that *b*_*ij*_ = 0 if the drug response value *r*_*ij*_ is missing and 1 otherwise, and the operator “*o*” is the entry-by-entry matrix product used to mask off the missing entries in *R*.

### 2.7 Miscellaneous methods

#### Random forest with gene filtering (RF-g) and with both gene and sample filtering (RF-gs)

Random forest is a regression approach that captures nonlinear relationships between predictors and response variables [38]. It first uses bootstrapping to split the data into several sub-datasets, fits a decision tree to each of them, and finally averages the results. In order to improve the accuracy of the drug response predictions, Riddick *et al.* incorporated the procedures of filtering gene signatures and removing cell line outliers in the random forest framework [39].

#### Kernel rank learning (KRL)

This method focuses on accurately recommending the *k* most sensitive drugs for each cell line [27]. It therefore proposed to use the normalized discounted cumulative gain (DCG) defined in Eqn. (23) to construct the loss function instead of the square of the loss between observed and predicted values:

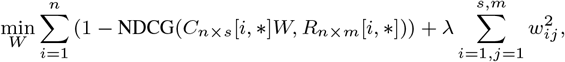

where *W* is a real *s × m* matrix. The drugs are recommended on the basis of the predicted rank vector *xW* for the cell line with the profile *x*.

#### TANDEM

When applied to multiple omics data of cell lines, the ENet or other regression methods make drug response prediction mainly on the basis of gene expression profile. To take advantage of information contained in other profiles, Aben et al. developed a two-stage approach to predict drug response, named TANDEM [15]. In this approach, an ENet model is first used to predict drug response using mutation, CNA, cancer type and methylation profiles and then the second ENet model is used to adjust the prediction made in the first stage using the gene expression profile.

#### Cancer drug response profile scan (CDRscan)

CDRscan is a deep convolutional neural network approach for drug response prediction [24]. CDRscan employs a two-stage convolution architecture, in which the genomic mutational profiles of cell lines and the molecular profiles of drugs are processed separately in Stage 1 and then merged by simulating the ‘docking’ process in Stage 2.

## 3 Evaluation Metrics

In this study, we chose nine widely used metrics to evaluate the 17 methods on the basis of the difference between the observed responses, 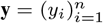, and the predicated responses, 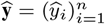, of *n* cell lines to a drug.

### Root mean square error (RMSE)

The basic RMSE attempts to capture the deviation between the true and predicated responses for a method and is defined as:

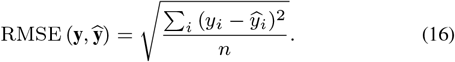

To facilitate the comparison of two prediction methods with different scales, one may use the following normalized RMSE:

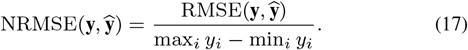

For a drug, one may be concerned only with the sensitive cell lines only or the resistant cell line only. For these purposes, RMSE is further modified as [12]:

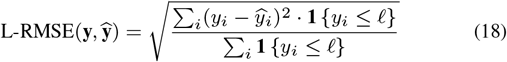

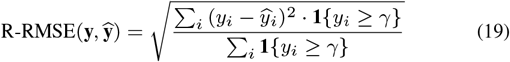

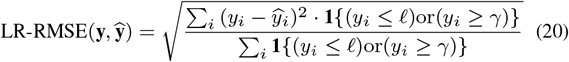

where **1**{} denotes the indicator function,

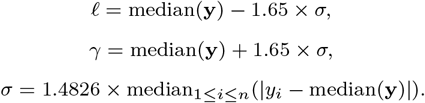

Note that ℓ and *γ* are the left and right cutoff points, respectively, and are computed by the lower and upper limit of the 90% confidence interval for the data following a Gaussian distribution with standard deviation *σ*.

### Pearson correlation coefficient (PCC)

Let *µ*(*X*) denote the mean of a random variable. The PCC of **y** and 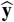 is defined as:

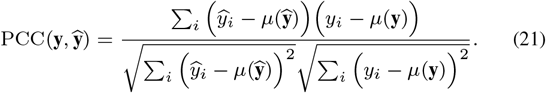

### Spearman correlation coefficient (SCC)

Assume that 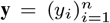 is sorted into 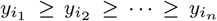 in decreasing order. The rank of the response 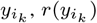, is defined to be *k*. The integral vector 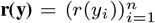 is called the rank vector of **y**. The rank vector of 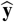 can be defined similarly. The SCC of **y** and 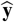 is then defined as:

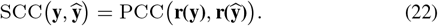

### Normalized DCG (NDCG)

This is a widely used score for ranking quality [18, 40]. The DCG for the response prediction made by a method is defined as:

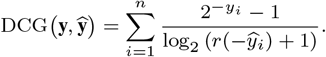

The numerator gives more weight to high sensitivity score whereas the denominator gives preference to high ranks. The normalized DCG is further defined as:

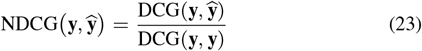

### Normalized weighted probabilistic c-index (NWPC)

In the 2014 NCI-DREAM drug sensitivity prediction challenge [11], NWPC was used to evaluate the performance of a method. It measures the agreement between a predicted ranked list and the true list, defined as follows:

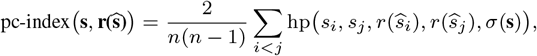

where *σ*(*X*) is the standard deviation of a random variable *X* and the function 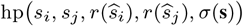, *σ*(**s**)) is defined as

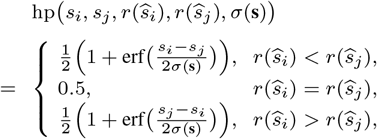

and 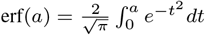.

For a method, the pc-index is calculated separately for each drug *d*, denoted 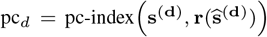 where s^(**d**)^,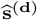 are the true and predicted score vectors for the drug d, respectively. The final score of a method *M* is calculated as the weighted average of the pc-index scores across all the drugs (i.e. the wpc-index) as follows:

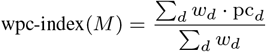

The weight *w*_*d*_ is calculated from the empirical null distribution of pc_*d*_. For each drug *d*, (i) produce a random ranking of *n* items, denoted 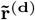, and calculate the pc-index 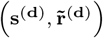; (ii) repeat this procedure 10,000 times to construct an empirical null distribution with a median of *µ*_*d*_ and a standard deviation of *σ*_*d*_; (iii) compute the pc-index with the standard ranking **r**(**s**^(**d**)^), 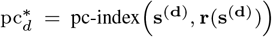; (iv) drug weight 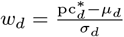.

In order to scale the range of the wpc-index into [0, 1], use the transformation:

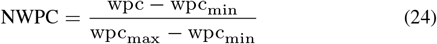

where wpc_max_, wpc_min_ are obtained from the real ranking vector **r**(**s**^(**d**)^) and its inverse, respectively, for each drug when computing pc_*d*_.

Unlike the previous eight metrics, the wpc-index measures the overall performance for all the predicted drugs. Therefore, only one value is given for each method.

Lastly, we point out that a smaller value indicates good performance for the first five metrics introduced above, whereas a value close to 1 suggests good performance for the last four metrics.

## 4 Test datasets

We assessed the selected methods with four public drug response datasets that were respectively downloaded from:

- The GDSC website (https://www.cancerrxgene.org/downloads, accessed 1 Feb. 2019),
- the CCLE website (https://portals.broadinstitute.org/ccle/data, accessed 1 Aug. 2019),
- the NCI-60 (version 2.6.0) [5] and
- The CTRP website (https://portals.broadinstitute.org/ctrp/, accessed 1 Aug. 2019).

The curated datasets (Tables S1–S44) are available on the first author’s website: https://github.com/Jinyu2019/Suppl-data-BBpaper.

### 4.1 The GDSC data

#### Genomic datasets

The GDSC genomic profiles include the gene expression, CNA, mutation, and methylation datasets. These datasets were preprocessed by (i) removing genes with little variation in expression, (ii) extracting both mutations and CNAs for the cancer genes listed in the COSMIC database (https://cancer.sanger.ac.uk/cosmic/download, version 88), and (iii) keeping CpG probes with variation in methylation levels, respectively. In the end, we obtained:

- A gene expression profile *E*_734*×*8046_,
- A DNA methylation profile *D*_734*×*8473_,
- A mutation profile *M*_734*×*636_, and
- A CNA profile *C*_734*×*694_

for 734 cancer cell lines (Tables S1–S4), of which annotations are provided in Table S5.

#### Drug response dataset

The level of drug response was measured in both IC_50_ and AUC in the GDSC database. After removing drugs of which the responses are missing for 80% or more cell lines, we obtained two drug response matrices (Tables S6 and S7) for 734 tumor cell lines and 201 drugs. Table S6 summarizes the common logarithm of the IC_50_ drug response readouts, whereas Table S7 summarizes the AUC readouts.

##### Protein–protein interaction network

A protein–protein interaction (PPI) network on 7,136 proteins (Table S8) was derived by extracting interactions relations between them in the Pathway Commons database (http://www.pathwaycommons.org/, accessed 11 March 2019) [41].

#### Drug groups and drug similarity matrix

The structural information on the 201 drugs include 1,444 1-dimensional, 2-dimensional descriptors and nine types of molecular fingerprints in PubChem [42] (Table S9). The structural information on the drugs were further processed with the PaDEL-Descriptor [43] to construct a drug similarity network, named *S*^201*×*201^ (Table S10).

The 201 drugs were divided into 23 groups (Table S12), each containing drugs targeting the same pathway, according to the information available in the GDSC database.

#### Drug-targeted pathways

We obtained the drug-targeted pathways via the data preprocessing method used in [17]. We downloaded 1,329 canonical pathways and 189 oncogenic signature gene sets in C2:CP and C6 collections in the MSigDB database (http://software.broadinstitute.org/gsea/msigdb/index.jsp, accessed 1 Feb 2019) [44]. We further extracted only those pathways that contained one or more primary targets of the 201 drugs, leading to 71 drug-targeted pathways. All the genes that were not in any extracted pathways formed a single group named “others”. We obtained 72 gene subsets in total (Table S11).

### 4.2 The CCLE data

The CCLE data contain six genomic profiles: gene expression, microRNA expression, protein expression, DNA methylation, mutation and CNA. These six profiles were preprocessed in the same way as we preprocessed the GDSC data. The final CCLE data for 385 cancer cell lines consisted of:

- an expression profile *E*_385*×*17726_ of 17,726 genes,
- an expression profile *O*_385*×*734_ of 734 microRNAs,
- an expression profile *P*_385*×*214_ of 214 proteins,
- a methylation profile *D*_385*×*4507_ of 4,507 DNA methylation sites,
- a mutational profile *M*_385*×*444_ of 444 cancer genes, and
- a CNA profile *C*_385*×*700_ of 700 cancer genes,

which can be found in Tables S13–S18.

The CCLE drug response datasets contain both IC_50_ and AUC readouts for 23 drugs and 385 cell lines (Tables S19–S20). Other CCLE datasets used for assessment include drug-similarity matrix, drug group information, a protein–protein interaction network, drug structural information, cell-line tissue information and gene groups (Tables S21– S26).

### 4.3 The NCI-60 data

The NCI-60 data were downloaded via calling a R program named rcellminer [45]. The NCI-60 genomic profiles were preprocessed in the same way as we preprocessed the GDSC and CCLE data. The final data for 59 cancer cell lines included:

- an expression profile *E*_59*×*18077_ of 18,977 genes,
- an expression profile *O*_59*×*410_ of 410 microRNAs,
- an expression profiles *P*_59*×*162_ of 162 proteins,
- a methylation profiles *D*_59*×*11330_ of 11,330 DNA methylation sites,
- a mutation profiles *M*_59*×*443_, and
- a CNA profile *C*_59*×*394_ of 394 cancer genes,

which are found in Tables S27–S32.

The NCI-60 drug response datasets contain only GI_50_ readouts for 215 drugs and 59 cell lines (Table S33), where GI_50_ is the concentration for 50% of maximal inhibition of cell proliferation. Other CCLE datasets used for assessment include a protein–protein interaction network, drug-similarity matrix, drug structural information, cell-line tissue information and gene group information, drug group information (Tables S34–S39)..

### 4.4 The Cancer Therapeutics Response Portal (CTRP) dataset

The CTRP data (version 2.1) include gene expression profile (Table S40), AUC readouts for 63 drugs in 720 caner cell lines (Table S41), cell line tissue information (Table S42) and drug group information (Table 43) [29].

## 5 Assessment methods

Using the GDSC, CCLE, NCI-60 and CTPR datasets, we assessed the following 13 representative methods developed in the past five years in the nine metrics (Eqn. (16)–(24)):

- CaDRReS (https://github.com/CSB5/CaDRReS),
- CDRscan (https://github.com/summatic/CDRScan),
- cwKBMF (https://research.cs.aalto.fi/pml/),
- DualNets (coded in R by the first author,)
- GENet-Lap and GENet-NLap (available in R),
- KRL (https://github.com/BorgwardtLab),
- MACAU (https://github.com/saezlab),
- MERGE (http://merge.cs.washington.edu/),
- MVLR (https://github.com/suleimank/mvlr),
- pairwiseMKL (https://github.com/aalto-ics-kepaco),
- RWEN-left (resp. right, and both) (https://github.com/kiwtir/RWEN),
- SRMF (https://github.com/linwang1982/SRMF),
- TANDEM (http://ccb.nki.nl/software/tandem),

as well as four classical approaches: BMTMKL (https://github.com/mehmetgonen/bmtmkl), ENet, RF-g and RF-gs, which are available in R.

The auxiliary omics data used by the methods are provided in Table 1. The input data were prepared in the format recommended by the developer for each method.

We used the five-cross validation approach to assess the 17 methods. All the cell lines were randomly partitioned into five equal parts, four of which were used for training and the other part was used for test. The partition was done under the constraint that for each tissue type, the cell lines in the training data were kept in the same proportion as in the test data. The predicted response level to the drugs in all cell lines were then compared against the observed values in each metric. This assessment process was repeated 10 times to obtain a robust evaluation for each dataset. To find out the extend to which the prediction accuracy increases through integrating multiple genomic profiles other than gene expression, we ran ENet and BMTMKL in two settings: (i) gene expression profile only and (ii) multiple profiles including gene expression, mutational, CNA, and DNA methylation profiles. The multi-view version of them are denoted M-Enet and M-BMTMKL in Table 2, respectively.

**Table 2.**
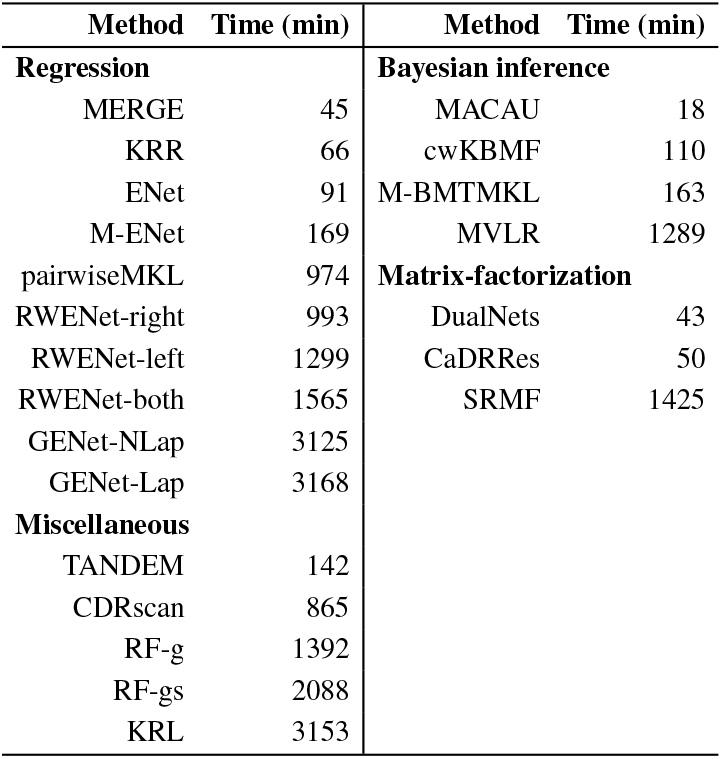
The run time of the 17 methods on the GDSC data. The CDRscan was assessed on a GPU machine, whereas the rest were assessed on a PC cluster with 6 cores and 45GB RAM. The time taken by parameter selection was included into the run time for each method. GNet-Lap and GNet-NLap are two variants of GNet; M-ENet is a variant of ENet that integrated multiple genomic profiles of cell lines; RWENet-left, RWENet-right and RWENet-both are three variants of RWENet;

A method that has low accuracy in general may perform extremely well when applied to the cell lines from a tissue or the drugs within group. To find out whether this is true or not, we tested the methods on the datasets containing cell lines derived from only one tissue. We assessed the methods on the following tissue specific datasets:

- GDSC: breast, central nervous, haematopoietic and lymphoid, lung and skin;
- CCLE: breast, haematopoietic and lymphoid, lung, ovary and skin;
- NCI-60: melanoma and lung.

We also tested whether a method trained in a dataset still performs well on another drug response dataset. To this purpose, we trained the methods on the GDSC data and test them on the CCLE and CTRP datasets, respectively.

Lastly, the one-sided paired Wilcoxon Sign-Rank test ([46, page 338]) was used to determine whether or not a method was significantly better than another in each metric. The heat map of the -log_10_(*p*-values) of the tests is summarized for pairwise method comparison.

The CDRscan method was validated on a PC cluster empowered with a GPU (Nvidia Tesla V100-SXM2). All other methods were validated on a PC cluster with 6 cores and 45GB RAM. The computational time of different methods are summarized in Table 2.

## 6 Results and discussions

The test results on the GDSC data are summarized in Figures 1–3 (and Supplementary file 1: Figures S1–S3) for the IC_50_ drug response readouts and in Supplementary file 2: Figures S9–S11 for the AUC readouts. The results on the CCLE data are summarized in Supplementary file 3: Figures S17–S19 for the IC_50_ readouts and in Supplementary file 4: Figures S25– S27 for the AUC readouts. The results on the NCI-60 data are summarized in Supplementary file 5: Figures S33–S35 for the GI_50_ readouts.

**Fig. 1.**
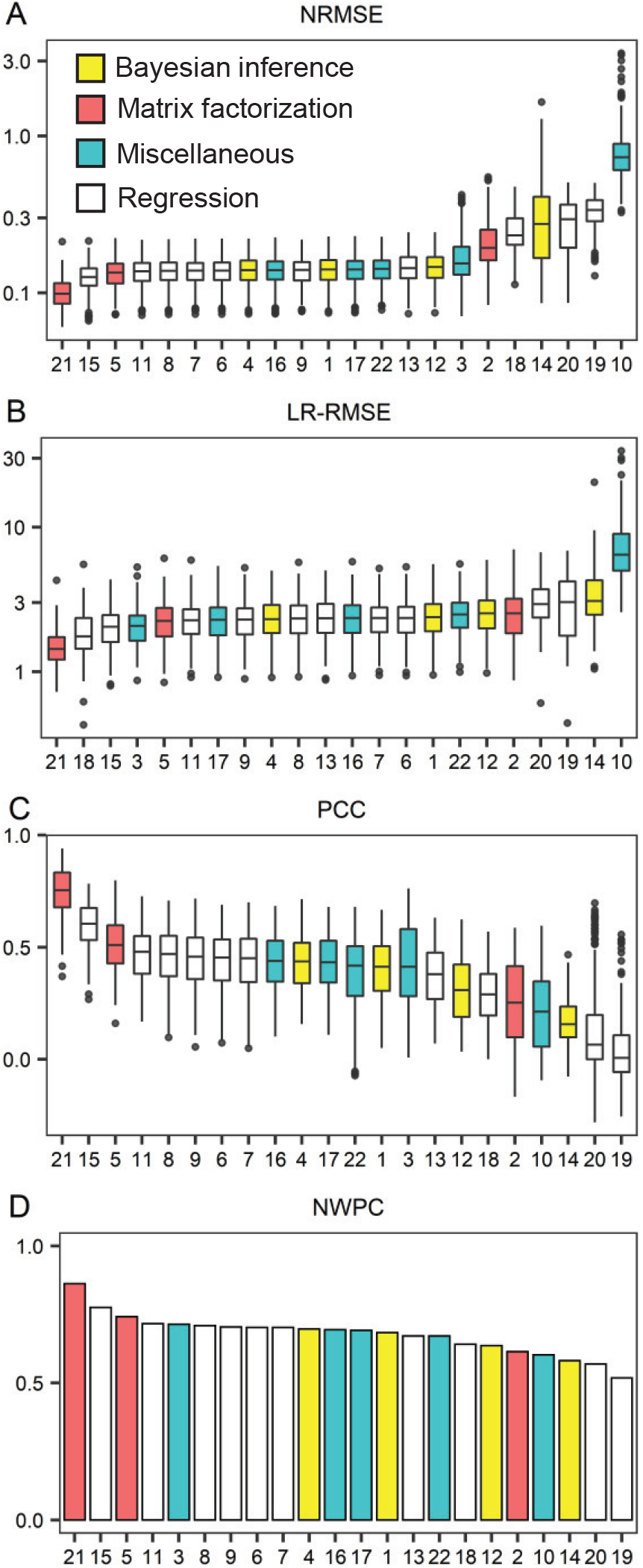
The performance of the 17 methods on the GDSC IC_50_ dataset. Box plots were drawn for the normalized root mean square error (NRMSE) (A), the left-right (LR)-RMSE (B), the Pearson correlation coefficient (PCC) (C) and the normalized weighted probabilistic c-index (NWPC) (D), where the methods were listed from the best (leftmost) to the worst. Each box plot was drawn on the basis of the scores of 10 repeated cross validation tests for the following methods: BMTMKL (1), CaDRReS (2), CDRscan (3), cwKBMF (4), DualNets (5), ENet (6), M-ENet (7), GENet-Lap (8), GENet-NLap (9), KRL (10), KRR (11), MACAU (12), MERGE (13), MVLR (14), pairwiseMKL (15), RF-g (16), RF-gs (17), RWENet-both (18), RWENet-left (19), RWENet-right (20), SRMF (21), TANDEM (22).

The performance of the methods on the GDSC tissue-specific data are summarized in Supplementary file 1: Figures S4–S8 and Supplementary file 2: Figures S12–S16 for the IC_50_ and AUC readouts, respectively. The performance of the methods on the CCLE tissue-specific data are summarized in Supplementary file 3: Figures S20–S24 and Supplementary file 4: Figures S28–S32 for the IC_50_ and AUC readouts, respectively. The performance of the methods on the NCI-60 tissue-specific data with the GI_50_ readouts are summarized in Supplementary file 5: Figures S36–S37.

The performance of the methods when trained on the GDSC data and tested on the CCLE data are summarized in Supplementary file 6: Figures S38–S44 for the IC_50_ readouts and Supplementary file 7: Figures S45–S51 for the AUC readouts. The performance of the methods when trained on the GDSC AUC data and tested on the CTRP AUC data are summarized in Supplementary file 8: Figures S52–S58.

### 6.1 Pros and cons of the methods

Different methods have quite different time complexity (Table 2). The fastest methods such as MACAU, MERGE and DualNets took less than an hour on average, including the parameter selection time. The slowest methods include GELNet, KRL, SRMF and RWENet and took over 20 hours.

Our studies suggest that the assessed methods can be divided into three groups in terms of prediction accuracy: good methods (DualNets (Method 5 in the figures), KRR (11), pairwiseMKL (15) and SRMF (21)), unstable methods (BMTMKL, CDRscan, cwKDMF, ENet and M-ENet, GENet-Lap and GENet-NLap, MACAU, MERGE, RF and TANDEM), and poor methods (CaDRReS (2), KRL(10), MVLR (14) and RWENet (18)) (Figure 1, Figures S1, S9, S17, S25 and S33). The four good methods outperformed constantly the other methods almost on every dataset in each metric, whereas the poor methods had the lowest accuracy almost on every test.

The accuracy of each unstable method fluctuated dramatically across datasets and evaluation metrics. For instance, on the GDSC dataset, the deep learning method CDRscan had the forth or fifth best performance if LR-RMSE, L-RMSE, R-RMSE or NDCG was used for evaluation; in contrast, it had the 7-th worst performance if each of the RMSE and NRMSE was used.

When trained on the GDSC data and tested on the CCLE or CTRP data, the four good methods did not retain a clear edge in performance (Figures S38–S58).

Our results show that the matrix-factorization-based approach, represented by SRMF, performed better than the regression models (with or without kernel extension), whereas the regression models performed better than the different Bayesian inference methods. This has the following implications.

First, the drug response is likely a complicated non-linear function of the cell line profiles and the drug profiles. When predicting the response to a known drug in a new cell line, as one of the best methods, DualNets assumes that the drug response level is linear to the drug profile but non-linear to the cell line profile (Eqn. (13)–(14)), whereas ENet and GENet and other regression methods consider the drug response function to be linear to the cell line profiles (see Eqn. (1)).

SRMF, another best method, assumes that the drug response matrix is a product of the low-rank proximity of the cell line similarity matrix and the low-rank proximity of the drug similarity matrix, which are both non-linear to cell line profiles and drug profiles, respectively.

Second, Bayesian inference is usually powerful for solving machine learning problems. Possible causes for the ineffectiveness of the assessed Bayesian inference methods may include the distribution of drug responses may not be as simple as a normal distribution, the parameter space in each method may be too large to compute an optimal set of parameter values and the parameter space is so large that the methods may overfit the training dataset. Our Shapiro-Wilk normality test indicates that most drug response dataset do not follow a normal distribution.

### 6.2 Trade-off between feature selection and prediction accuracy

Feature selection is an important function for methods that are developed for drug response prediction. It allows to identify valuable biomarkers and thus enables the follow-up biological studies into drugs. The methods with this function include ENet and its generalization GEnet, MERGE, MVLR, RF, RWEN and TANDEM (Table 1). ENet and GENet are the best among these methods. They were fast (Table 2) and had slightly lower accuracy than the most accurate methods DualNets, KRR, pairwiseMKL and SRMF in our test (Figure 1, Supplementary file 1: Figure S2, Supplementary file 2: Figure S9, Supplementary file 3: Figure S17, Supplementary file 4: Figure S25 and Supplementary file 5: Figure S33).

### 6.3 Protein–protein interaction networks are informative

GENet is the only model that uses explicitly cellular protein–protein interaction networks for drug response prediction. GENet-Lap and GENet-NLap were the two versions developed from ENet through incorporating the (normalized) Laplacian matrix of a protein–protein interaction network into the last regularization term in Eqn. (1). Interestingly, the two methods outperformed ENet for seven out of the nine metrics (Figure 1–2, see also Figure S1–S2). This fact may indicate that drug response prediction can benefit from the integration of both protein–protein interaction networks and gene expression profiles with the drug response data.

**Fig. 2.**
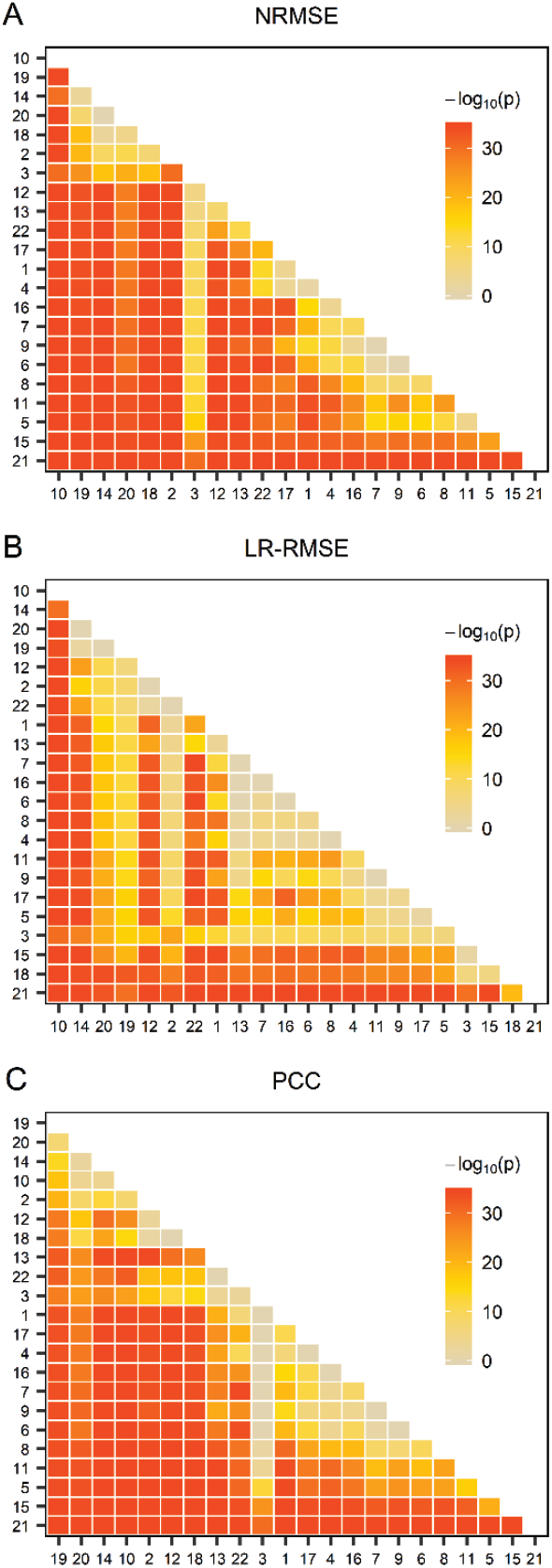
The comparison of the performance of the methods on the GDSC IC_50_ dataset. The one-sided paired Wilcoxon Sign-Rank test was conducted on the data appearing in Figure 1 (A–C). The heatmap was drawn from the negative common logarithm of the p-value for the hypothesis that a method performs better than another. BMTMKL (1), CaDRReS (2), CDRscan (3), cwKBMF (4), DualNets (5), ENet (6), M-ENet (7), GENet-Lap (8), GENet-NLap (9), KRL (10), KRR (11), MACAU (12), MERGE (13), MVLR (14), pairwiseMKL (15), RF-g (16), RF-gs (17), RWENet-both (18), RWENet-left (19), RWENet-right (20), SRMF (21), TANDEM (22).

### 6.4 Mutation, CNV and methylation profiles of cell lines and drug grouping add little to drug response prediction

We assessed ENet and BMTMKL in two settings: (a) only gene expression profile was used and (b) the gene expression, mutation, CNA and methylation profiles were integrated. We did not observe clear improvement in accuracy when multiple genomic profiles of cell lines were used (Figure 1–2 and Figure S1–S2).

MVLR is the only method that used the information on drug-targeted pathways. It builds a linear regression model for each group of drugs that have the same target pathway. Its predictive performance varied from group to group (Figure 3). Both Figures 1 and 3 suggest that MVLR performed better only on four out of 23 drug groups than without group information.

**Fig. 3.**
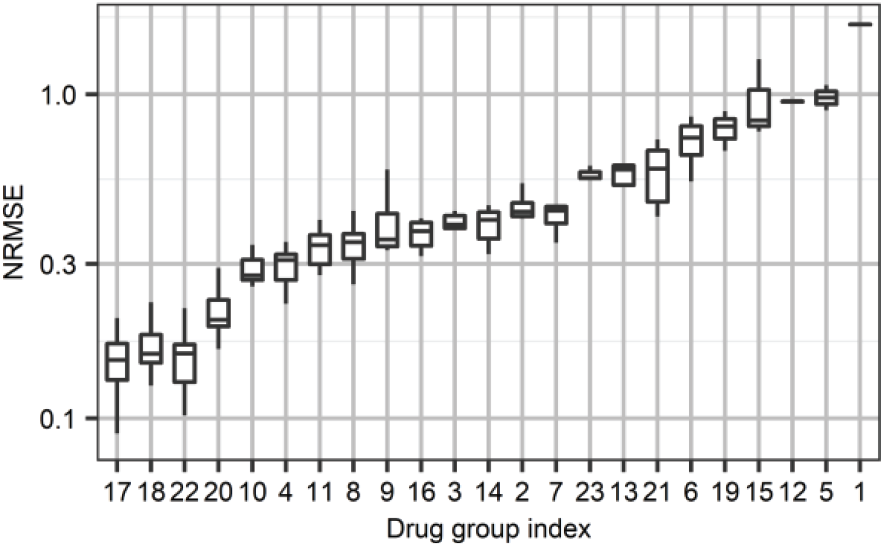
The distribution of the NRMSE scores of the MVLR method when trained and tested within drug groups. All drugs were divided into 23 groups depending on their signaling pathway target. For each group, MVLR learnt a model and made prediction. The drug groups are indexed alphabetically: ABL signaling (1), Apoptosis regulation (2), Cell cycle (3), Chromatin histone acetylation (4), Chromatin histone methylation (5), Chromatin other (6), Cytoskeleton (7), DNA replication (8), EGFR signaling (9), ERK MAPK signaling (10), Genome integrity (11), Hormone-related (12), IGFR signaling (13), JNK and p38 signaling (14), Metabolism (15), Mitosis(16), other drugs (17), Kinases (18), P53 pathway (19), PI3K/MTOR signaling (20), Protein stability and degradation (21), RTK signaling (22), WNT signaling (23).

### 6.5 The performance of a method varies a lot under different measurements

We used nine performance metrics in this study. RMSE and NRMSE measure the performance of a method on all the data points, whereas L-RMSE, R-RMSE and LR-RMSE focus on the performance at the head, tail or both sides of the drug response data distribution. SCC, NDCG and NWPC consider all the data points but measure the rank coherency between the predicted and observed values.

Our study suggests that the performance can be very sensitive to the metric selected for assessment for unstable methods. An unstable method could significantly outperform another method for a metric, but perform poorly when comparing other metrics. For instance, cwKBMF significantly outperformed RF-gs, with *P*-value 10^*−*3^, 10^*−*4^ and 10^*−*6^ in the RMSE, NRMSE and SCC, respectively (Figures 2 and S2); in contrast, RF-gs significantly outperformed cwKBMF, with *P*-value 10^*−*10^, 10^*−*9^ and 10^*−*2^ in L-RMSE, LR-RMSE and NDCG; and there was no significant difference between cwkBMF and RF-gs for R-RMSE and PCC.

Taken together, the two facts suggest that three or four different metrics such as NRMSE, PCC and NDCG may be used for a robust assessment of a method for drug response prediction.

## 7 Conclusion

We assessed four classical methods and 13 representative computational methods that have recently been developed for drug response prediction using public cell line–drug response data and nine metrics. This work provides lessons worth considering ahead of future research.

First, the study reconfirms that the gene expression profiles of cell lines provide crucial information for predicating the responses to a drug in cell lines [10, 11]. How to generate and use the time point series expression profiles of cell lines is an important issue for drug response prediction in future preclinical and clinical research.

It is commonly accepted that omics data other than gene expression profiles of cell lines often add slight improvements for drug response prediction [9, 10]. However, the excellent performance of DualNets suggests that smart integration of other omics data, particularly drug structure data and protein interaction networks, with gene expression profiles should not be ignored in the study of drug response prediction.

Second, SRMF learns drug response functions from the similarity matrices of cell lines and drugs by an iterative approach. This may suggest deep neural network and other AI approaches are likely to be powerful for drug response prediction. Given that the deep learning method CDRscan did not perform better than the top four methods mentioned above, it is non-trivial to apply deep learning approach to drug response prediction. Another research question is how to improve the performance of Bayesian inference based methods for drug response prediction.

Third, interpretability and reproducibility are two important factors for drug response models in translation medicine research [11, 47, 48]. For pan-cancer drug response data, different metrics give different pictures of the evaluated methods’ performance. Linear regression models and their generalizations are good at feature selection and thus has advantage in interpretability. Unfortunately, these methods have low accuracy. Therefore, how to balance predication accuracy and interpretability is an important issue. Aben et al. has made a promising progress toward this goal in [15]. In future, computational biologists may focus on cancer-specific drug response data to design robust prediction methods with interpretable features [49].

### Key Points

- The computational methods for drug response prediction can be grouped into three classes: (a) linear regression models (with or without kernel extension), (b) Bayesian inference methods and (c) matrix factorization via nonlinear optimization methods. In our test, the matrix factorization approach outperformed the linear regression models, whereas linear regression models outperformed different Bayesian inference methods. It remains unclear how to improve the performance of Bayesian inference methods for drug response prediction.
- SRMF with the best performance learns drug response functions from the similarity matrices of cell lines and drugs by an iterative approach, revealing that the drug response functions of cell lines are non-linear and complicated. This may suggest deep neural network and other AI approaches are likely to be able to achieve high accuracy for drug response prediction. On the other hand, our analyses also suggest that it is far from trivial to design a deep learning method that surpasses the top regression or matrix factorization-based methods for drug response prediction.
- The mutation, CNV and methylation profiles of cell lines and drug grouping seem to add little to drug response prediction when they are used with the existing methods. It is worth studying how to use wisely these omics data.
- The performance of a method is sensitive to the choice of measurements. One method could significantly outperform another method for one metric, but performs poorly when comparing other metrics. Three or more metrics that are defined for assessing different aspects of drug response predication, such as the combination of NRMSE, PCC and NDCG, should be used for evaluation of a new method.
- Four large datasets of cell line genomic profiles and drug profiles were curated for benchmarking for future study into drug response prediction.

## Supplementary documents

Supplementary tables are available at https://github.com/Jinyu2019/Suppl-data-BBpaper

## Acknowledgements

The authors would like to thank Rudi Alberts for help in testing CDRscan and Lin Wang for useful discussion on matrix factorization based methods. They also thank anonymous reviewers for their useful comments on the first version of the manuscript, which improve significantly our work.

