## Supplementary File 1 for "A Survey and Systematic Assessment of Computational Methods for Drug Response Prediction"

**Figure S1**. The performance of the methods on the GDSC IC50 drug response dataset under different metrics

**Figure S2**. The performance comparison of the methods on the GDSC IC50 dataset.

**Figure S3.** The performance of the methods when trained and tested within 23 drug groups on the GDSC IC50 drug response dataset under different metrics.

**Figure S4**. The performance of the methods on breast cell lines appearing in the GDSC dataset with IC50 drug response readouts.

**Figure S5**. The performance of the methods on cell lines of central nervous system appearing in the GDSC dataset with IC50 drug response readouts.

**Figure S6**. The performance of the methods on the cell lines of Haematopoietic and lymphoid tissue appearing in the GDSC dataset with IC50 drug response readouts.

**Figure S7**. The performance of the methods on the lung cell lines appearing in the GDSC dataset with IC50 drug response readouts.

**Figure S8**. The performance of the methods on the skin cell lines appearing in the GDSC dataset with IC50 drug response readouts

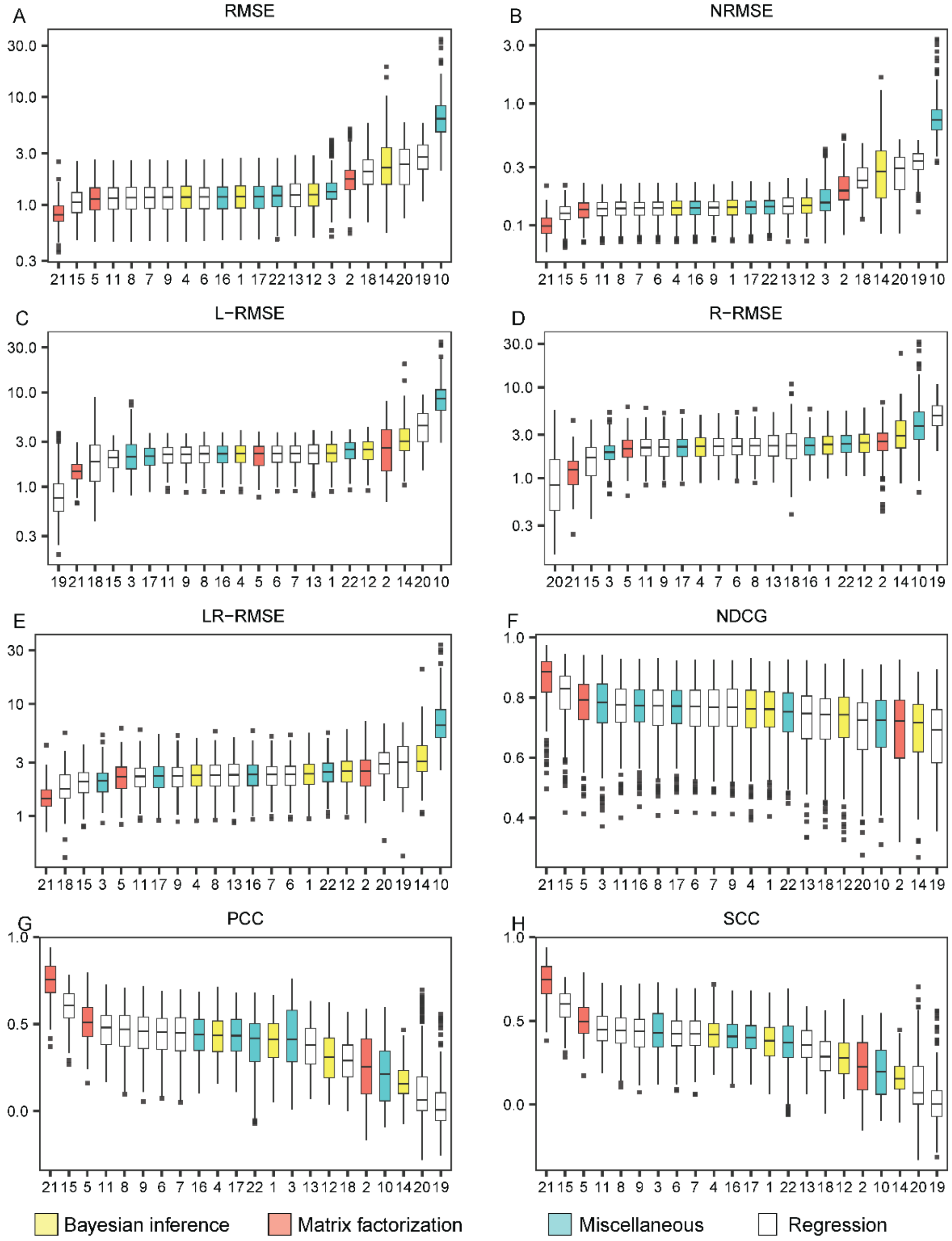


**Figure S1. The performance of the methods on the GDSC IC50 drug response dataset under different metrics.** Box plots were drawn for the root mean square error (RMSE) (A), the normalized RMSE (B), the left-RMSE (C), the right-RMSE (D), the left-right-RMSE (E), the normalized discounted cumulative gain (NDCG), the Pearson correlation coefficient (PCC) (G) and the Spearman correlation coefficient (SCC) (H), in which the methods are listed from the best (leftmost) to the worst. Each figure was made on the basis of the scores of 10 repeated cross validation tests for each method. The methods are indexed alphabetically: BMTMKL (1), CaDRReS (2), CDRscan (3), cwKBMF (4), DualNets (5), ENet (6), M-ENet (7), GENet-Lap (8), GENet-NLap (9), KRL (10), KRR (11), MACAU (12), MERGE (13), MVLR (14), pairwiseMKL (15), RF-g (16), RF-gs (17), RWENet-both (18), RWENet-left (19), RWENet-right (20), SRMF (21), TANDEM (22).

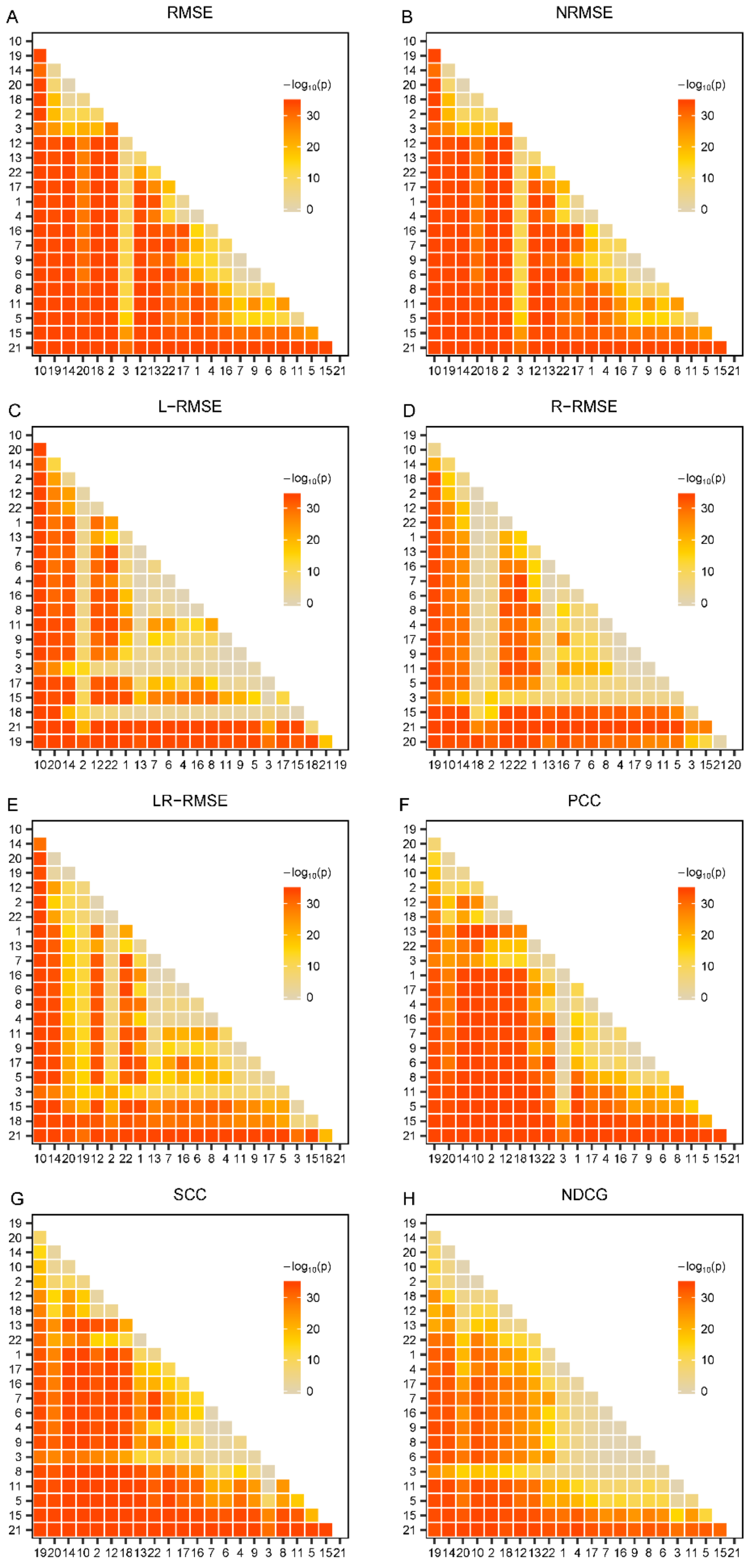


**Figure S2. Performance comparison of the methods on the GDSC IC50 dataset.**
(Cont’ed on next page)

(Cont'ed) The one-sided paired Wilcoxon Sign-Rank test was conducted on the basis of the data shown in Figure S1. The heatmap was drawn from the negative common logarithm of the p-value for the hypothesis that a method performs better than another. BMTMKL (1), CaDRReS (2), CDRscan (3), cwKBMF (4), DualNets (5), ENet (6), M-ENet (7), GENet-Lap (8), GENet-NLap (9), KRL (10), KRR (11), MACAU (12), MERGE (13), MVLR (14), pairwiseMKL (15), RF-g (16), RF-gs (17), RWENet-both (18), RWENet-left (19), RWENet-right (20), SRMF (21), TANDEM (22).

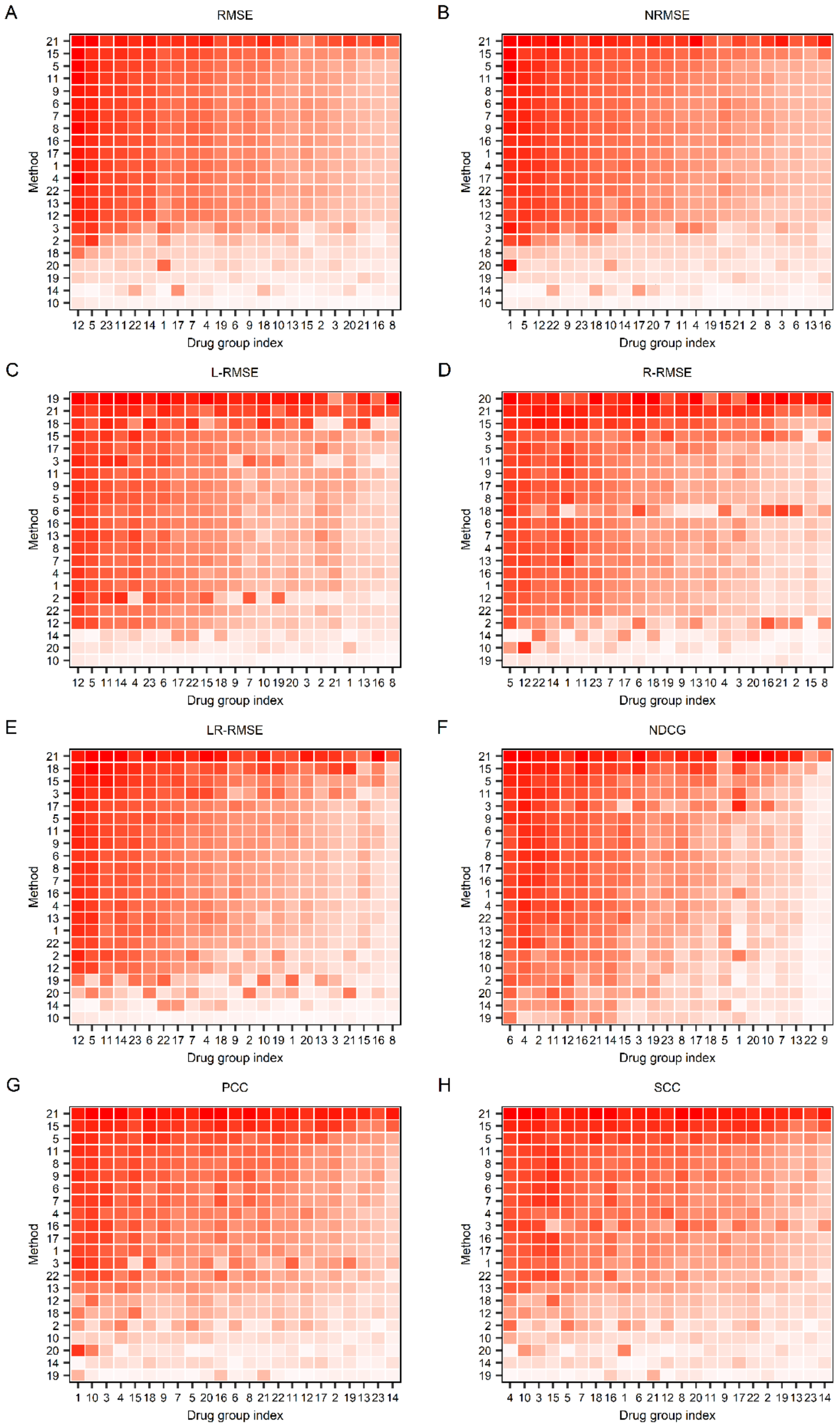


**Figure S3. Comparison of the performance of the methods when trained and tested within 23 drug groups on the GDSC IC50 drug response dataset under different metrics.** (Cont'd on next page)

**(Cont'd)**

The root mean square error (RMSE) (A), the normalized RMSE (B), the left-RMSE (C), the right-RMSE (D), the left-right-RMSE (E), the normalized discounted cumulative gain (NDCG) (F), the Pearson correlation coefficient (PCC) (G), and Spearman correlation coefficient (SCC) (H) metrics. Each row and column represent one method and one drug group, respectively. For the method (row) i and drug group (column) j, the median of the measures for drugs in the group j computed by the method i is used to color the corresponding cell, where dark red represents good performance, whereas light red represents bad performance. The methods are indexed alphabetically: BMTMKL (1), CaDRReS (2), CDRscan (3), cwKBMF (4), DualNets (5), ENet (6), M-ENet (7), GENet-Lap (8), GENet-NLap (9), KRL (10), KRR (11), MACAU (12), MERGE (13), MVLR (14), pairwiseMKL(15), RF-g (16), RF-gs (17), RWENet-both (18), RWENet-left (19), RWENet-right (20), SRMF (21), TANDEM (22). Drug groups are also index alphabetically: ABL signaling (1), Apoptosis regulation(2), Cell cycle(3), Chromatin histone acetylation(4), Chromatin histone methylation(5), Chromatin other(6), Cytoskeleton(7), DNA replication(8), EGFR signaling (9), ERK MAPK signaling(10), Genome integrity(11), Hormone-related(12), IGFR signaling(13), JNK and p38 signaling(14), Metabolism(15), Mitosis(16), Other(17), Kinases(18), p53 pathway(19), PI3K/MTOR signaling(20), Protein stability and degradation(21), RTK signaling(22), WNT signaling (23).

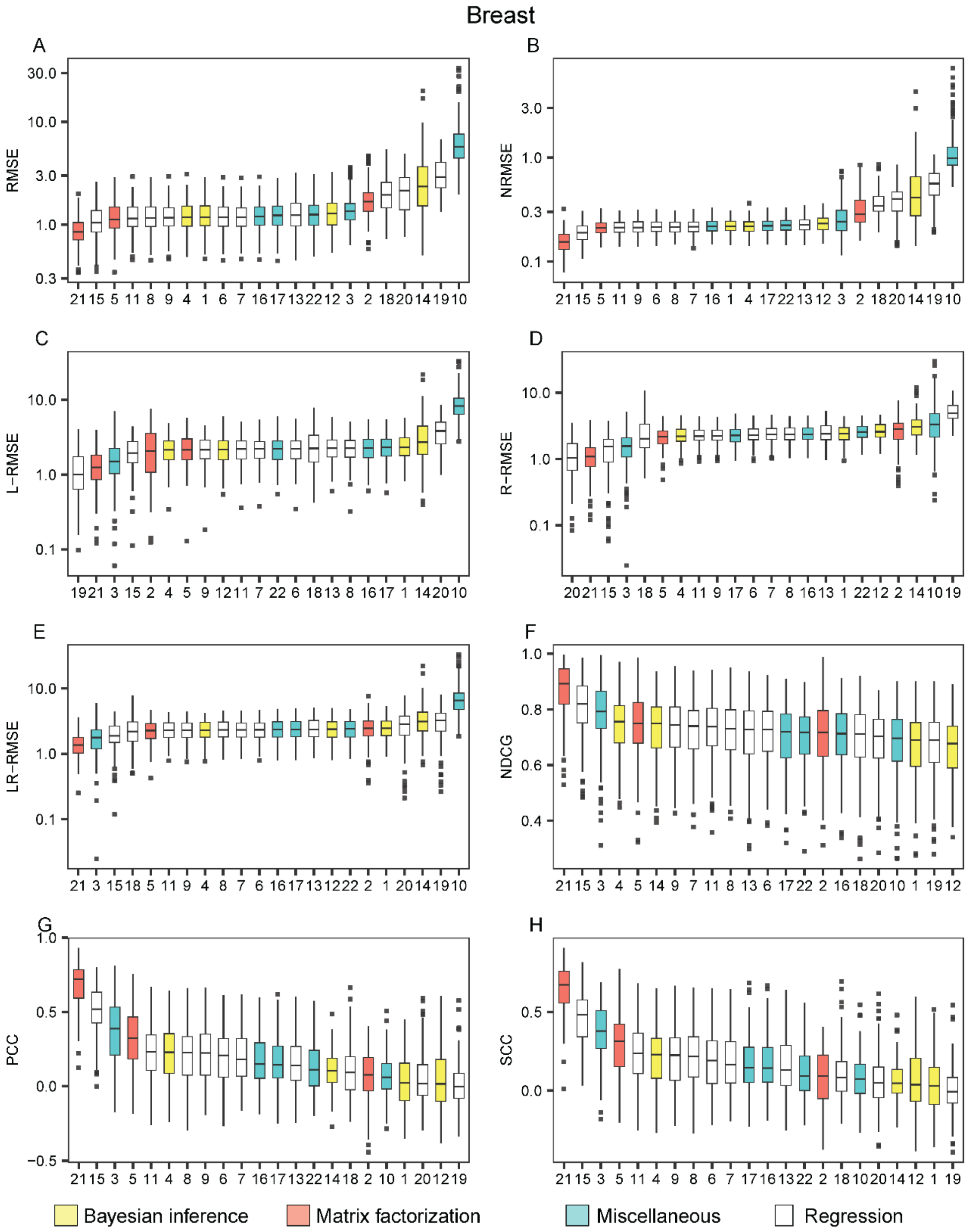


**Figure S4. The performance of the methods on breast cell lines appearing in the GDSC dataset with IC50 drug response readouts**. The methods are listed from the best (leftmost) to the worst in each metric: the root mean square error (RMSE) (A), the normalized RMSE (B), the left-RMSE (C), the right-RMSE (D), the left-right-RMSE (E), the normalized discounted cumulative gain (NDCG), the Pearson correlation coefficient (PCC) (G) and the Spearman correlation coefficient (SCC) (H). The methods are indexed alphabetically: BMTMKL (1), CaDRReS (2), CDRscan (3), cwKBMF (4), DualNets (5), ENet (6), M-ENet (7), GENet-Lap (8), GENet-NLap (9), KRL (10), KRR (11), MACAU (12), MERGE (13), MVLR (14), pairwiseMKL (15), RF-g (16), RF-gs (17), RWENet-both (18), RWENet-left (19), RWENet-right (20), SRMF (21), TANDEM (22).

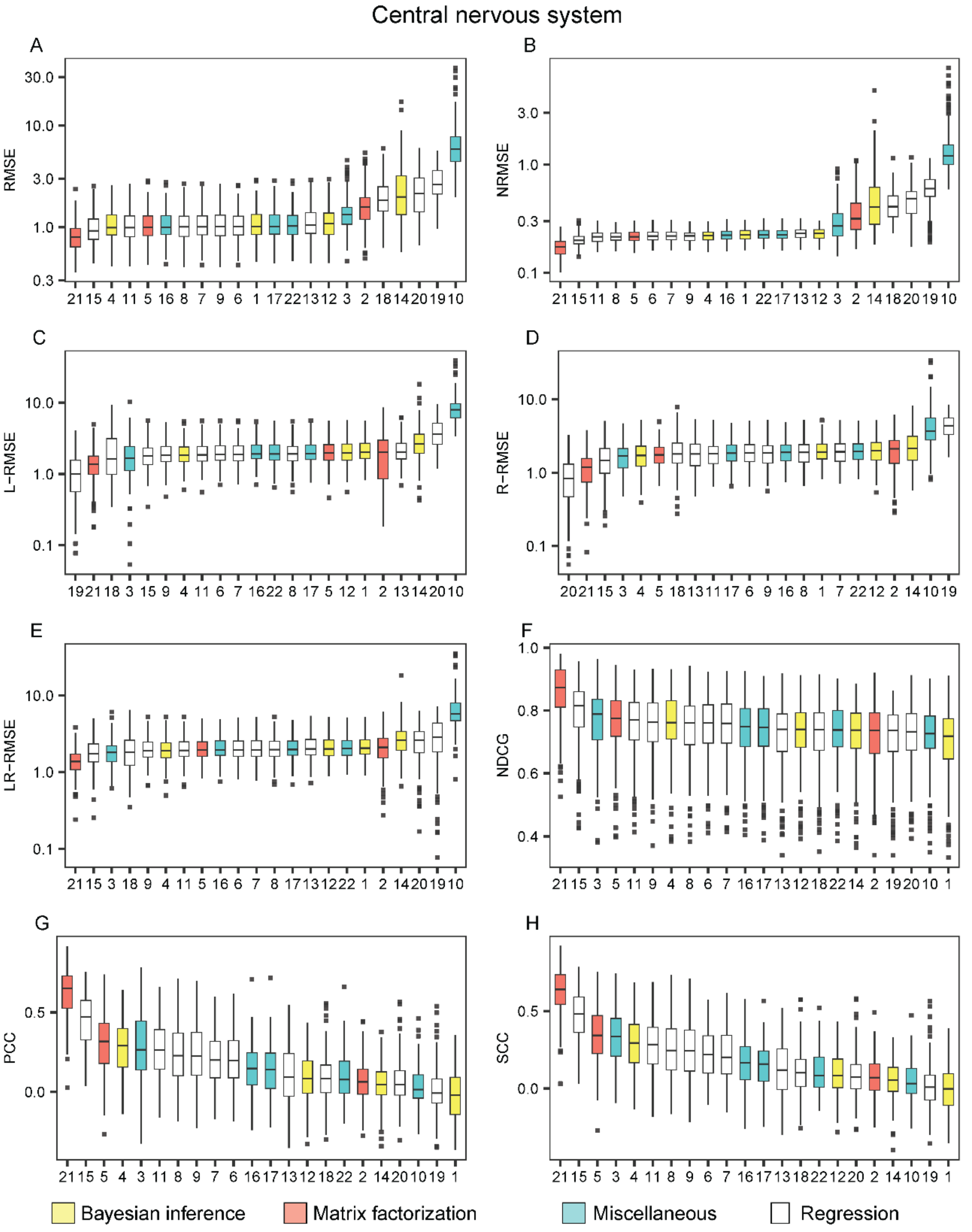


**Figure S5. The performance of the methods on cell lines of central nervous system appearing in the GDSC dataset with IC50 drug response readouts**. The methods are listed from the best (leftmost) to the worst in each metric: the root mean square error (RMSE) (A), the normalized RMSE (B), the left-RMSE (C), the right-RMSE (D), the left-right-RMSE (E), the normalized discounted cumulative gain (NDCG), the Pearson correlation coefficient (PCC) (G) and the Spearman correlation coefficient (SCC) (H). The methods are indexed alphabetically: BMTMKL (1), CaDRReS (2), CDRscan (3), cwKBMF (4), DualNets (5), ENet (6), M-ENet (7), GENet-Lap (8), GENet-NLap (9), KRL (10), KRR (11), MACAU (12), MERGE (13), MVLR (14), pairwiseMKL (15), RF-g (16), RF-gs (17), RWENet-both (18), RWENet-left (19), RWENet-right (20), SRMF (21), TANDEM (22).

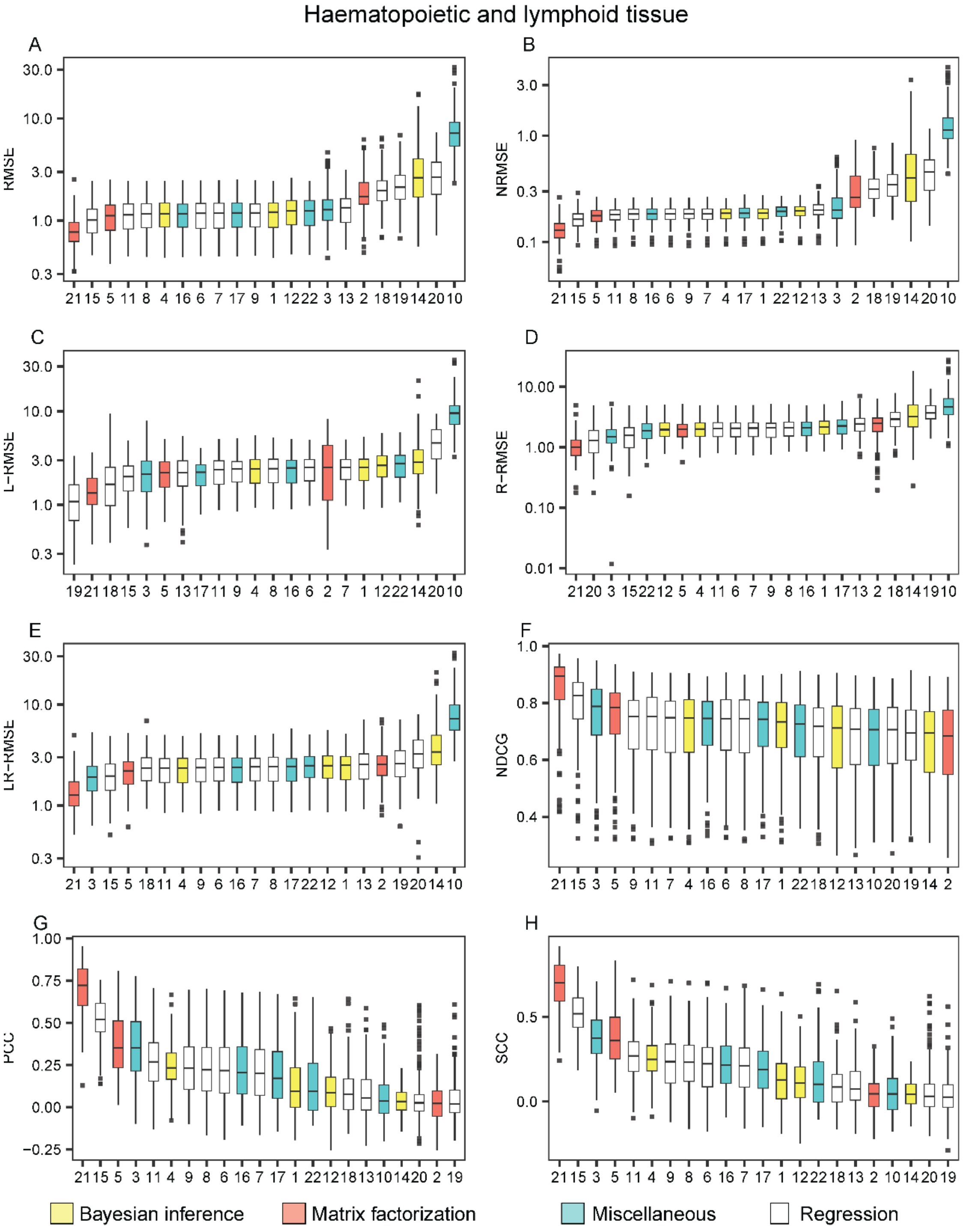


**Figure S6. The performance of the methods on the cell lines of Haematopoietic and lymphoid tissue appearing in the GDSC dataset with IC50 drug response readouts**. The methods are listed from the best (leftmost) to the worst in each metric: the root mean square error (RMSE) (A), the normalized RMSE (B), the left-RMSE (C), the right-RMSE (D), the left-right-RMSE (E), the normalized discounted cumulative gain (NDCG), the Pearson correlation coefficient (PCC) (G) and the Spearman correlation coefficient (SCC) (H). The methods are indexed alphabetically: BMTMKL (1), CaDRReS (2), CDRscan (3), cwKBMF (4), DualNets (5), ENet (6), M-ENet (7), GENet-Lap (8), GENet-NLap (9), KRL (10), KRR (11), MACAU (12), MERGE (13), MVLR (14), pairwiseMKL (15), RF-g (16), RF-gs (17), RWENet-both (18), RWENet-left (19), RWENet-right (20), SRMF (21), TANDEM (22).

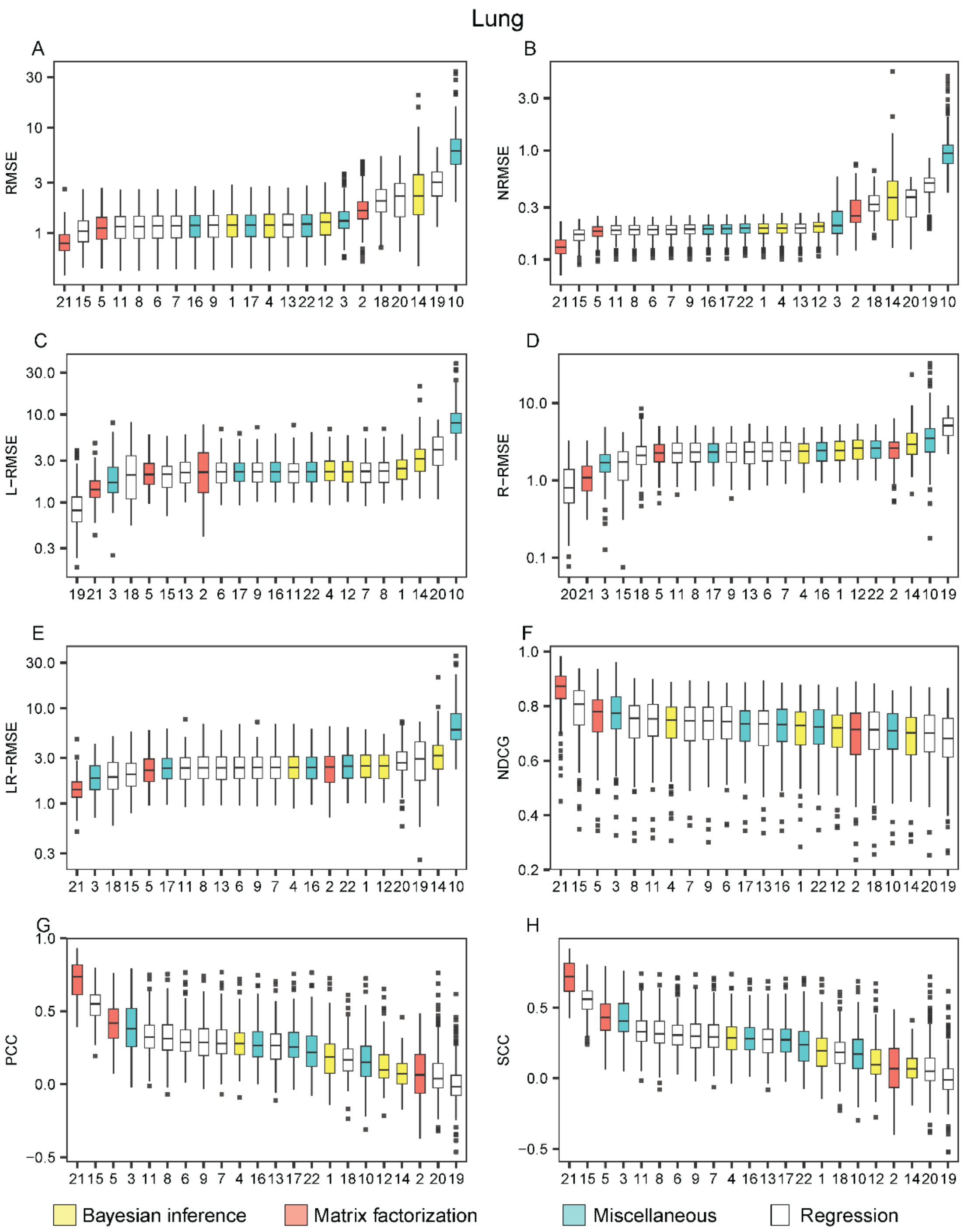


**Figure S7. The performance of the methods on the lung cell lines appearing in the GDSC dataset with IC50 drug response readouts**. The methods are listed from the best (leftmost) to the worst in each metric: the root mean square error (RMSE) (A), the normalized RMSE (B), the left-RMSE (C), the right-RMSE (D), the left-right-RMSE (E), the normalized discounted cumulative gain (NDCG), the Pearson correlation coefficient (PCC) (G) and the Spearman correlation coefficient (SCC) (H). The methods are indexed alphabetically: BMTMKL (1), CaDRReS (2), CDRscan (3), cwKBMF (4), DualNets (5), ENet (6), M-ENet (7), GENet-Lap (8), GENet-NLap (9), KRL (10), KRR (11), MACAU (12), MERGE (13), MVLR (14), pairwiseMKL (15), RF-g (16), RF-gs (17), RWENet-both (18), RWENet-left (19), RWENet-right (20), SRMF (21), TANDEM (22).

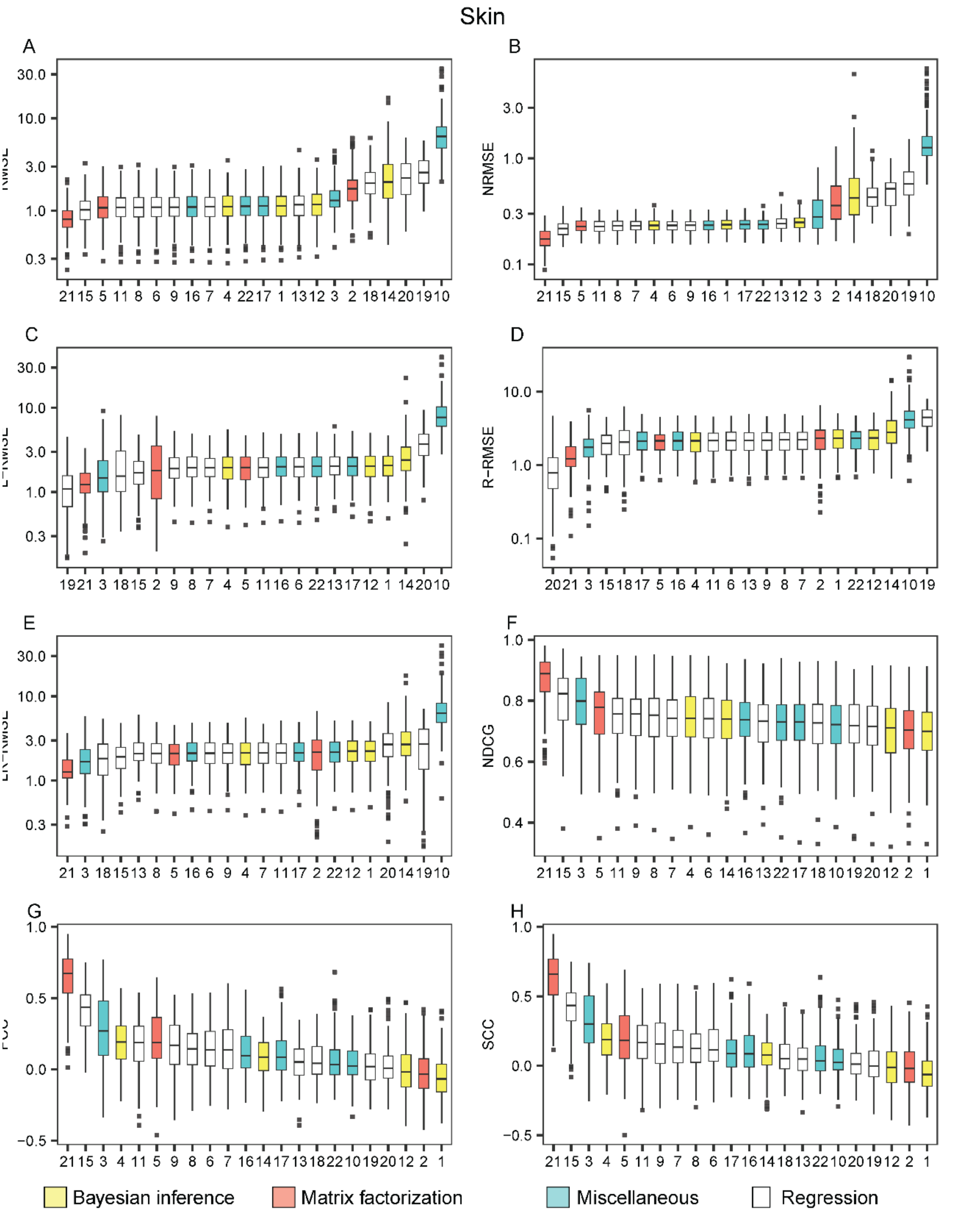


**Figure S8. The performance of the methods on the skin cell lines appearing in the GDSC dataset with IC50 drug response readouts**. The methods are listed from the best (leftmost) to the worst in each metric: the root mean square error (RMSE) (A), the normalized RMSE (B), the left-RMSE (C), the right-RMSE (D), the left-right-RMSE (E), the normalized discounted cumulative gain (NDCG), the Pearson correlation coefficient (PCC) (G) and the Spearman correlation coefficient (SCC) (H). The methods are indexed alphabetically: BMTMKL (1), CaDRReS (2), CDRscan (3), cwKBMF (4), DualNets (5), ENet (6), M-ENet (7), GENet-Lap (8), GENet-NLap (9), KRL (10), KRR (11), MACAU (12), MERGE (13), MVLR (14), pairwiseMKL (15), RF-g (16), RF-gs (17), RWENet-both (18), RWENet-left (19), RWENet-right (20), SRMF (21), TANDEM (22).
