## Supplementary File 4 for "A Survey and Systematic Assessment of Computational Methods for Drug Response Prediction"

**Figure S25**. The performance of the methods on the CCLE AUC drug response dataset under different metrics

**Figure S26**. Performance comparison of the methods on the CCLE AUC dataset.

**Figure S27.** The performance of the methods when trained and tested within 17 drug groups on the CCLE AUC drug response dataset under different metrics.

**Figure S28**. The performance of the methods on the breast cell lines appearing in the CCLE dataset with AUC drug response readouts.

**Figure S29**. The performance of the methods on the cell lines in the haematopoietic and lymphoid tissue appearing in the CCLE dataset with AUC drug response readouts.

**Figure S30**. The performance of the methods on the lung cell lines appearing in the CCLE dataset with AUC drug response readouts.

**Figure S31**. The performance of the methods on the ovary cell lines appearing in the CCLE dataset with AUC drug response readouts.

**Figure S32**. The performance of the methods on the skin cell lines appearing in the CCLE dataset with AUC drug response readouts

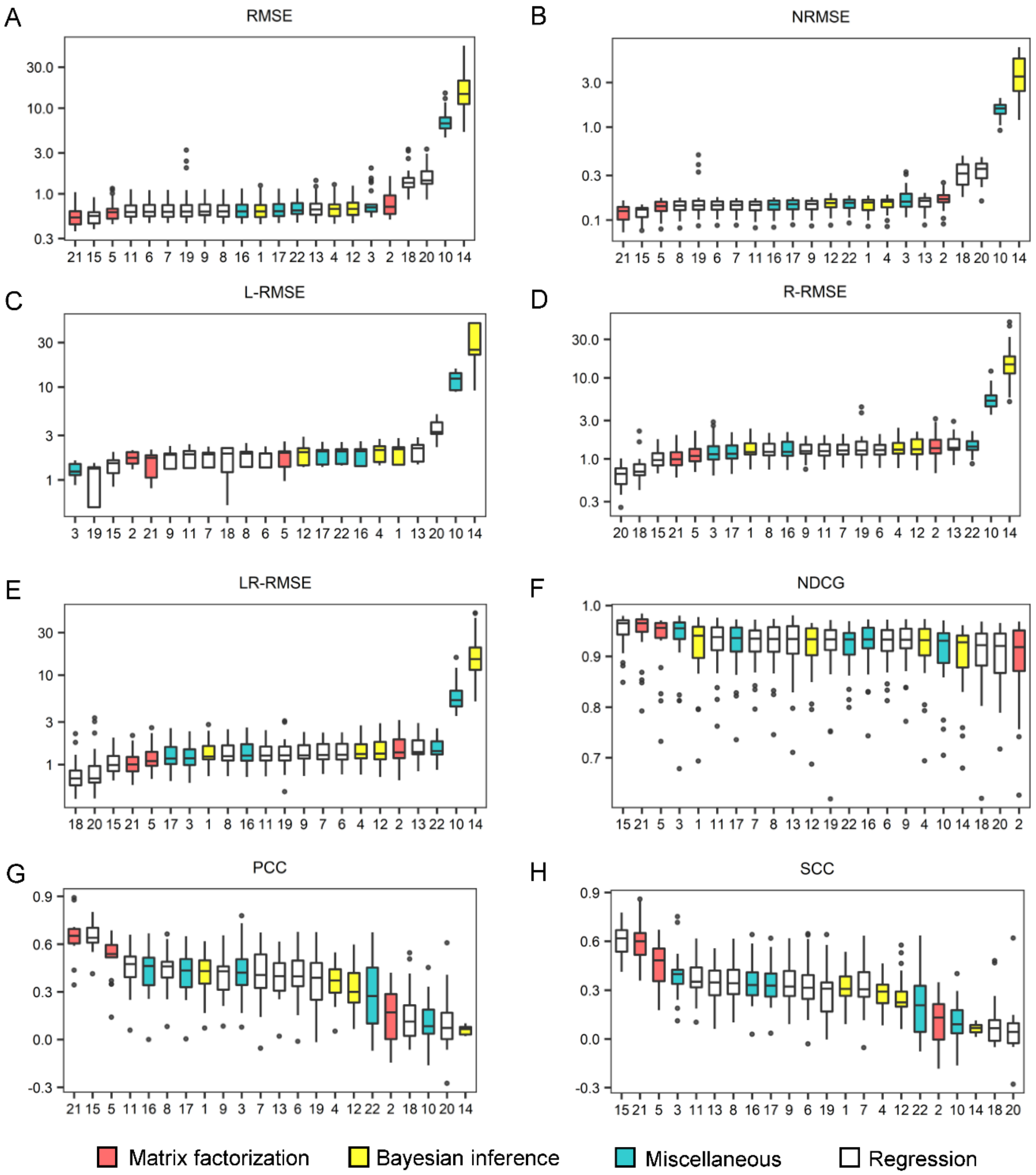


**Figure S25. Comparison of the performance of the methods on the CCLE AUC drug response dataset under different metrics.** Box plots were drawn for the root mean square error (RMSE) (A), the normalized RMSE (B), the left-RMSE (C), the right-RMSE (D), the left-right-RMSE (E), the normalized discounted cumulative gain (NDCG), the Pearson correlation coefficient (PCC) (G) and the Spearman correlation coefficient (SCC) (H), in which the methods are listed from the best (leftmost) to the worst. Each figure was made on the basis of the scores of 10 repeated cross validation tests for each method. The methods are indexed alphabetically: BMTMKL (1), CaDRReS (2), CDRscan (3), cwKBMF (4), DualNets (5), ENet (6), M-ENet (7), GENet-Lap (8), GENet-NLap (9), KRL (10), KRR (11), MACAU (12), MERGE (13), MVLR (14), pairwiseMKL (15), RF-g (16), RF-gs (17), RWENet-both (18), RWENet-left (19), RWENet-right (20), SRMF (21), TANDEM (22).

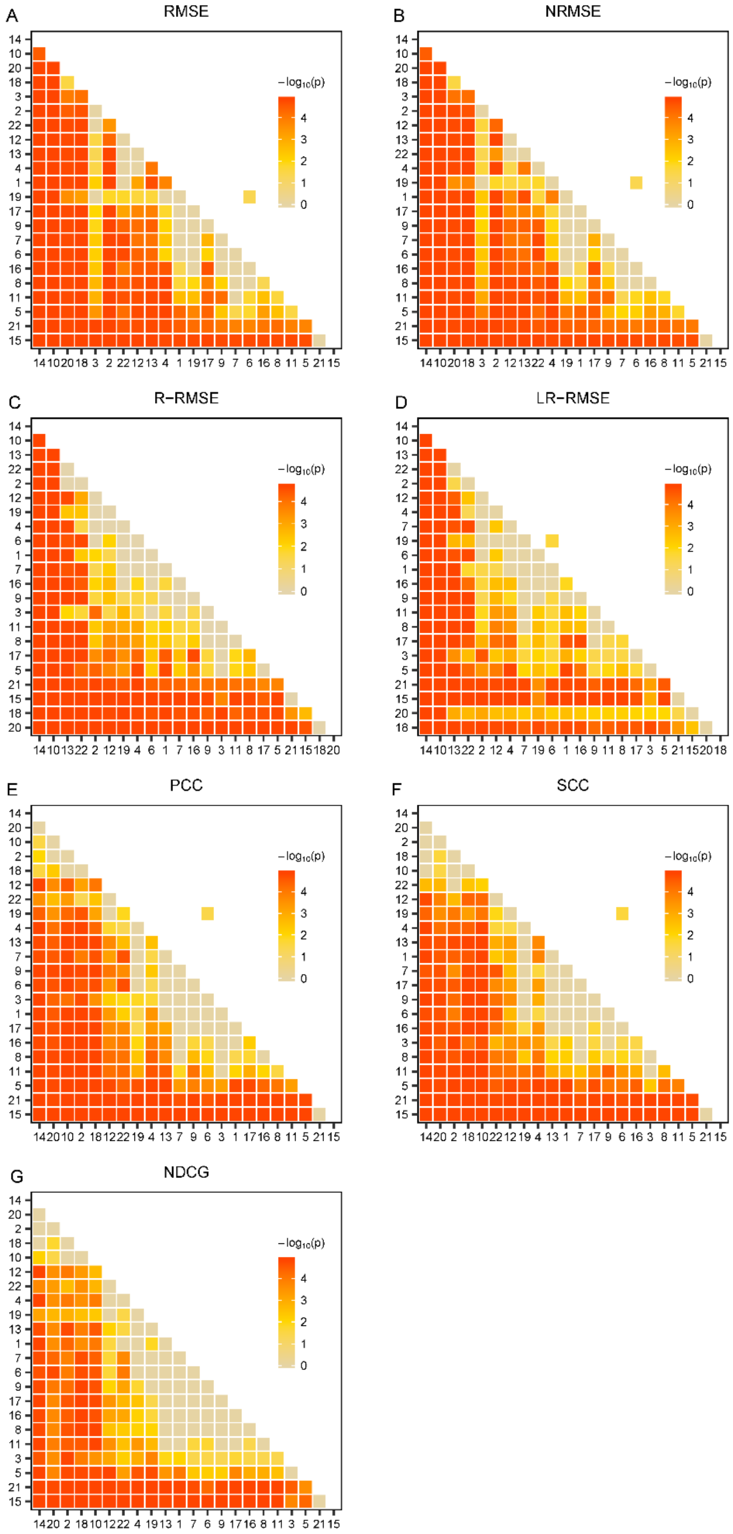


**Figure S26. Performance comparison of the methods on the CCLE AUC dataset.** The one-sided paired Wilcoxon Sign-Rank test was conducted on the basis of the data shown in Figure S1. The heatmap was drawn from the negative common logarithm of the p-value for the hypothesis that a method performs better than another. BMTMKL (1), CaDRReS (2), CDRscan (3), cwKBMF (4), DualNets (5), ENet (6), M-ENet (7), GENet-Lap (8), GENet-NLap (9), KRL (10), KRR (11), MACAU (12), MERGE (13), MVLR (14), pairwiseMKL (15), RF-g (16), RF-gs (17), RWENet-both (18), RWENet-left (19), RWENet-right (20), SRMF (21), TANDEM (22). Note that L-L-RMSE measure were computed for only 5 drugs hence this significance test is not reported for the L-RMSE metric.

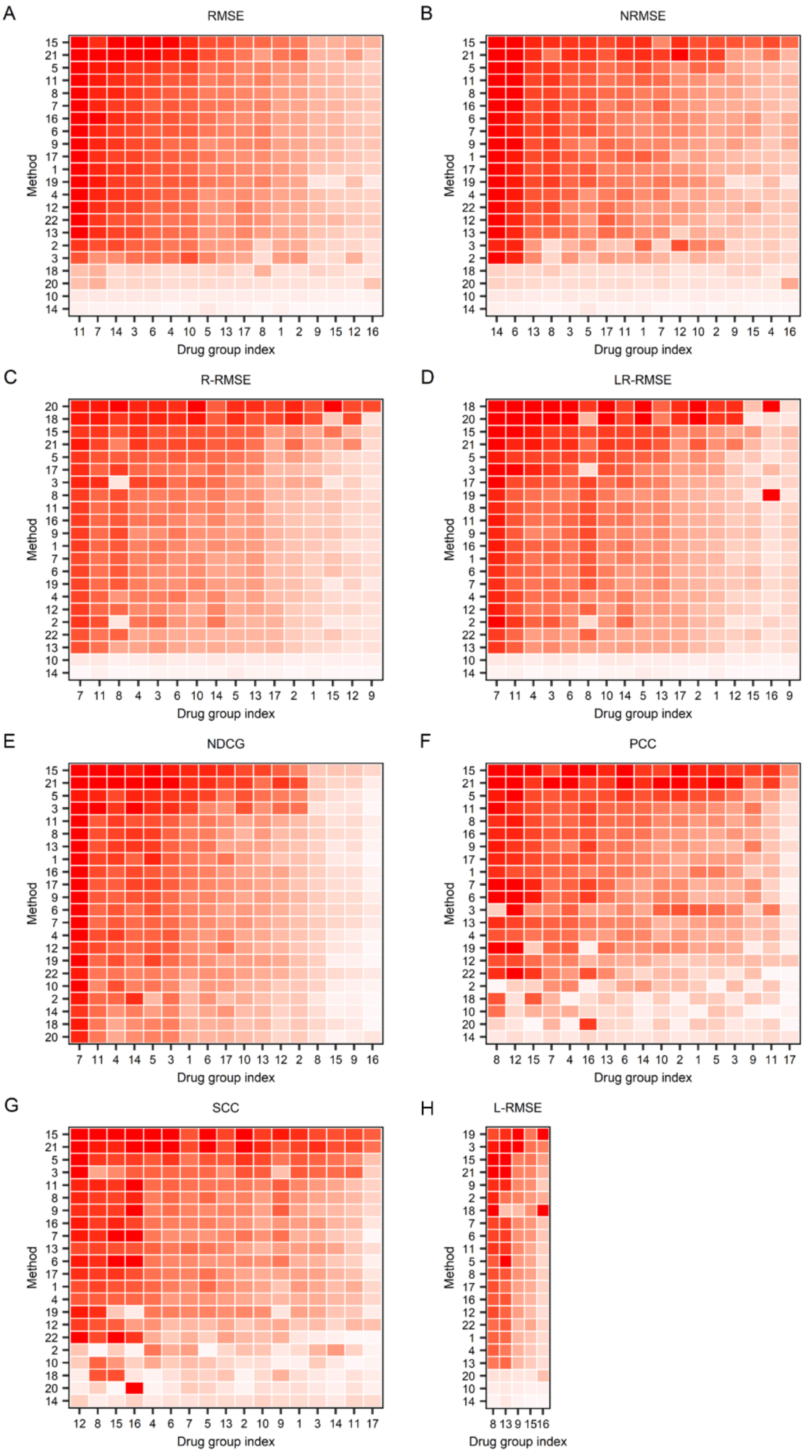


**Figure S27**. **Performance of the methods when trained and tested within 17 drug groups on the CCLE AUC drug response dataset under different metrics**. (Details on next page)

**(Cont'd)**

The root mean square error (RMSE) (A), the normalized RMSE (B), the left-RMSE (C), the right-RMSE (D), the left-right-RMSE (E), the normalized discounted cumulative gain (NDCG) (F), the Pearson correlation coefficient (PCC) (G), and Spearman correlation coefficient (SCC) (H) metrics. Each row and column represent one method and one drug group, respectively. For the method (row) i and drug group (column) j, the median of the measures for drugs in the group j computed by the method I is used to color the corresponding cell, where dark red represents good performance, whereas light red represents bad performance. The methods are indexed alphabetically: BMTMKL (1), CaDRReS (2), CDRscan (3), cwKBMF (4), DualNets (5), ENet (6), M-ENet (7), GENet-Lap (8), GENet-NLap (9), KRL (10), KRR (11), MACAU (12), MERGE (13), MVLR (14), pairwiseMKL(15), RF-g (16), RF-gs (17), RWENet-both (18), RWENet-left (19), RWENet-right (20), SRMF (21), TANDEM (22). Drug groups are also index alphabetically: ABL (1), ALK (2), c-MET (3), CDK4 (4), EGFR (5), FGFR (6), GS (7), HDAC (8), HSP90 (9), IGF1R (10), MDM2 (11), MEK (12), RAF (13), RTK (14), TOP1 (15), TUBB1 (16), XIAP (17).

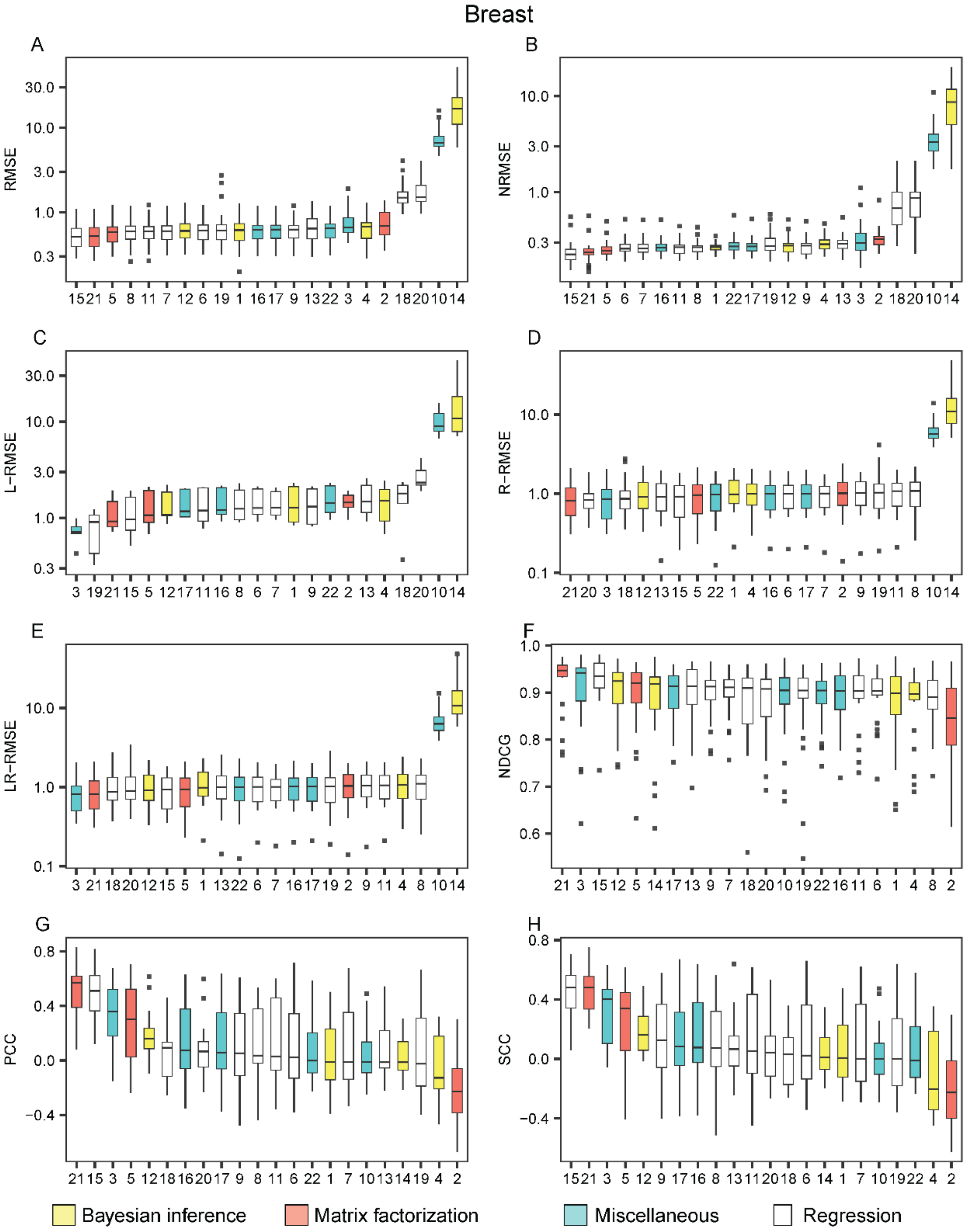


**Figure S28. The performance of the methods on the breast cell lines appearing in the CCLE dataset with AUC drug response readouts**. The methods are listed from the best (leftmost) to the worst in each metric: the root mean square error (RMSE) (A), the normalized RMSE (B), the left-RMSE (C), the right-RMSE (D), the left-right-RMSE (E), the normalized discounted cumulative gain (NDCG), the Pearson correlation coefficient (PCC) (G) and the Spearman correlation coefficient (SCC) (H). The methods are indexed alphabetically: BMTMKL (1), CaDRReS (2), CDRscan (3), cwKBMF (4), DualNets (5), ENet (6), M-ENet (7), GENet-Lap (8), GENet-NLap (9), KRL (10), KRR (11), MACAU (12), MERGE (13), MVLR (14), pairwiseMKL (15), RF-g (16), RF-gs (17), RWENet-both (18), RWENet-left (19), RWENet-right (20), SRMF (21), TANDEM (22).

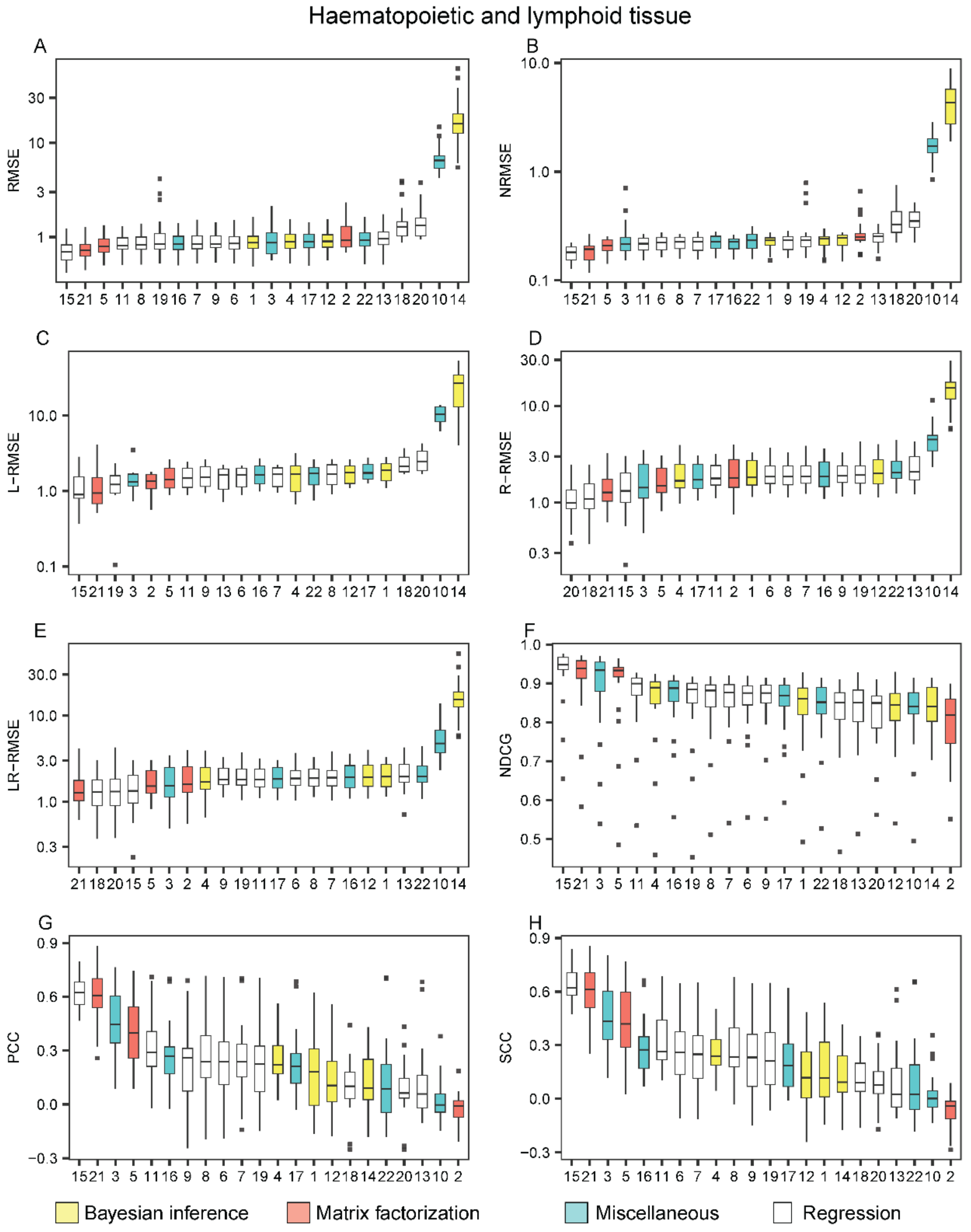


**Figure S29. The performance of the methods on the cell lines in the haematopoietic and lymphoid tissue appearing in the CCLE dataset with AUC drug response readouts**. The methods are listed from the best (leftmost) to the worst in each metric: the root mean square error (RMSE) (A), the normalized RMSE (B), the left-RMSE (C), the right-RMSE (D), the left-right-RMSE (E), the normalized discounted cumulative gain (NDCG), the Pearson correlation coefficient (PCC) (G) and the Spearman correlation coefficient (SCC) (H). The methods are indexed alphabetically: BMTMKL (1), CaDRReS (2), CDRscan (3), cwKBMF (4), DualNets (5), ENet (6), M-ENet (7), GENet-Lap (8), GENet-NLap (9), KRL (10), KRR (11), MACAU (12), MERGE (13), MVLR (14), pairwiseMKL (15), RF-g (16), RF-gs (17), RWENet-both (18), RWENet-left (19), RWENet-right (20), SRMF (21), TANDEM (22).

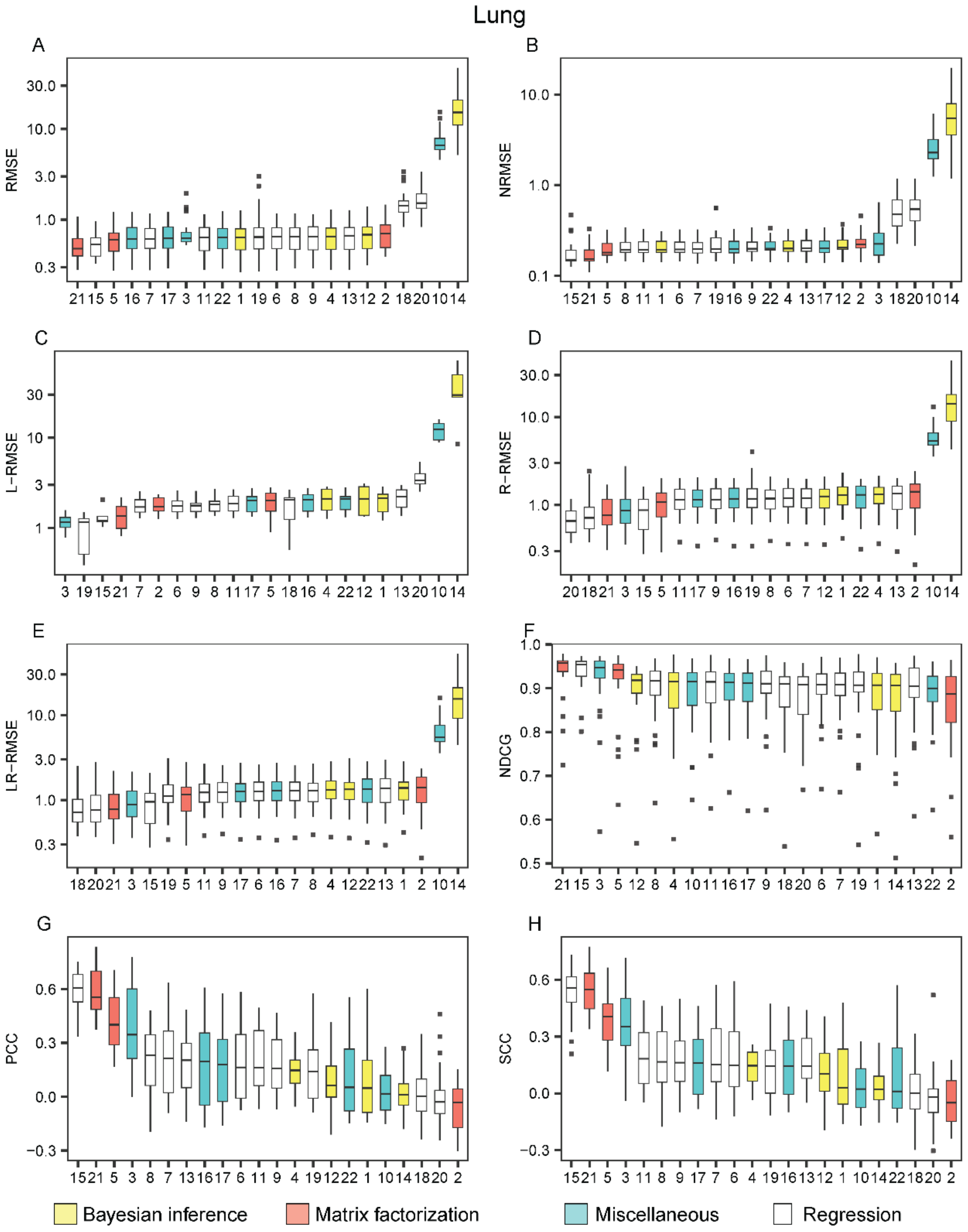


**Figure S30. Performance of the methods on lung cell lines appearing in the CCLE dataset with AUC drug response readouts**. The methods are listed from the best (leftmost) to the worst in each metric: the root mean square error (RMSE) (A), the normalized RMSE (B), the left-RMSE (C), the right-RMSE (D), the left-right-RMSE (E), the normalized discounted cumulative gain (NDCG), the Pearson correlation coefficient (PCC) (G) and the Spearman correlation coefficient (SCC) (H). The methods are indexed alphabetically: BMTMKL (1), CaDRReS (2), CDRscan (3), cwKBMF (4), DualNets (5), ENet (6), M-ENet (7), GENet-Lap (8), GENet-NLap (9), KRL (10), KRR (11), MACAU (12), MERGE (13), MVLR (14), pairwiseMKL (15), RF-g (16), RF-gs (17), RWENet-both (18), RWENet-left (19), RWENet-right (20), SRMF (21), TANDEM (22).

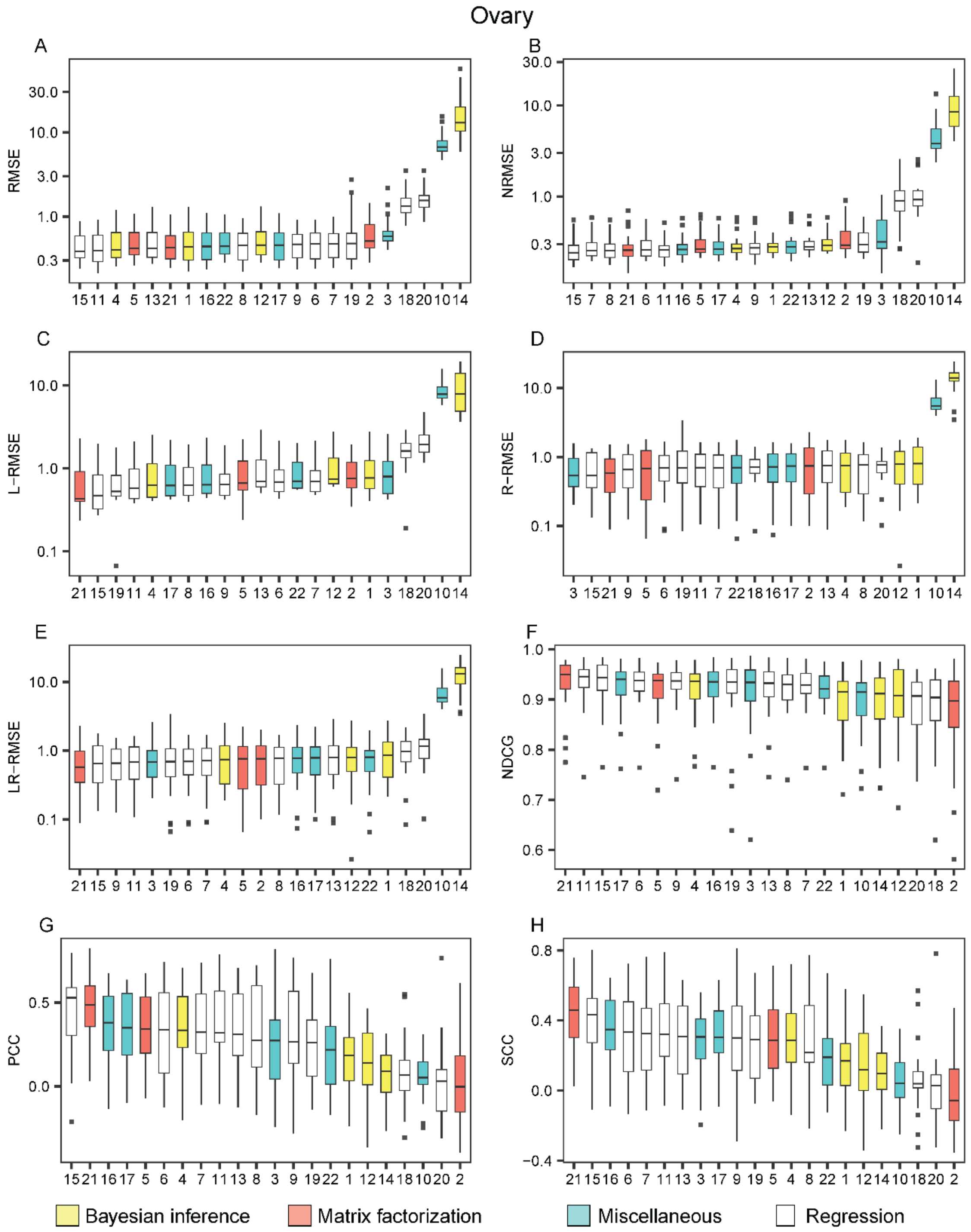


**Figure S31. Performance of the 17 methods on ovary cell lines appearing in the CCLE dataset with AUC drug response readouts**. The methods are listed from the best (leftmost) to the worst in each metric: the root mean square error (RMSE) (A), the normalized RMSE (B), the left-RMSE (C), the right-RMSE (D), the left-right-RMSE (E), the normalized discounted cumulative gain (NDCG), the Pearson correlation coefficient (PCC) (G) and the Spearman correlation coefficient (SCC) (H). The methods are indexed alphabetically: BMTMKL (1), CaDRReS (2), CDRscan (3), cwKBMF (4), DualNets (5), ENet (6), M-ENet (7), GENet-Lap (8), GENet-NLap (9), KRL (10), KRR (11), MACAU (12), MERGE (13), MVLR (14), pairwiseMKL (15), RF-g (16), RF-gs (17), RWENet-both (18), RWENet-left (19), RWENet-right (20), SRMF (21), TANDEM (22).

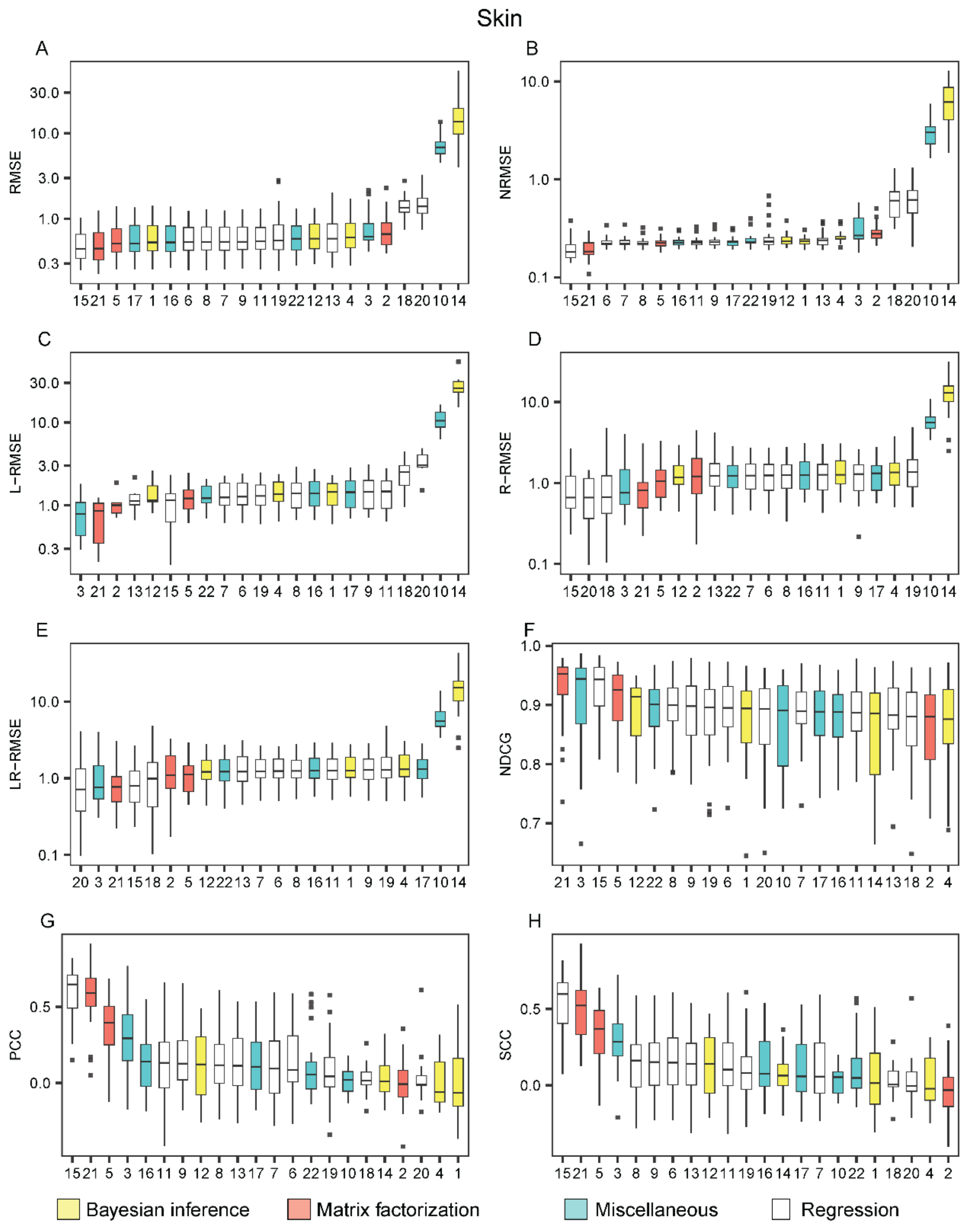


**Figure S32. The performance of the 17 methods on the skin cell lines appearing in the CCLE dataset with AUC drug response readouts**. The methods are listed from the best (leftmost) to the worst in each metric: the root mean square error (RMSE) (A), the normalized RMSE (B), the left-RMSE (C), the right-RMSE (D), the left-right-RMSE (E), the normalized discounted cumulative gain (NDCG) (F), the Pearson correlation coefficient (PCC) (G) and the Spearman correlation coefficient (SCC) (H). The methods are indexed alphabetically: BMTMKL (1), CaDRReS (2), CDRscan (3), cwKBMF (4), DualNets (5), ENet (6), M-ENet (7), GENet-Lap (8), GENet-NLap (9), KRL (10), KRR (11), MACAU (12), MERGE (13), MVLR (14), pairwiseMKL (15), RF-g (16), RF-gs (17), RWENet-both (18), RWENet-left (19), RWENet-right (20), SRMF (21), TANDEM (22).
