## Supplementary File 5 for "A Survey and Systematic Assessment of Computational Methods for Drug Response Prediction"

**Figure S33**.  The performance of the methods on the NCI-60 GI50 drug response dataset under different metrics

**Figure S34**. Performance comparison of the methods on the NCI-60 GI50 drug response dataset.

**Figure S35.** The performance of the methods when trained and tested within 37 drug groups on the NCI-60 GI50  drug response dataset under different metrics.

**Figure S36**. The performance of the methods on lung cell lines appearing in the NCI-60 dataset with GI50 drug response readouts.

**Figure S37**. The performance of the methods on the cell lines of the melanoma tissue appearing in the NCI-60 dataset with GI50 drug response readouts.

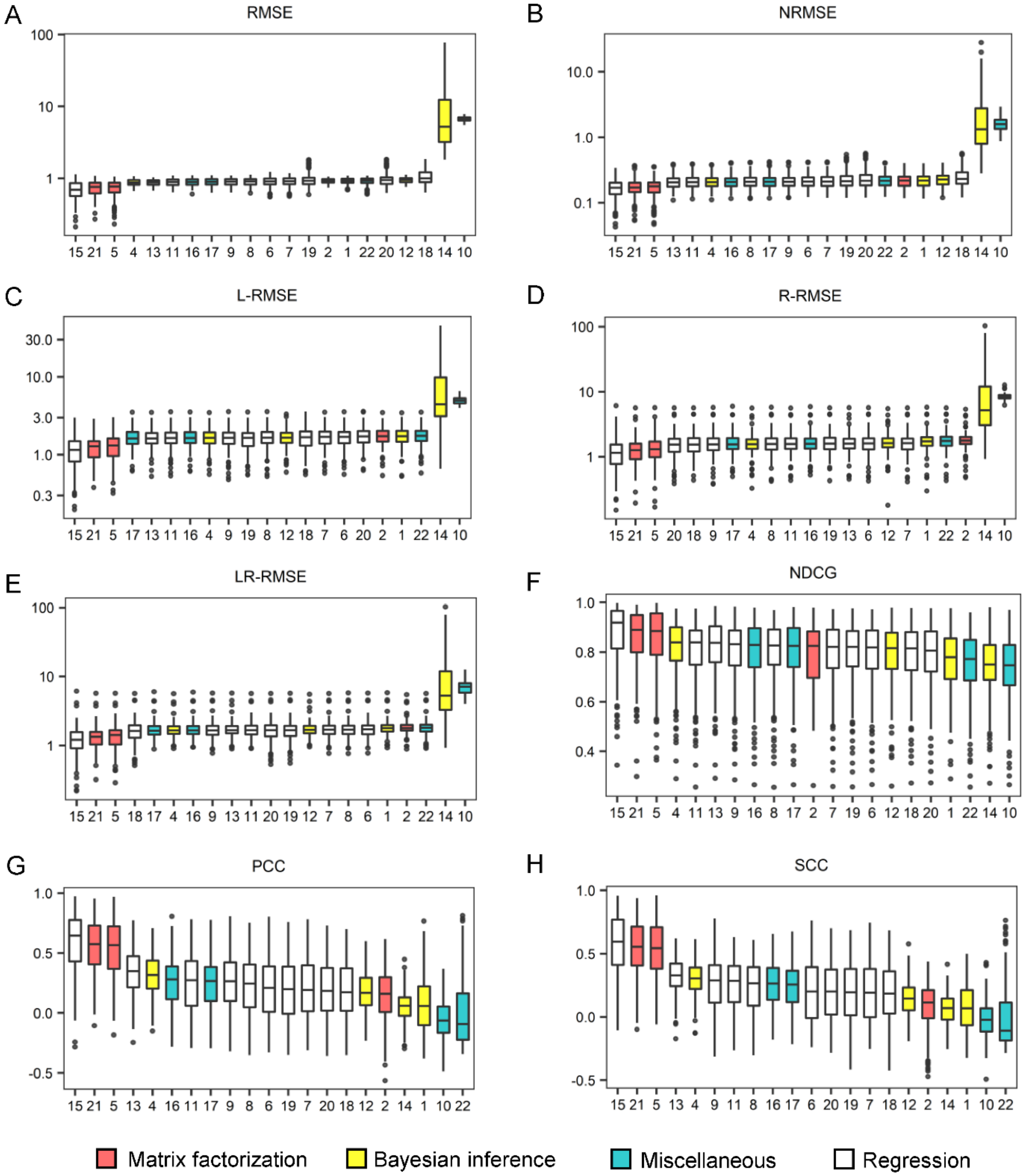


**Figure S33. The performance of the methods on the NCI_60 GI_50 drug response dataset under different metrics.** Box plots were drawn for the root mean square error (RMSE) (A), the normalized RMSE (B), the left-RMSE (C), the right-RMSE (D), the left-right-RMSE (E), the normalized discounted cumulative gain (NDCG), the Pearson correlation coefficient (PCC) (G) and the Spearman correlation coefficient (SCC) (H), in which the methods are listed from the best (leftmost) to the worst. Each figure was made on the basis of the scores of 10 repeated cross validation tests for each method. The methods are indexed alphabetically: BMTMKL (1), CaDRReS (2), CDRscan (3), cwKBMF (4), DualNets (5), ENet (6), M-ENet (7), GENet-Lap (8), GENet-NLap (9), KRL (10), KRR (11), MACAU (12), MERGE (13), MVLR (14), pairwiseMKL (15), RF-g (16), RF-gs (17), RWENet-both (18), RWENet-left (19), RWENet-right (20), SRMF (21), TANDEM (22).

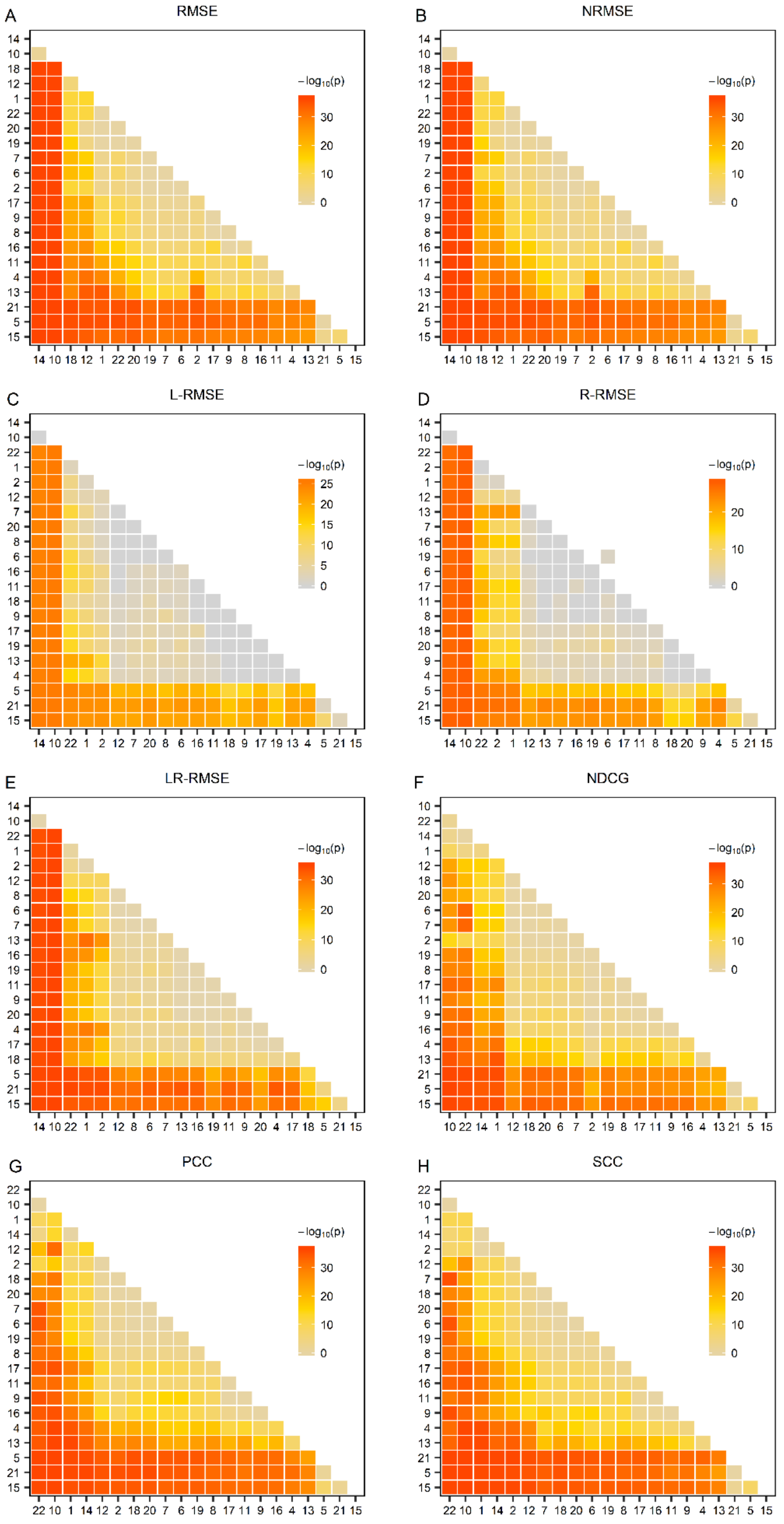


**Figure S34. Performance comparison of the methods on the NCI-60 GI50 drug response dataset.** (Cont’ed on next page)

(Cont'ed) The one-sided paired Wilcoxon Sign-Rank test was conducted on the basis of the data shown in Figure S1. The heatmap was drawn from the negative common logarithm of the p-value for the hypothesis that a method performs better than another. BMTMKL (1), CaDRReS (2), CDRscan (3), cwKBMF (4), DualNets (5), ENet (6), M-ENet (7), GENet-Lap (8), GENet-NLap (9), KRL (10), KRR (11), MACAU (12), MERGE (13), MVLR (14), pairwiseMKL (15), RF-g (16), RF-gs (17), RWENet-both (18), RWENet-left (19), RWENet-right (20), SRMF (21), TANDEM (22).

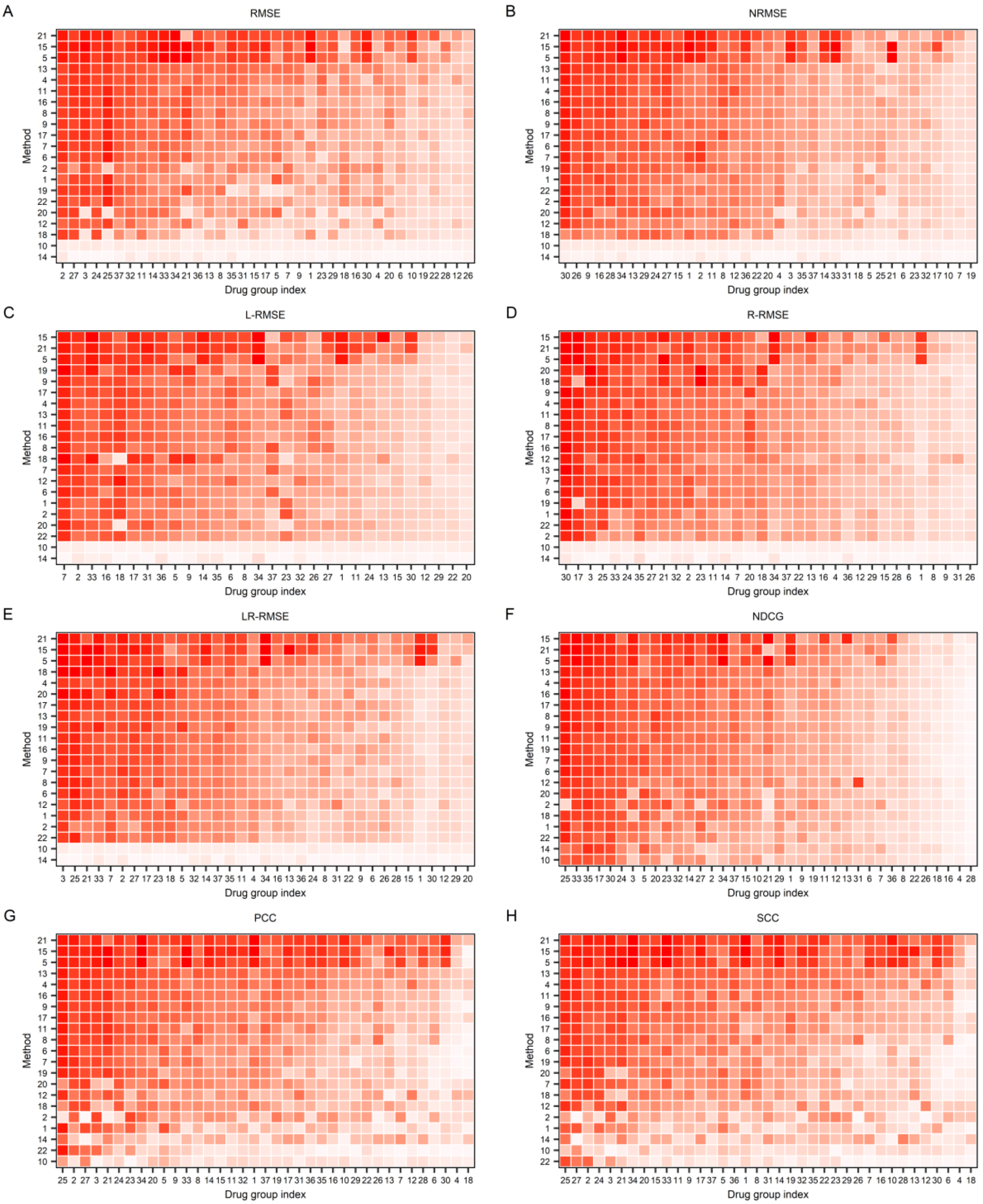


**Figure S35. The performance of the methods when trained and tested within 37 drug groups on the NCI-60 GI50 drug response dataset under different metrics.** (Cont'd on next page)

**(Cont'd)**

The root mean square error (RMSE) (A), the normalized RMSE (B), the left-RMSE (C), the right-RMSE (D), the left-right-RMSE (E), the normalized discounted cumu-lative gain (NDCG) (F), the Pearson correlation coefficient (PCC) (G), and Spearman correlation coefficient (SCC) (H) metrics. Each row and column represent one method and one drug group, respectively. For the method (row) i and drug group (column) j, the median of the measures for drugs in the group j computed by the method I is used to color the corresponding cell, where dark red represents good performance, whereas light red represents bad performance. The methods are indexed alphabetically: BMTMKL (1), CaDRReS (2), CDRscan (3), cwKBMF (4), DualNets (5), ENet (6), M-ENet (7), GENet-Lap (8), GENet-NLap (9), KRL (10), KRR (11), MACAU (12), MERGE (13), MVLR (14), pairwiseMKL(15), RF-g (16), RF-gs (17), RWENet-both (18), RWENet-left (19), RWENet-right (20), SRMF (21), TANDEM (22). Drug groups are also index alphabetically: Alkylating at N-2 position of guanine (1), Alkylating at N-7 position of guanine (2), Alkylating at O-6 of guanine (3), Angiogenesis (4), Antibiotic (5), Antibodies (6), Antifols (7), Antimetabolite (8), Apoptosis inducer (9), Cell cycle (10), DNA binder (11), DNA damage repair (12), DNA methylttransferase inhibitor (13), DNA synthesis inhibitor (14), Histone deacetylase (15), Hormone (16), HSP90 (17), Hypoxia (18), Mito-chondria; Warburg e ect (19), Cardiac glycoside (20), Cathartic (21), Antipsychotic (22), Nuclear factor of kappa light polypeptide gene enhancer in B-cells (23), Phosphatase inhibitor (24), BRAF inhibitor (25), PIK3 inhibitor (26), PRKCA (PRK) inhibitor (27), Poly (ADP-ribose) polymerase (28), Protea-some inhibitor (29), RNA synthesis inhibitor (30), ROS1 inhibitor (31), Serine threonine kinase (32), Topoisomerase 1 inhibitor (33), Topoisomerase 2 inhibitor (34), Tubulin affecting (35), Tyrosine kinase inhibitor (36), Unknown (37).

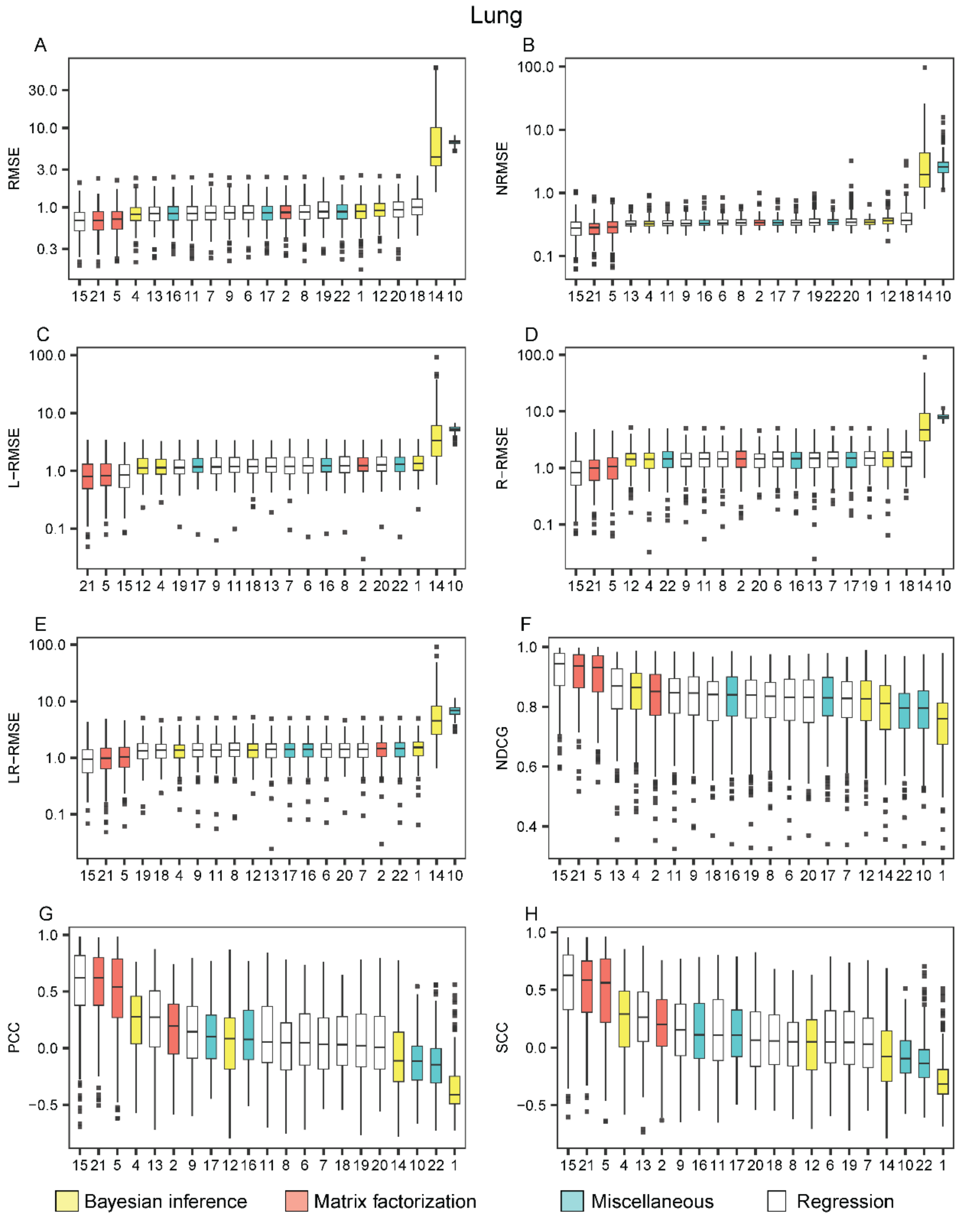


**Figure S36. The performance of the methods on lung cell lines appearing in the NCI-60 dataset with GI50 drug response readouts**. The methods are listed from the best (leftmost) to the worst in each metric: the root mean square error (RMSE) (A), the normalized RMSE (B), the left-RMSE (C), the right-RMSE (D), the left-right-RMSE (E), the normalized discounted cumulative gain (NDCG), the Pearson correlation coefficient (PCC) (G) and the Spearman correlation coefficient (SCC) (H). The methods are indexed alphabetically: BMTMKL (1), CaDRReS (2), CDRscan (3), cwKBMF (4), DualNets (5), ENet (6), M-ENet (7), GENet-Lap (8), GENet-NLap (9), KRL (10), KRR (11), MACAU (12), MERGE (13), MVLR (14), pairwiseMKL (15), RF-g (16), RF-gs (17), RWENet-both (18), RWENet-left (19), RWENet-right (20), SRMF (21), TANDEM (22).

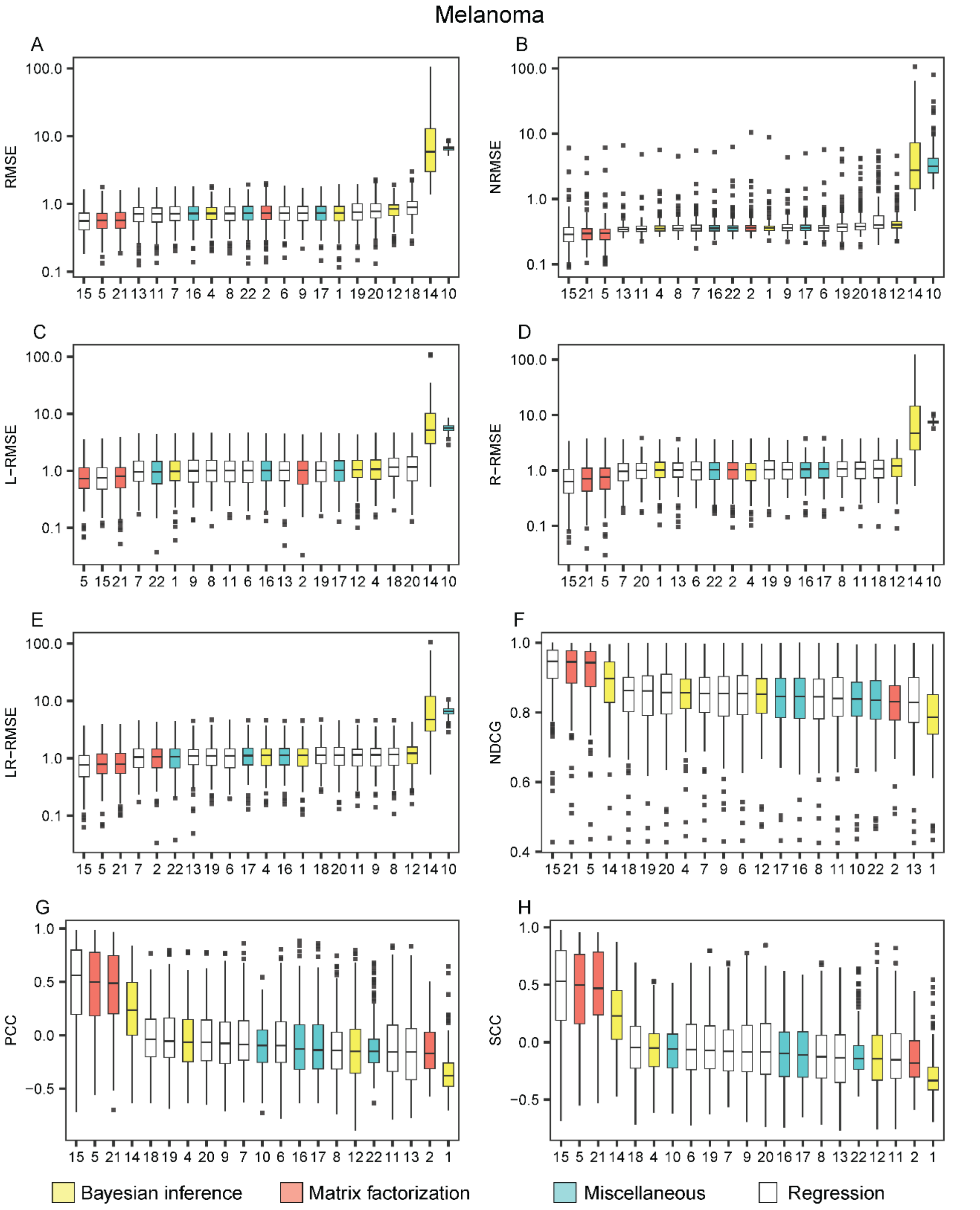


**Figure S37. The performance of the methods on the cell lines of the melanoma tissue appearing in the NCI-60 dataset with GI50 drug response readouts**. The methods are listed from the best (leftmost) to the worst in each metric: the root mean square error (RMSE) (A), the normalized RMSE (B), the left-RMSE (C), the right-RMSE (D), the left-right-RMSE (E), the normalized discounted cumulative gain (NDCG), the Pearson correlation coefficient (PCC) (G) and the Spearman correlation coefficient (SCC) (H). The methods are indexed alphabetically: BMTMKL (1), CaDRReS (2), CDRscan (3), cwKBMF (4), DualNets (5), ENet (6), M-ENet (7), GENet-Lap (8), GENet-NLap (9), KRL (10), KRR (11), MACAU (12), MERGE (13), MVLR (14), pairwiseMKL (15), RF-g (16), RF-gs (17), RWENet-both (18), RWENet-left (19), RWENet-right (20), SRMF (21), TANDEM (22).
