## Supplementary File 6 for "A Survey and Systematic Assessment of Computational Methods for Drug Response Prediction"

**Figure S38**. The performance of the methods when trained on the GDSC IC50 dataset and tested on the CCLE IC50 dataset under different metrics.

**Figure S39**. The performance of the methods when trained and tested within 5 common drug groups if trained on the GDSC IC50 dataset and tested on the CCLE IC50 drug response dataset under different methods.

**Figure S40**. The performance of the methods on the breast cell lines when trained on the GDSC IC50 dataset and tested on the CCLE IC50 dataset.

**Figure S41**. The performance of the methods on the cell lines of haematopoietic and lymphoid tissue when trained on the GDSC IC50 dataset and tested on the CCLE IC50 dataset.

**Figure S42**. The performance of the methods on the ovary cell lines when trained on the GDSC IC50 dataset and tested on CCLE IC50 dataset.

**Figure S43**. The performance of the methods on the lung cell lines when trained on the GDSC IC50 dataset and tested on CCLE IC50 dataset.

**Figure S44**. The performance of the methods on the skin cell lines when trained on the GDSC IC50 dataset and tested on CCLE IC50 dataset

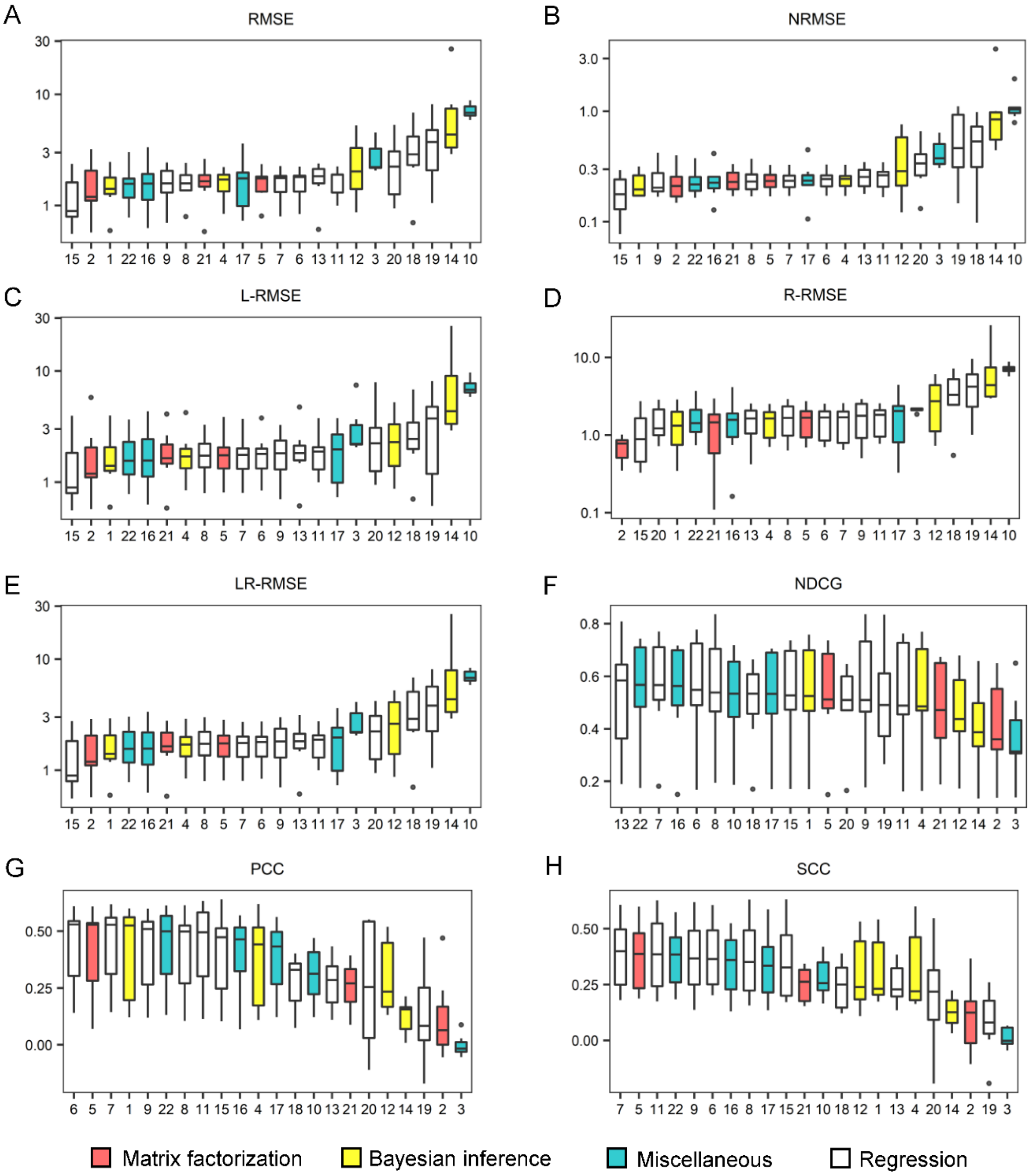


**Figure S38. The performance of the methods when trained on the GDSC IC50 dataset and tested on the CCLE IC50 dataset under different metrics.** Box plots were drawn for the root mean square error (RMSE) (A), the normalized RMSE (B), the left-RMSE (C), the right-RMSE (D), the left-right-RMSE (E), the normalized discounted cumulative gain (NDCG), the Pearson correlation coefficient (PCC) (G) and the Spearman correlation coefficient (SCC) (H), in which the methods are listed from the best (leftmost) to the worst. Each figure was made on the basis of the scores of 10 repeated cross validation tests for each method. The methods are indexed alphabetically: BMTMKL (1), CaDRReS (2), CDRscan (3), cwKBMF (4), DualNets (5), ENet (6), M-ENet (7), GENet-Lap (8), GENet-NLap (9), KRL (10), KRR (11), MACAU (12), MERGE (13), MVLR (14), pairwiseMKL (15), RF-g (16), RF-gs (17), RWENet-both (18), RWENet-left (19), RWENet-right (20), SRMF (21), TANDEM (22).

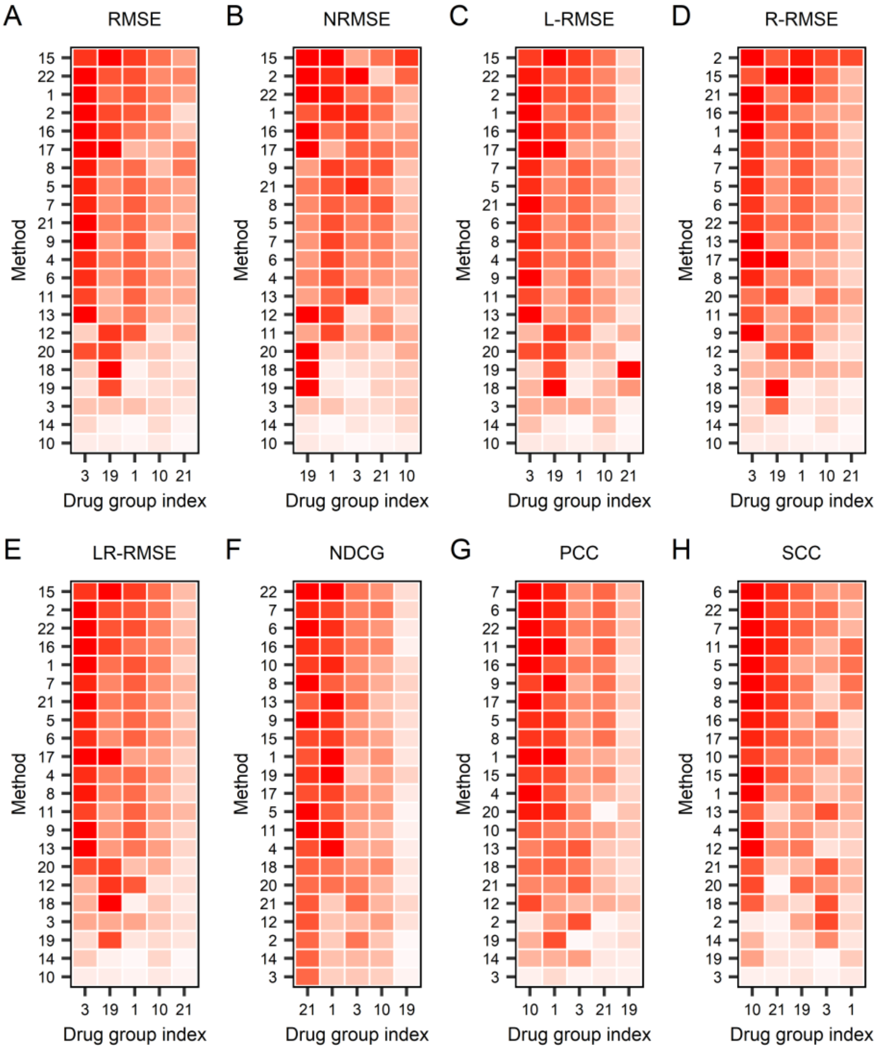


**Figure S39**. **The performance of the methods when trained and tested within 5 common drug groups if trained on the GDSC IC50 dataset and tested on the CCLE IC50 drug response dataset under different methods**. The root mean square error (RMSE) (A), the normalized RMSE (B), the left-RMSE (C), the right-RMSE (D), the left-right-RMSE (E), the normalized discounted cumulative gain (NDCG) (F), the Pearson correlation coefficient (PCC) (G), and Spearman correlation coefficient (SCC) (H) metrics. Each row and column represent one method and one drug group, respectively. For the method (row) i and drug group (column) j, the median of the measures for drugs in the group j computed by the method I is used to color the corresponding cell, where dark red represents good performance, whereas light red represents bad performance. The methods are indexed alphabetically: BMTMKL (1), CaDRReS (2), CDRscan (3), cwKBMF (4), DualNets (5), ENet (6), M-ENet (7), GENet-Lap (8), GENet-NLap (9), KRL (10), KRR (11), MACAU (12), MERGE (13), MVLR (14), pairwiseMKL(15), RF-g (16), RF-gs (17), RWENet-both (18), RWENet-left (19), RWENet-right (20), SRMF (21), TANDEM (22). Drug groups are also indexed as follows: ABL signaling (1), Cell cycle(3), ERK MAPK signaling(10), p53 pathway(19), Protein stability and degradation(21). The evaluation on other groups cannot be properly done.

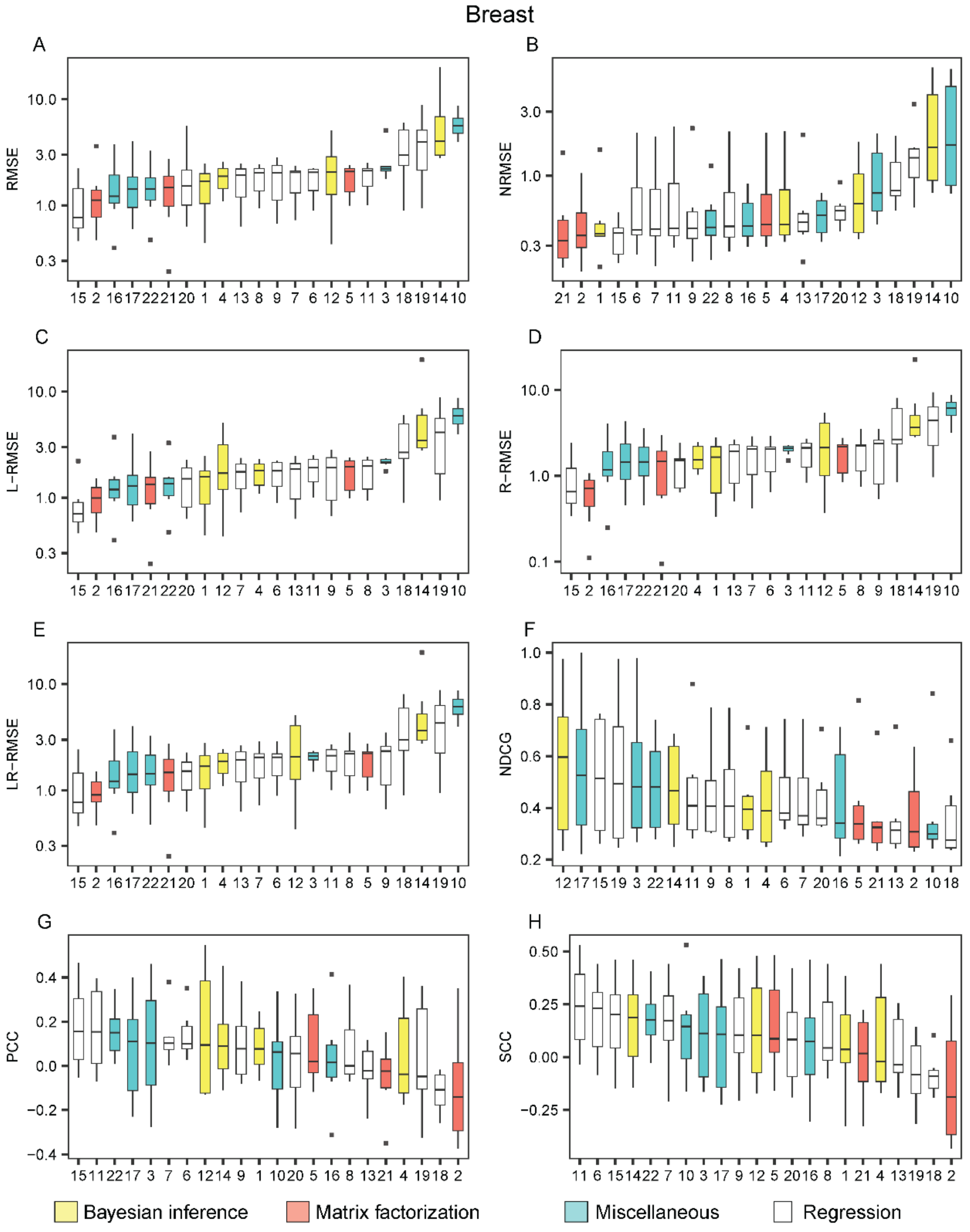


**Figure S40. The performance of the methods on the breast cell lines when trained on the GDSC IC50 dataset and tested on the CCLE IC50 dataset**. The methods are listed from the best (leftmost) to the worst in each metric: the root mean square error (RMSE) (A), the normalized RMSE (B), the left-RMSE (C), the right-RMSE (D), the left-right-RMSE (E), the normalized discounted cumulative gain (NDCG), the Pearson correlation coefficient (PCC) (G) and the Spearman correlation coefficient (SCC) (H). The methods are indexed alphabetically: BMTMKL (1), CaDRReS (2), CDRscan (3), cwKBMF (4), DualNets (5), ENet (6), M-ENet (7), GENet-Lap (8), GENet-NLap (9), KRL (10), KRR (11), MACAU (12), MERGE (13), MVLR (14), pairwiseMKL (15), RF-g (16), RF-gs (17), RWENet-both (18), RWENet-left (19), RWENet-right (20), SRMF (21), TANDEM (22).

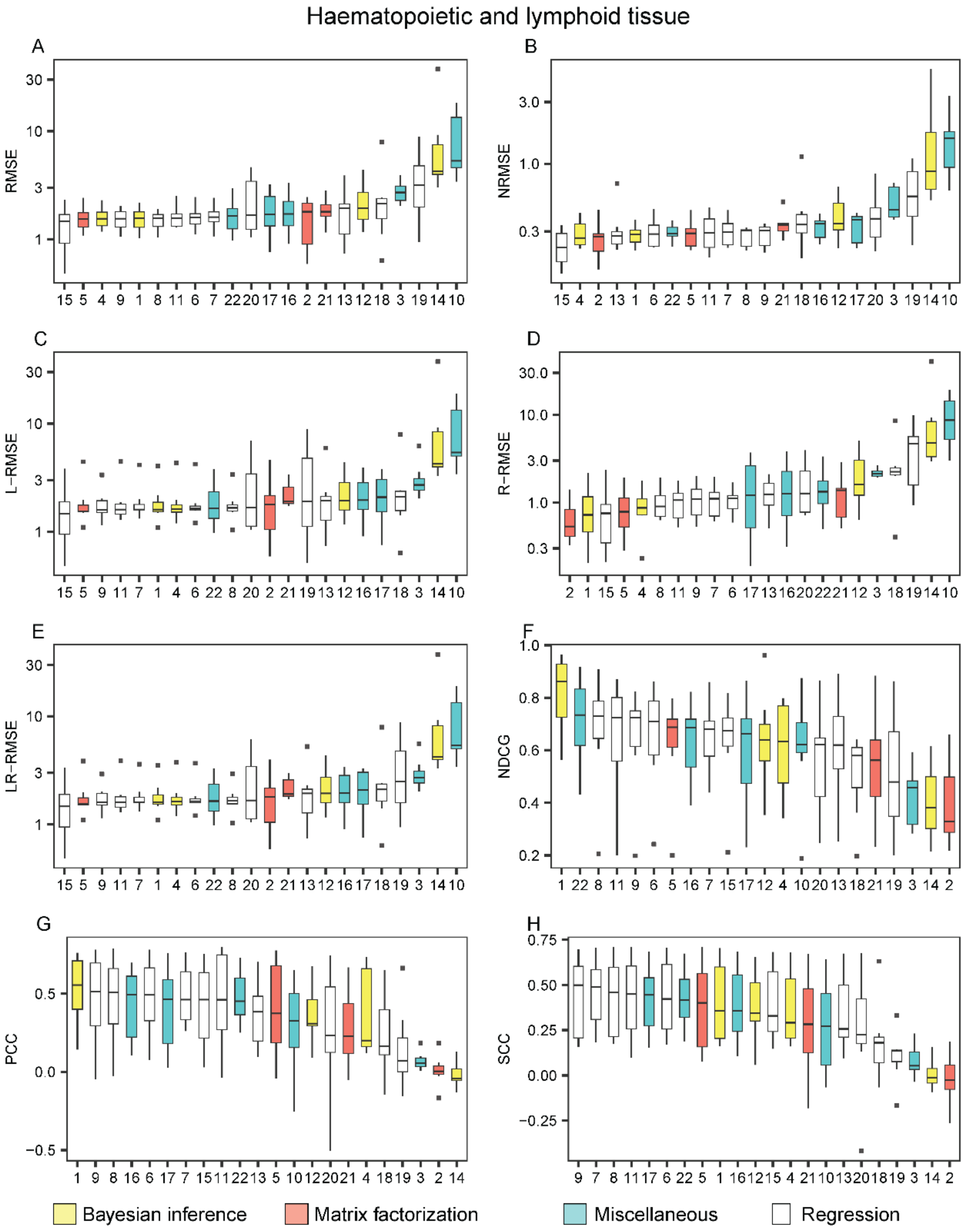


**Figure S41. The performance of the methods on the cell lines of haematopoietic and lymphoid tissue when trained on the GDSC IC50 dataset and tested on the CCLE IC50 dataset**. The methods are listed from the best (leftmost) to the worst in each metric: the root mean square error (RMSE) (A), the normalized RMSE (B), the left-RMSE (C), the right-RMSE (D), the left-right-RMSE (E), the normalized discounted cumulative gain (NDCG), the Pearson correlation coefficient (PCC) (G) and the Spearman correlation coefficient (SCC) (H). The methods are indexed alphabetically: BMTMKL (1), CaDRReS (2), CDRscan (3), cwKBMF (4), DualNets (5), ENet (6), M-ENet (7), GENet-Lap (8), GENet-NLap (9), KRL (10), KRR (11), MACAU (12), MERGE (13), MVLR (14), pairwiseMKL (15), RF-g (16), RF-gs (17), RWENet-both (18), RWENet-left (19), RWENet-right (20), SRMF (21), TANDEM (22).

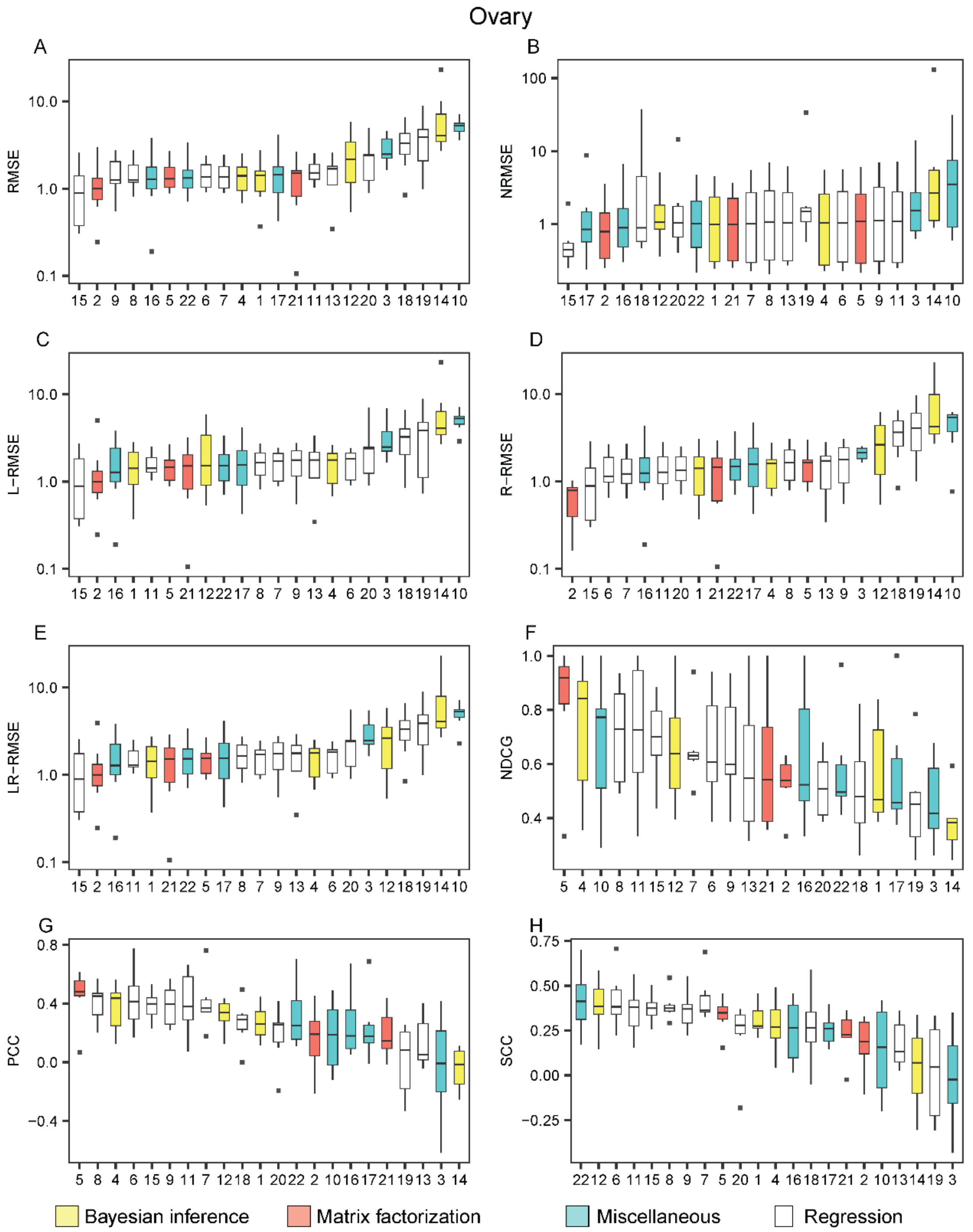


**Figure S42. The performance of the methods on the ovary cell lines when trained on the GDSC IC50 dataset and tested on CCLE IC50 dataset**. The methods are listed from the best (leftmost) to the worst in each metric: the root mean square error (RMSE) (A), the normalized RMSE (B), the left-RMSE (C), the right-RMSE (D), the left-right-RMSE (E), the normalized discounted cumulative gain (NDCG), the Pearson correlation coefficient (PCC) (G) and the Spearman correlation coefficient (SCC) (H). The methods are indexed alphabetically: BMTMKL (1), CaDRReS (2), CDRscan (3), cwKBMF (4), DualNets (5), ENet (6), M-ENet (7), GENet-Lap (8), GENet-NLap (9), KRL (10), KRR (11), MACAU (12), MERGE (13), MVLR (14), pairwiseMKL (15), RF-g (16), RF-gs (17), RWENet-both (18), RWENet-left (19), RWENet-right (20), SRMF (21), TANDEM (22).

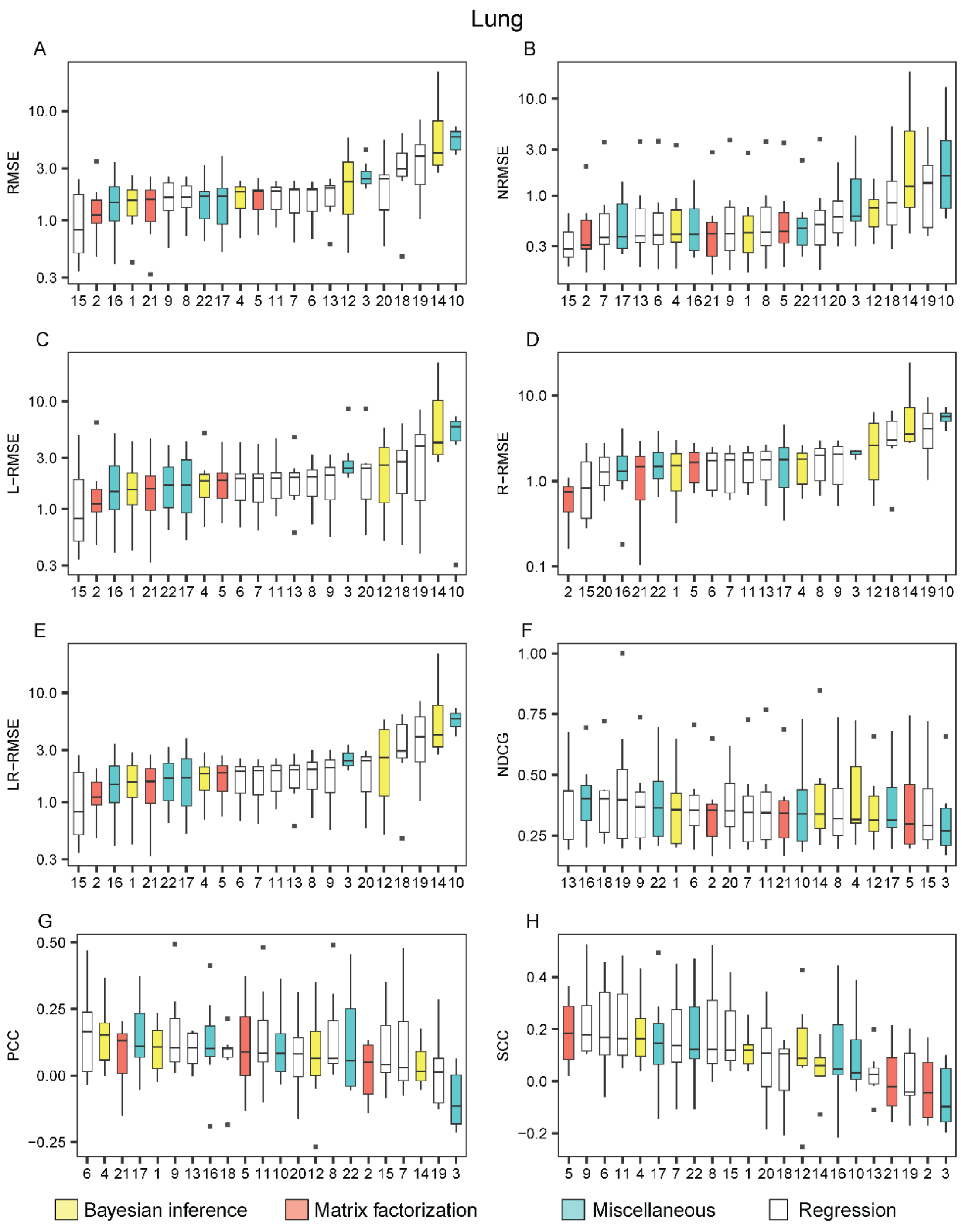


**Figure S43. The performance of the methods on the lung cell lines when trained on the GDSC IC50 dataset and tested on CCLE IC50 dataset**. The methods are listed from the best (leftmost) to the worst in each metric: the root mean square error (RMSE) (A), the normalized RMSE (B), the left-RMSE (C), the right-RMSE (D), the left-right-RMSE (E), the normalized discounted cumulative gain (NDCG), the Pearson correlation coefficient (PCC) (G) and the Spearman correlation coefficient (SCC) (H). The methods are indexed alphabetically: BMTMKL (1), CaDRReS (2), CDRscan (3), cwKBMF (4), DualNets (5), ENet (6), M-ENet (7), GENet-Lap (8), GENet-NLap (9), KRL (10), KRR (11), MACAU (12), MERGE (13), MVLR (14), pairwiseMKL (15), RF-g (16), RF-gs (17), RWENet-both (18), RWENet-left (19), RWENet-right (20), SRMF (21), TANDEM (22).

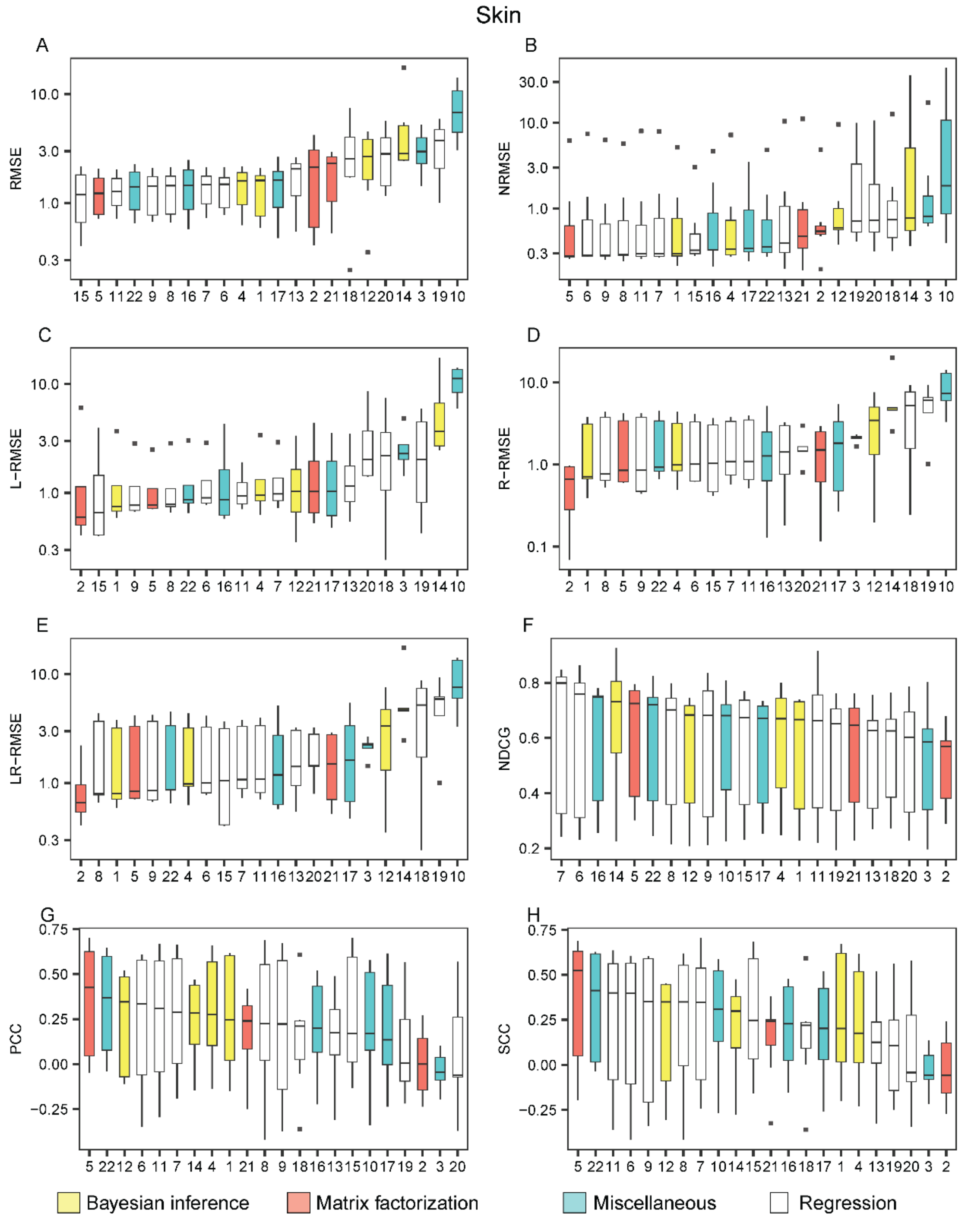


**Figure S44. The performance of the methods on the skin cell lines when trained on the GDSC IC50 dataset and tested on CCLE IC50 dataset**. The methods are listed from the best (leftmost) to the worst in each metric: the root mean square error (RMSE) (A), the normalized RMSE (B), the left-RMSE (C), the right-RMSE (D), the left-right-RMSE (E), the normalized discounted cumulative gain (NDCG), the Pearson correlation coefficient (PCC) (G) and the Spearman correlation coefficient (SCC) (H). The methods are indexed alphabetically: BMTMKL (1), CaDRReS (2), CDRscan (3), cwKBMF (4), DualNets (5), ENet (6), M-ENet (7), GENet-Lap (8), GENet-NLap (9), KRL (10), KRR (11), MACAU (12), MERGE (13), MVLR (14), pairwiseMKL (15), RF-g (16), RF-gs (17), RWENet-both (18), RWENet-left (19), RWENet-right (20), SRMF (21), TANDEM (22).
