## Supplementary File 7 for "A Survey and Systematic Assessment of Computational Methods for Drug Response Prediction"

**Figure S45**. The performance of the methods if trained on the GDSC AUC dataset and tested on the CCLE AUC dataset under different metrics.

**Figure S46**. The performance of the methods within five common drug groups when trained on the GDSC AUC dataset and tested on the CCLE AUC dataset under different metrics.

**Figure S47**. The performance of the methods on the breast cell lines when trained on the GDSC AUC dataset and tested on the CCLE AUC dataset

**Figure S48**. The performance of the methods on the cell lines of haematopoietic and lymphoid tissue when trained on the GDSC AUC dataset and tested on the CCLE AUC dataset..

**Figure S49**. The performance of the methods on the lung cell lines when trained on the GDSC AUC dataset and tested on CCLE AUC dataset.

**Figure S50**. The performance of the methods on the ovary cell lines when trained on the GDSC AUC dataset and tested on CCLE AUC dataset.

**Figure S51**. The performance of the methods on the skin cell lines when trained on the GDSC AUC dataset and tested on CCLE AUC dataset

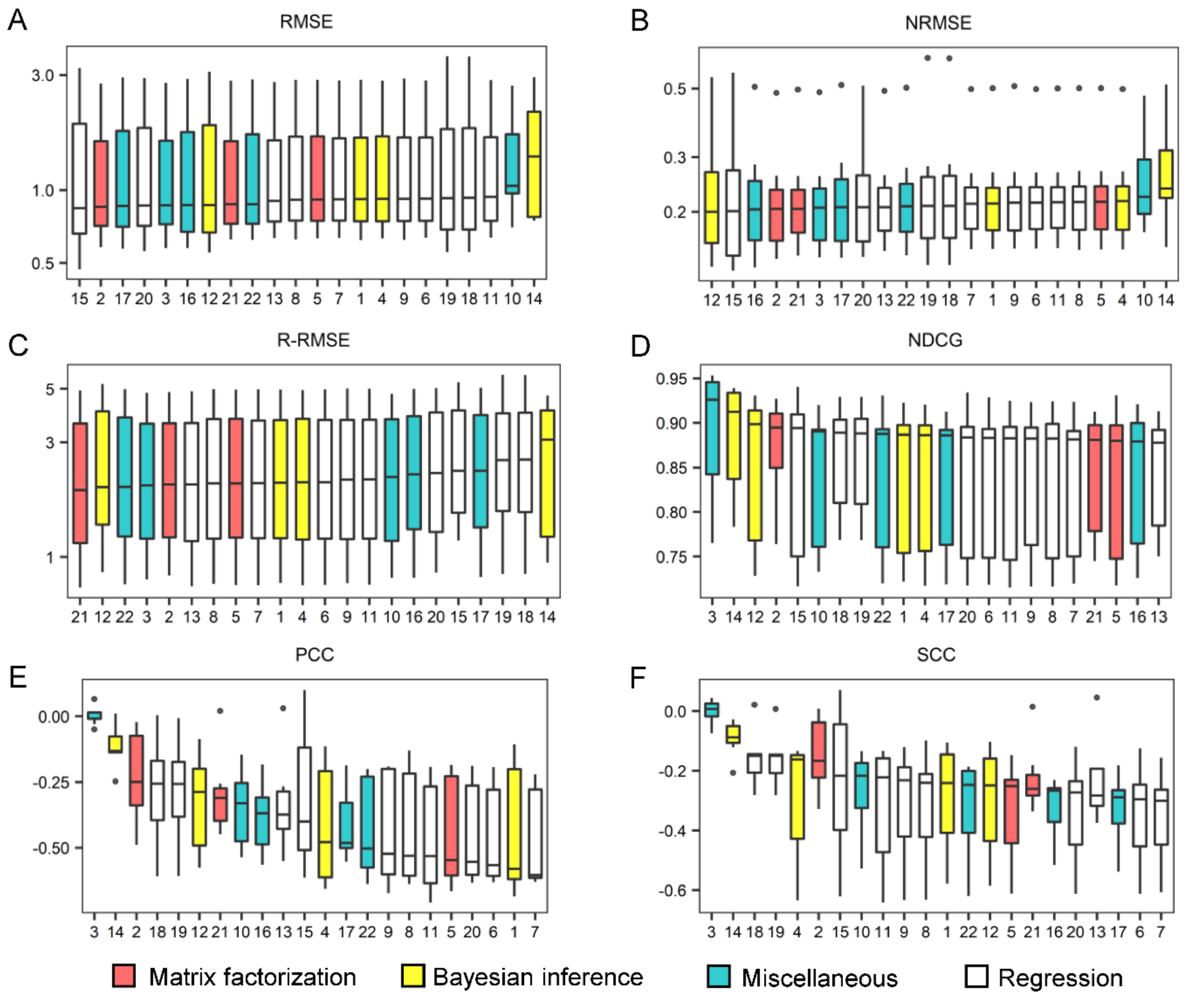


**Figure S45. The performance of the methods if trained on the GDSC AUC dataset and tested on the CCLE AUC dataset under different metrics.** Box plots were drawn for the root mean square error (RMSE) (A), the normalized RMSE (B), the right-RMSE (C), the normalized discounted cumulative gain (NDCG) (D), the Pearson correlation coefficient (PCC) (E) and the Spearman correlation coefficient (SCC) (F), in which the methods are listed from the best (leftmost) to the worst. Each figure was made on the basis of the scores of 10 repeated cross validation tests for each method. The methods are indexed alphabetically: BMTMKL (1), CaDRReS (2), CDRscan (3), cwKBMF (4), DualNets (5), ENet (6), M-ENet (7), GENet-Lap (8), GENet-NLap (9), KRL (10), KRR (11), MACAU (12), MERGE (13), MVLR (14), pairwiseMKL (15), RF-g (16), RF-gs (17), RWENet-both (18), RWENet-left (19), RWENet-right (20), SRMF (21), TANDEM (22). For this test setting, the evaluation is not available for the L-RMSE metric and thus the evaluation results were identical for R-RMSE and LR-RMSE.

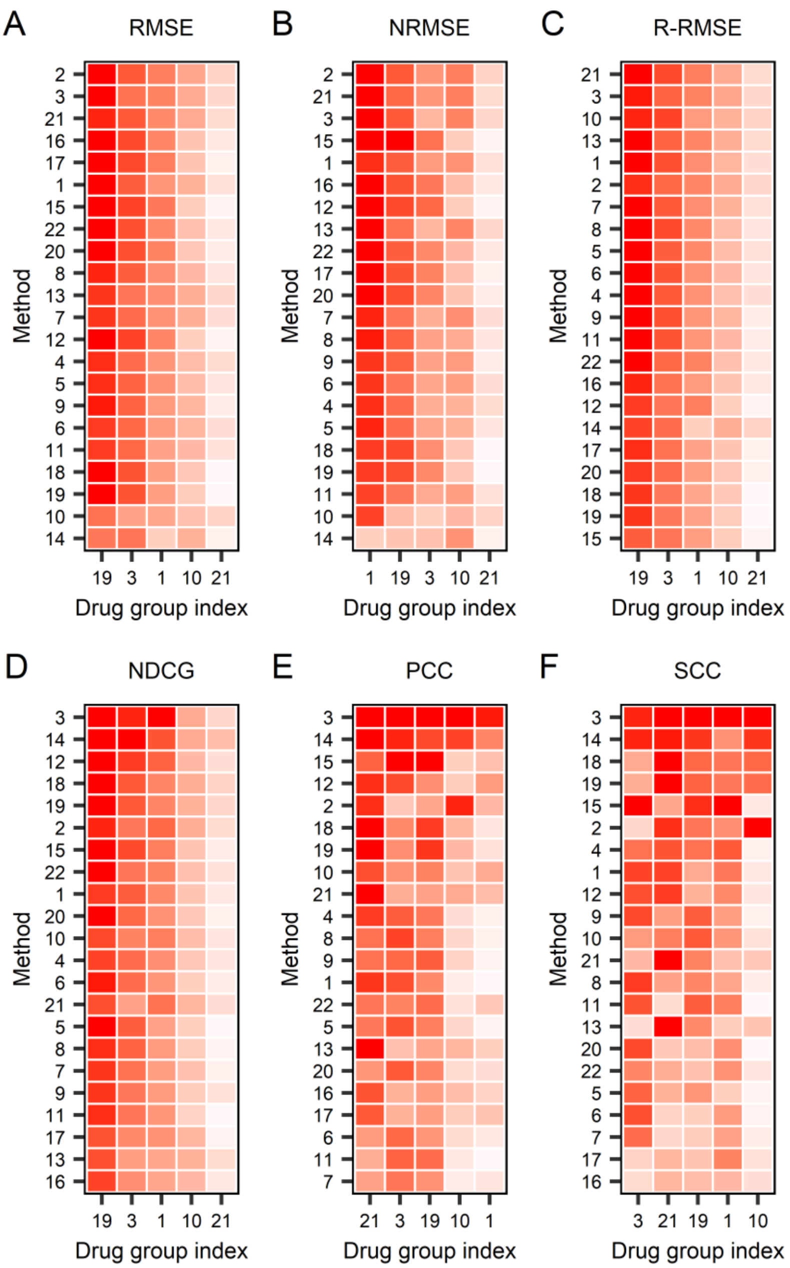


**Figure S46**. **The performance of the methods within five common drug groups when trained on the GDSC AUC dataset and tested on the CCLE AUC dataset under different metrics**. The root mean square error (RMSE) (A), the normalized RMSE (B), the right-RMSE (C), the normalized discounted cumulative gain (NDCG) (D), the Pearson correlation coefficient (PCC) (E), and Spearman correlation coefficient (SCC) (F). Each row and column represent one method and one drug group, respectively. For the method (row) i and drug group (column) j, the median of the measures for drugs in the group j computed by the method I is used to color the corresponding cell, where dark red represents good performance, whereas light red represents bad performance. The methods are indexed alphabetically: BMTMKL (1), CaDRReS (2), CDRscan (3), cwKBMF (4), DualNets (5), ENet (6), M-ENet (7), GENet-Lap (8), GENet-NLap (9), KRL (10), KRR (11), MACAU (12), MERGE (13), MVLR (14), pairwiseMKL(15), RF-g (16), RF-gs (17), RWENet-both (18), RWENet-left (19), RWENet-right (20), SRMF (21), TANDEM (22). Drug groups are also indexed as follows: ABL signaling (1), Cell cycle(3), ERK MAPK signaling(10), p53 pathway(19), Protein stability and degradation(21). The data on the other drug groups were not sufficient for evaluation.

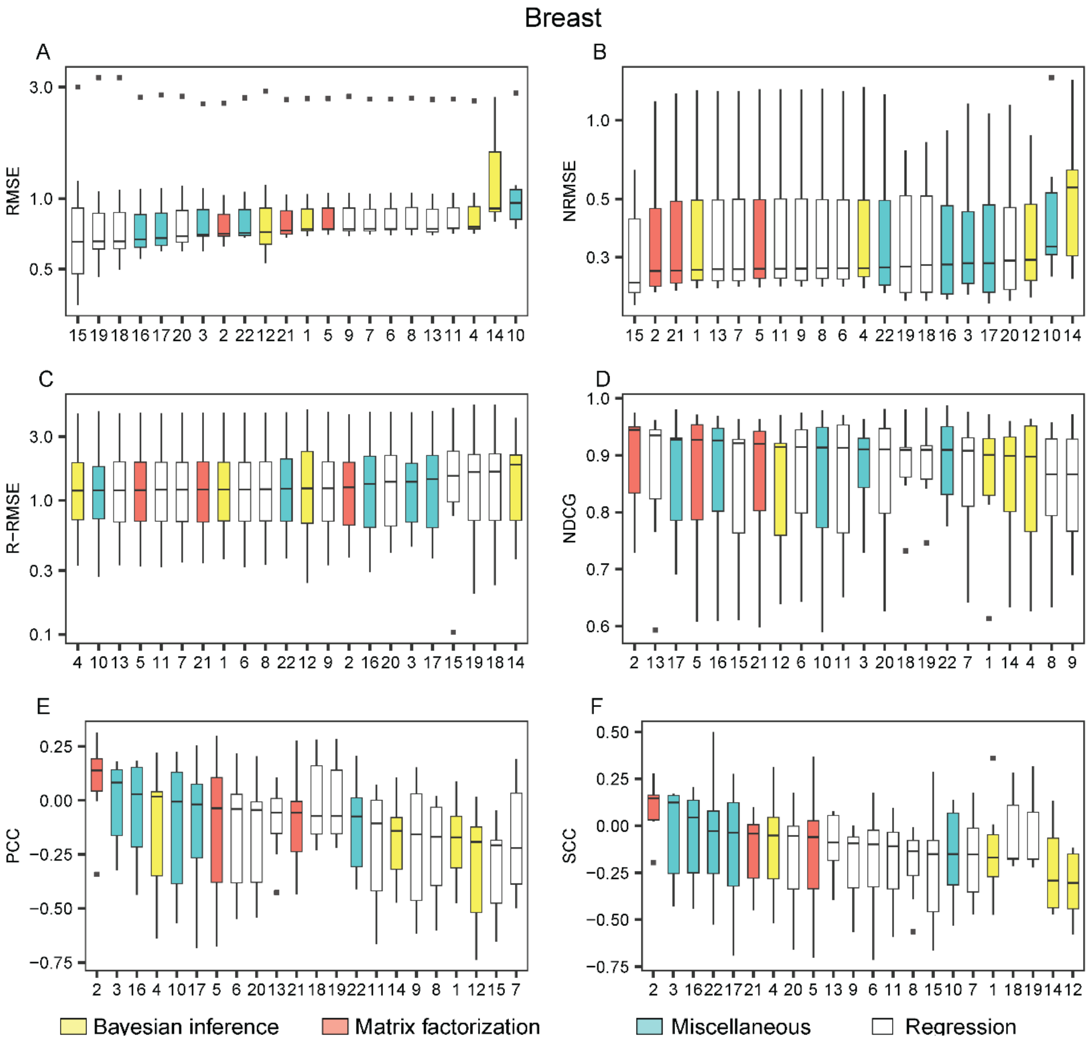


**Figure S47. The performance of the methods on the breast cell lines when trained on the GDSC AUC dataset and tested on the CCLE AUC dataset**. The methods are listed from the best (leftmost) to the worst in each metric: the root mean square error (RMSE) (A), the normalized RMSE (B), the right-RMSE (C), the normalized discounted cumulative gain (NDCG) (D), the Pearson correlation coefficient (PCC) (E) and the Spearman correlation coefficient (SCC) (F). The methods are indexed alphabetically: BMTMKL (1), CaDRReS (2), CDRscan (3), cwKBMF (4), DualNets (5), ENet (6), M-ENet (7), GENet-Lap (8), GENet-NLap (9), KRL (10), KRR (11), MACAU (12), MERGE (13), MVLR (14), pairwiseMKL (15), RF-g (16), RF-gs (17), RWENet-both (18), RWENet-left (19), RWENet-right (20), SRMF (21), TANDEM (22). This dataset was not sufficient for L-RMSE and hence the evaluation for R-RMSE and LR-RMSE were identical.

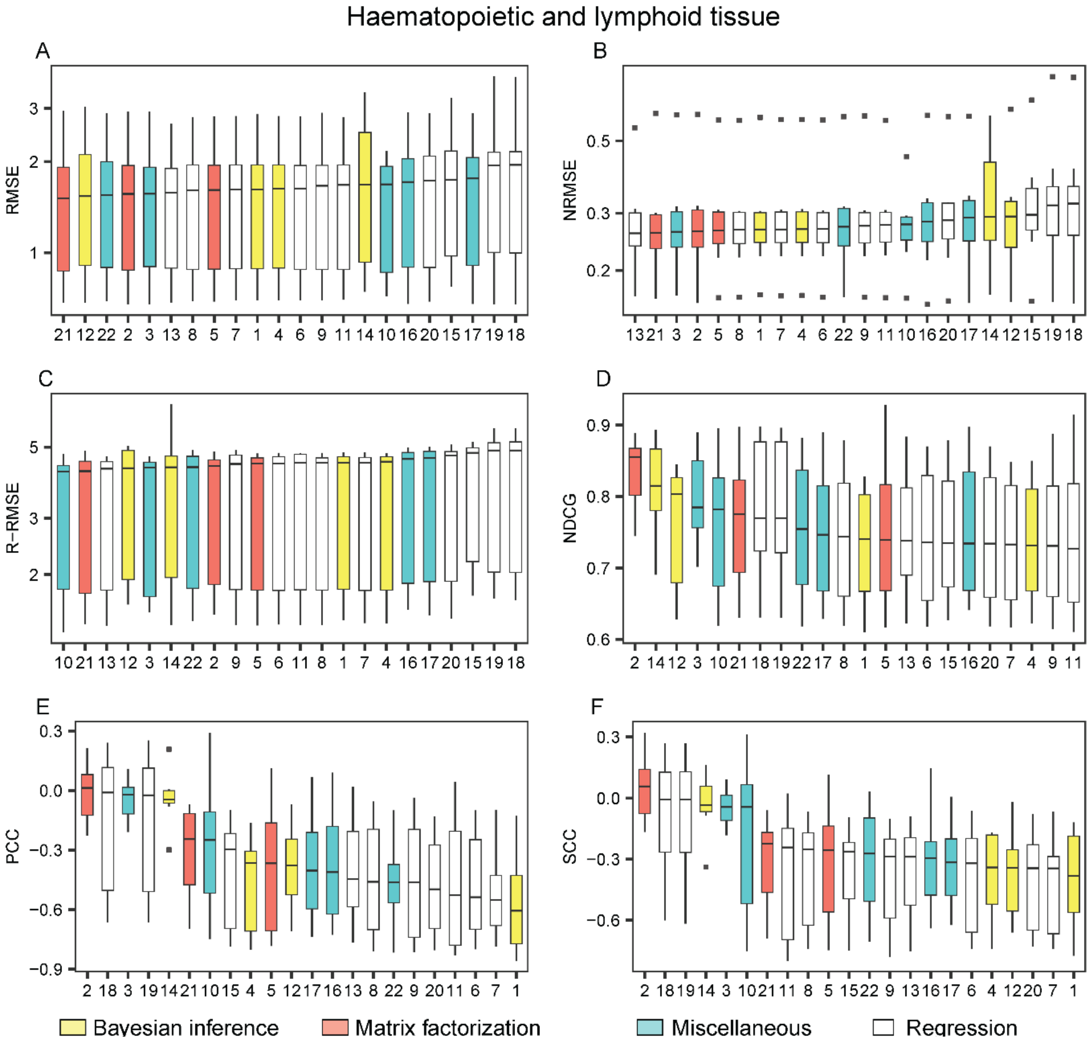


**Figure S48. The performance of the methods on the cell lines of haematopoietic and lymphoid tissue when trained on the GDSC AUC dataset and tested on the CCLE AUC dataset**. The methods are listed from the best (leftmost) to the worst in each metric: the root mean square error (RMSE) (A), the normalized RMSE (B), the right-RMSE (C), the normalized discounted cumulative gain (NDCG) (D), the Pearson correlation coefficient (PCC) (E) and the Spearman correlation coefficient (SCC) (F). The methods are indexed alphabetically: BMTMKL (1), CaDRReS (2), CDRscan (3), cwKBMF (4), DualNets (5), ENet (6), M-ENet (7), GENet-Lap (8), GENet-NLap (9), KRL (10), KRR (11), MACAU (12), MERGE (13), MVLR (14), pairwiseMKL (15), RF-g (16), RF-gs (17), RWENet-both (18), RWENet-left (19), RWENet-right (20), SRMF (21), TANDEM (22). This dataset was not sufficient for L-RMSE and hence the evaluation for R-RMSE and LR-RMSE were identical.

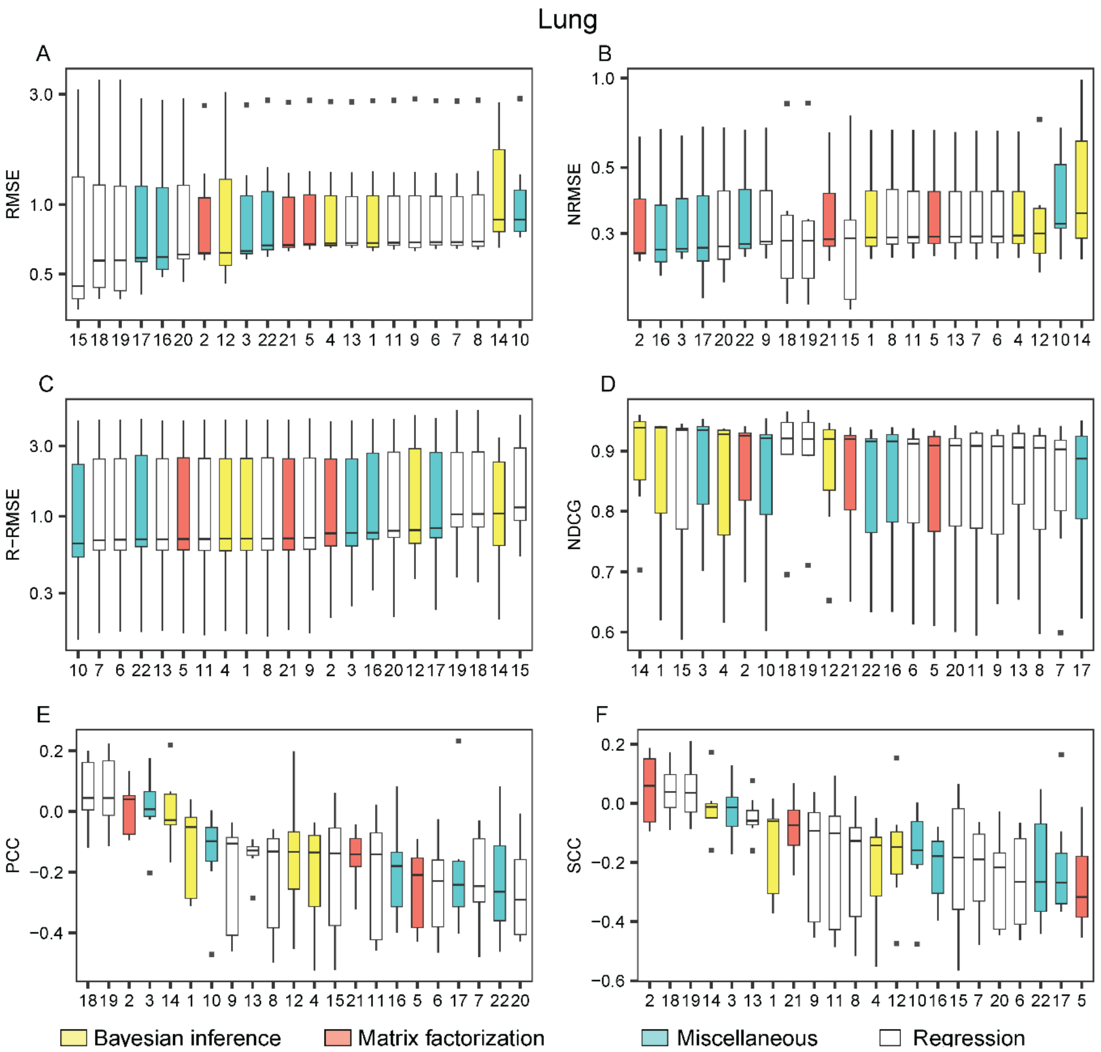


**Figure S49. The performance of the methods on the lung cell lines when trained on the GDSC AUC dataset and tested on CCLE AUC dataset**. The methods are listed from the best (leftmost) to the worst in each metric: the root mean square error (RMSE) (A), the normalized RMSE (B), the right-RMSE (C), the normalized discounted cumulative gain (NDCG) (D), the Pearson correlation coefficient (PCC) (E) and the Spearman correlation coefficient (SCC) (F). The methods are indexed alphabetically: BMTMKL (1), CaDRReS (2), CDRscan (3), cwKBMF (4), DualNets (5), ENet (6), M-ENet (7), GENet-Lap (8), GENet-NLap (9), KRL (10), KRR (11), MACAU (12), MERGE (13), MVLR (14), pairwiseMKL (15), RF-g (16), RF-gs (17), RWENet-both (18), RWENet-left (19), RWENet-right (20), SRMF (21), TANDEM (22). This dataset was not sufficient for L-RMSE and hence the evaluation for R-RMSE and LR-RMSE were identical.

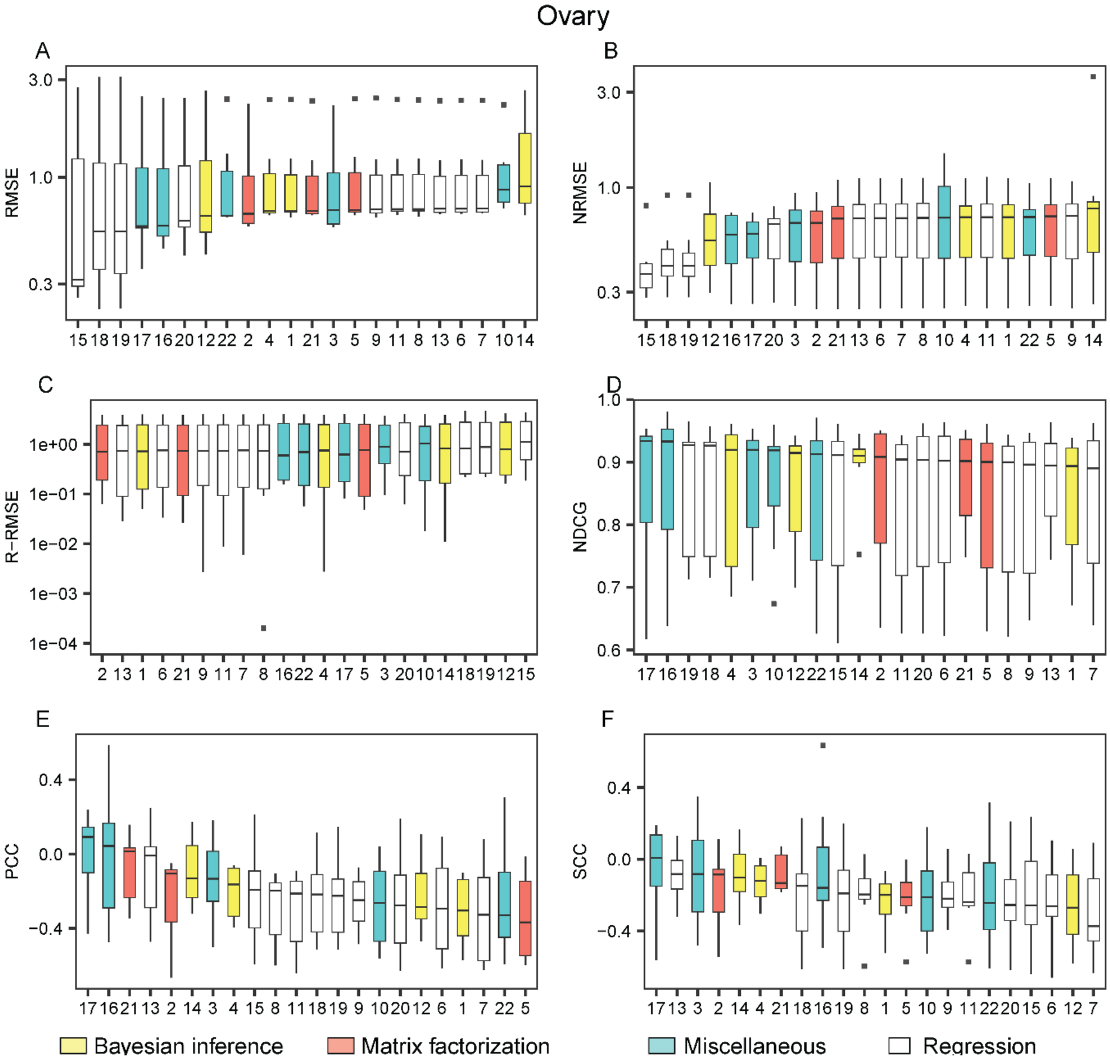


**Figure S50. The performance of the methods on the ovary cell lines when trained on the GDSC AUC dataset and tested on CCLE AUC dataset**. The methods are listed from the best (leftmost) to the worst in each metric: the root mean square error (RMSE) (A), the normalized RMSE (B), the left-RMSE (C), the right-RMSE (D), the left-right-RMSE (E), the normalized discounted cumulative gain (NDCG), the Pearson correlation coefficient (PCC) (G) and the Spearman correlation coefficient (SCC) (H). The methods are indexed alphabetically: BMTMKL (1), CaDRReS (2), CDRscan (3), cwKBMF (4), DualNets (5), ENet (6), M-ENet (7), GENet-Lap (8), GENet-NLap (9), KRL (10), KRR (11), MACAU (12), MERGE (13), MVLR (14), pairwiseMKL (15), RF-g (16), RF-gs (17), RWENet-both (18), RWENet-left (19), RWENet-right (20), SRMF (21), TANDEM (22). This dataset was not sufficient for L-RMSE and hence the evaluation for R-RMSE and LR-RMSE were identical.

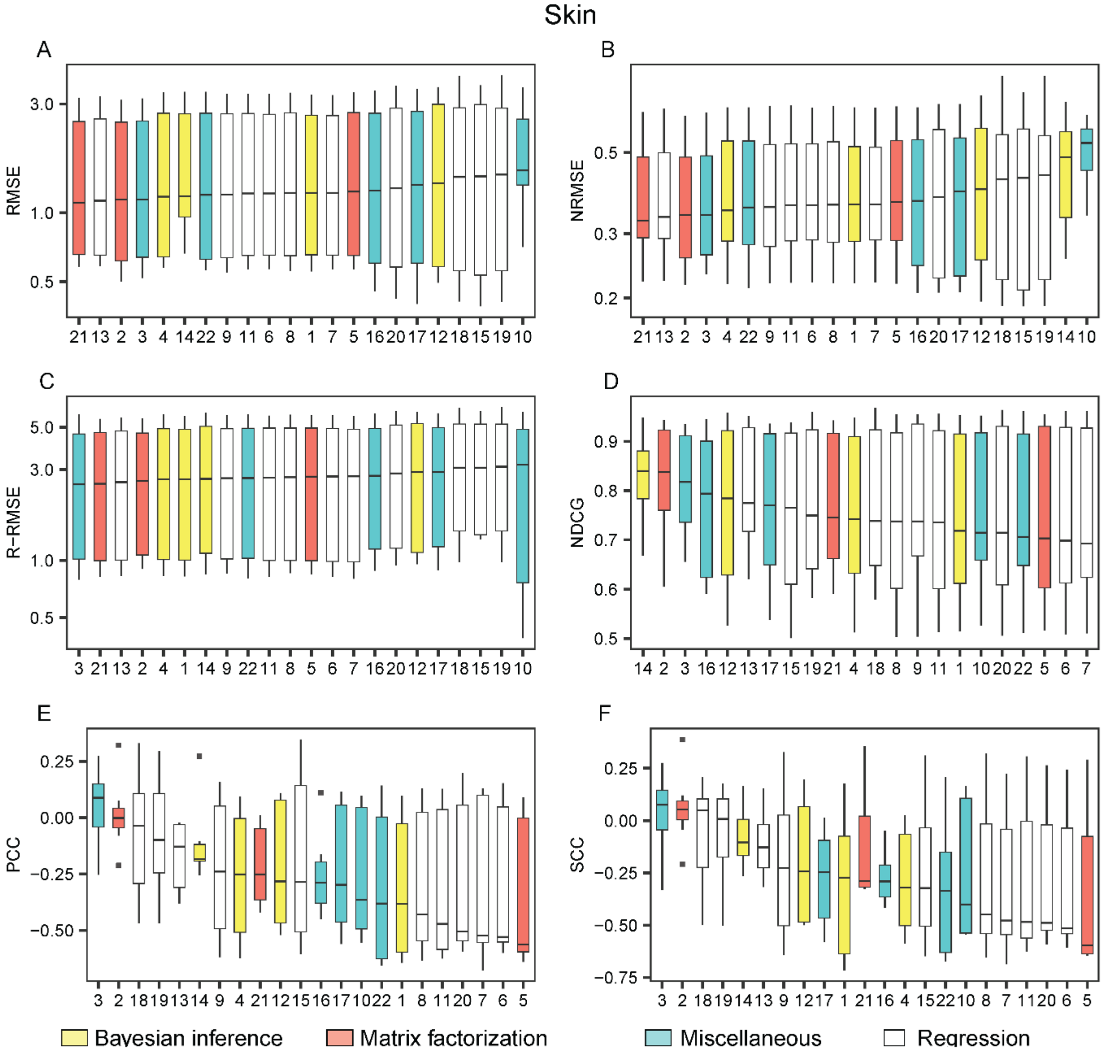


**Figure S51. The performance of the methods on the skin cell lines when trained on the GDSC AUC dataset and tested on CCLE AUC dataset**. The methods are listed from the best (leftmost) to the worst in each metric: the root mean square error (RMSE) (A), the normalized RMSE (B), the left-RMSE (C), the right-RMSE (D), the left-right-RMSE (E), the normalized discounted cumulative gain (NDCG), the Pearson correlation coefficient (PCC) (G) and the Spearman correlation coefficient (SCC) (H). The methods are indexed alphabetically: BMTMKL (1), CaDRReS (2), CDRscan (3), cwKBMF (4), DualNets (5), ENet (6), M-ENet (7), GENet-Lap (8), GENet-NLap (9), KRL (10), KRR (11), MACAU (12), MERGE (13), MVLR (14), pairwiseMKL (15), RF-g (16), RF-gs (17), RWENet-both (18), RWENet-left (19), RWENet-right (20), SRMF (21), TANDEM (22). This dataset was not sufficient for L-RMSE and hence the evaluation for R-RMSE and LR-RMSE were identical.
