## Supplementary File 8 for "A Survey and Systematic Assessment of Computational Methods for Drug Response Prediction"

**Figure S52**. The performance of the methods if trained on the GDSC AUC dataset and tested on the CTRP AUC dataset under different metrics.

**Figure S53**. The performance of the methods within five common drug groups when trained on the GDSC AUC dataset and tested on the CTRP AUC dataset under different metrics.

**Figure S54**. The performance of the methods on the cell lines of centeral nervous systems when trained on the GDSC AUC dataset and tested on the CTRP AUC dataset

**Figure S55**. The performance of the methods on the cell lines of haematopoietic and lymphoid tissue when trained on the GDSC AUC dataset and tested on the CTRP AUC dataset..

**Figure S56**. The performance of the methods on the large intestine cell lines when trained on the GDSC AUC dataset and tested on CTRP AUC dataset.

**Figure S57**. The performance of the methods on the lung cell lines when trained on the GDSC AUC dataset and tested on CTRP AUC dataset.

**Figure S58**. The performance of the methods on the skin cell lines when trained on the GDSC AUC dataset and tested on CTRP AUC dataset

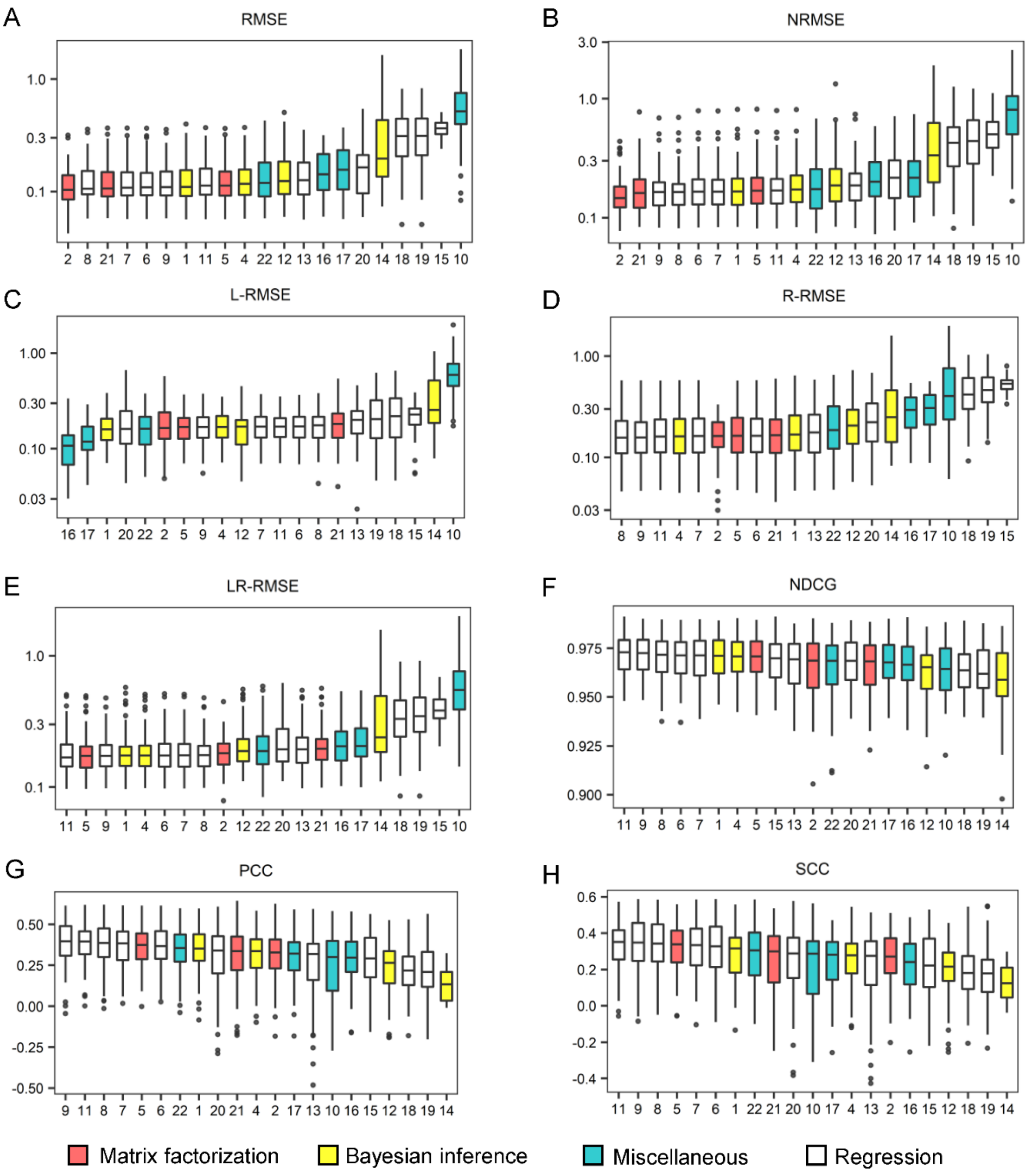


**Figure S52. The performance of the methods if trained on the GDSC AUC dataset and tested on the CTRP AUC dataset under different metrics.** Box plots were drawn for the root mean square error (RMSE) (A), the normalized RMSE (B), the left-RMSE (C), the right-RMSE (D), the left-right-RMSE (E), the normalized discounted cumulative gain (NDCG) (F), the Pearson correlation coefficient (PCC) (G) and the Spearman correlation coefficient (SCC) (H), in which the methods are listed from the best (leftmost) to the worst. Each figure was made on the basis of the scores of 10 repeated cross validation tests for each method. The methods are indexed alphabetically: BMTMKL (1), CaDRReS (2), CDRscan (3), cwKBMF (4), DualNets (5), ENet (6), M-ENet (7), GENet-Lap (8), GENet-NLap (9), KRL (10), KRR (11), MACAU (12), MERGE (13), MVLR (14), pairwiseMKL (15), RF-g (16), RF-gs (17), RWENet-both (18), RWENet-left (19), RWENet-right (20), SRMF (21), TANDEM (22).

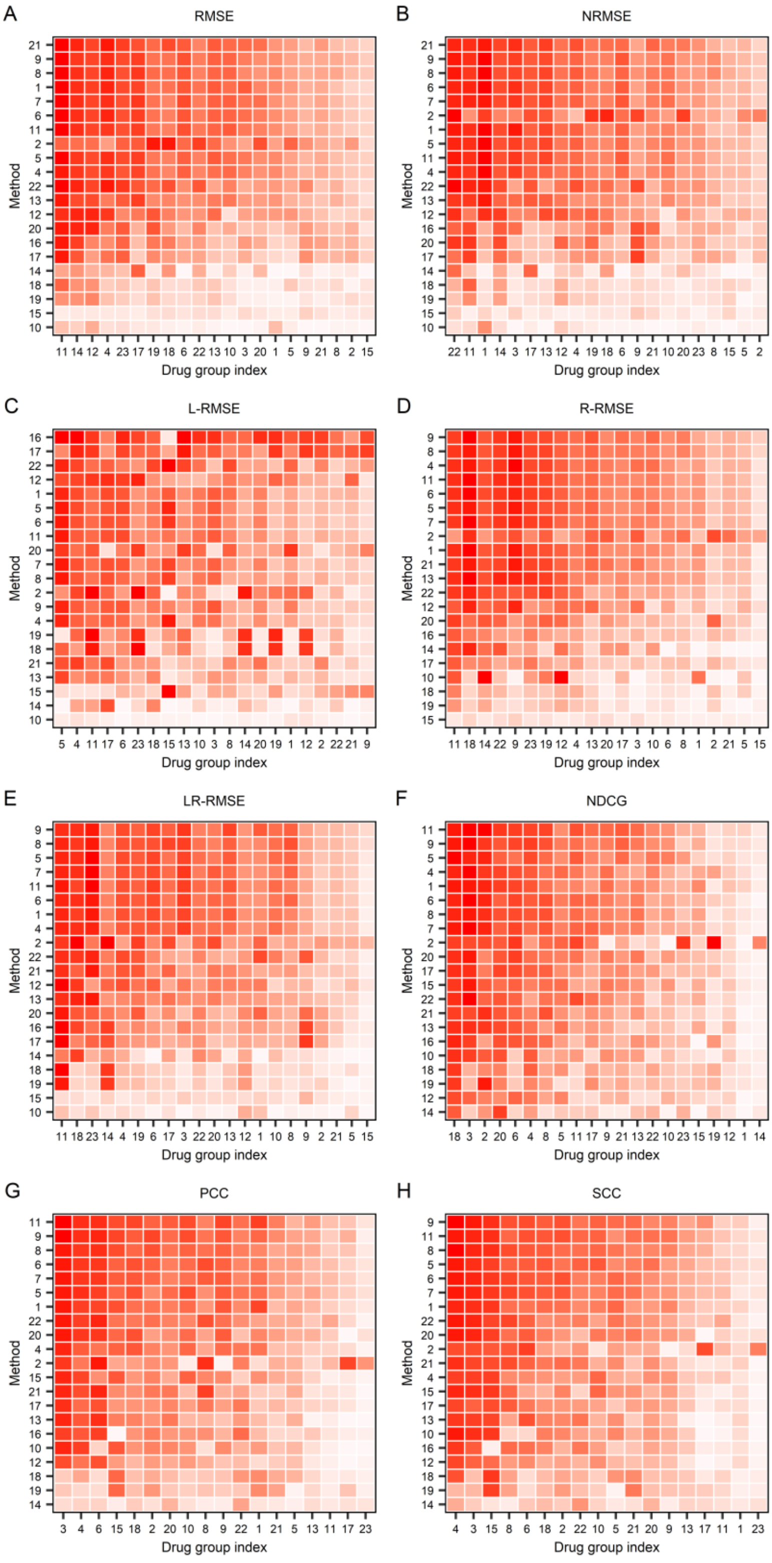


**Figure S53**. **Performance of the methods across 21 common drug groups if trained on the GDSC AUC dataset and tested on the CTRP AUC drug response dataset under different metrics.** (Cont'd on next page)

(Cont'd) The root mean square error (RMSE) (A), the normalized RMSE (B), the right-RMSE (C), the normalized discounted cumulative gain (NDCG) (D), the Pearson correlation coefficient (PCC) (E), and Spearman correlation coefficient (SCC) (F). Each row and column represent one method and one drug group, respectively. For the method (row) i and drug group (column) j, the median of the measures for drugs in the group j computed by the method I is used to color the corresponding cell, where dark red represents good performance, whereas light red represents bad performance. The methods are indexed alphabetically: BMTMKL (1), CaDRReS (2), CDRscan (3), cwKBMF (4), DualNets (5), ENet (6), M-ENet (7), GENet-Lap (8), GENet-NLap (9), KRL (10), KRR (11), MACAU (12), MERGE (13), MVLR (14), pairwiseMKL(15), RF-g (16), RF-gs (17), RWENet-both (18), RWENet-left (19), RWENet-right (20), SRMF (21), TANDEM (22). Drug groups are also indexed as follows: ABL signaling (1), Apoptosis regulation(2), Cell cycle(3), Chromatin histone acetylation(4), Chromatin histone methylation(5), Chromatin other(6), DNA replication(8), EGFR signaling(9), ERK MAPK signaling(10), Genome integrity(11), Hormone-related(12), IGFR signaling(13), JNK and p38 signaling(14), Metabolism(15), Other(17), Kinases(18), p53 pathway(19), PI3K/MTOR signaling(20), Protein stability and degradation(21), RTK signaling(22), WNT signaling (23). Drug 7 and 16 were not in the figure.

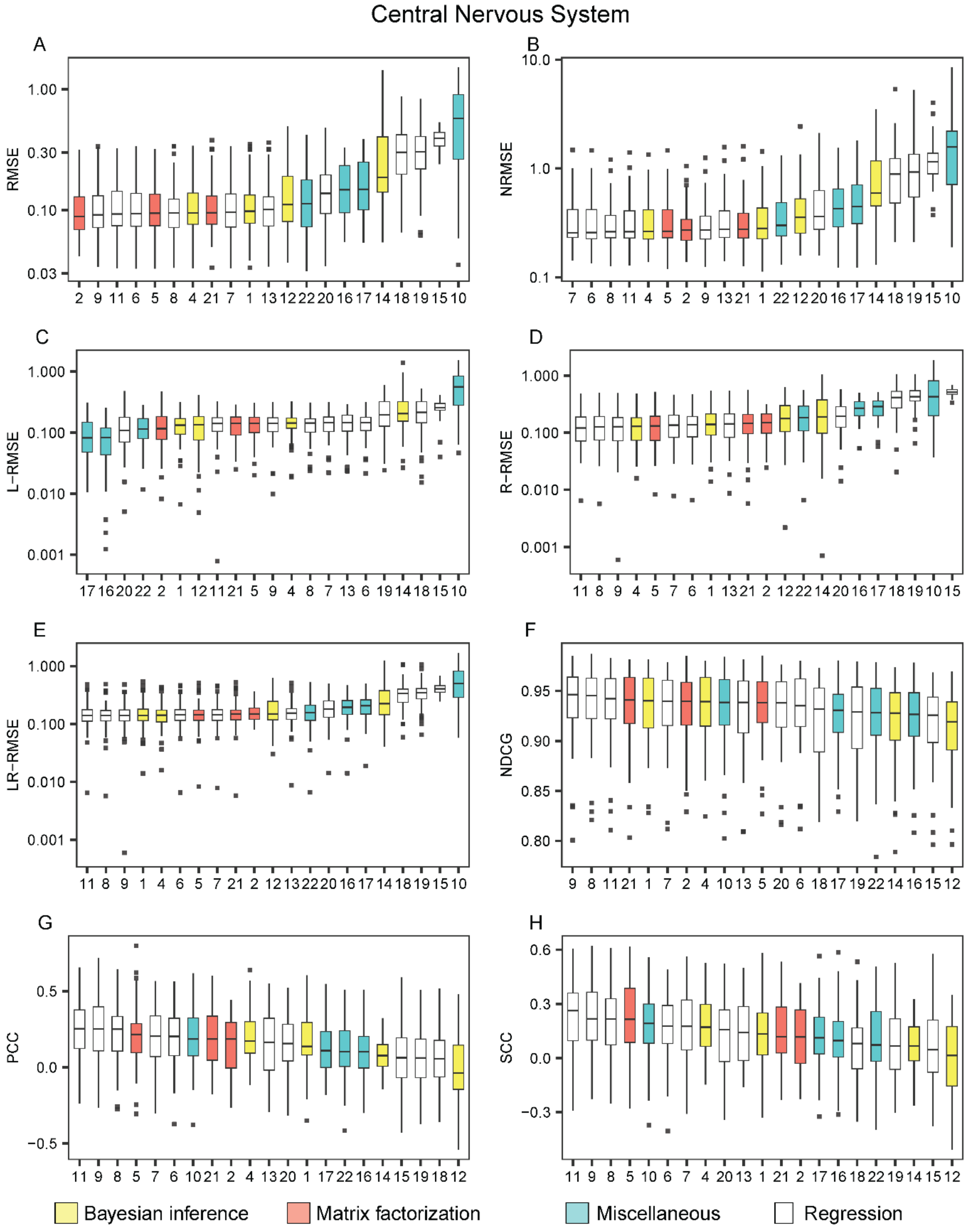


**Figure S54. The performance of the methods on the cell lines of the central nervous system when trained on the GDSC AUC dataset and tested on the CTRP AUC dataset**. The methods are listed from the best (leftmost) to the worst in each metric: the root mean square error (RMSE) (A), the normalized RMSE (B), the left-RMSE (C), the right-RMSE (D), the left-right-RMSE (E), the normalized discounted cumulative gain (NDCG), the Pearson correlation coefficient (PCC) (G) and the Spearman correlation coefficient (SCC) (H). The methods are indexed alphabetically: BMTMKL (1), CaDRReS (2), CDRscan (3), cwKBMF (4), DualNets (5), ENet (6), M-ENet (7), GENet-Lap (8), GENet-NLap (9), KRL (10), KRR (11), MACAU (12), MERGE (13), MVLR (14), pairwiseMKL (15), RF-g (16), RF-gs (17), RWENet-both (18), RWENet-left (19), RWENet-right (20), SRMF (21), TANDEM (22).

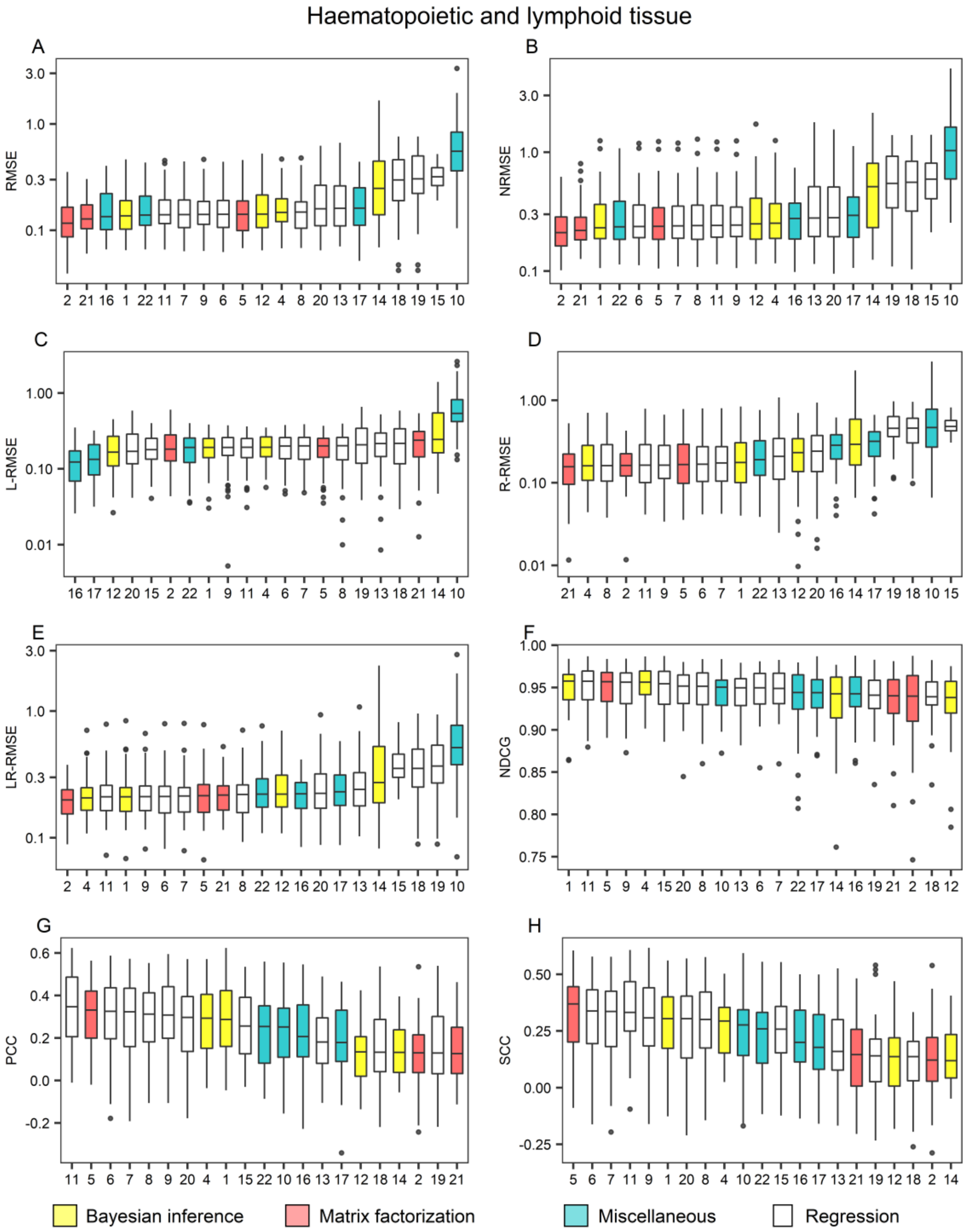


**Figure S55. The performance of the methods on cell lines of haematopoietic and lymphoid tissue when trained on the GDSC AUC dataset and tested on the CTPR AUC dataset.** The methods are listed from the best (leftmost) to the worst in each metric: the root mean square error (RMSE) (A), the normalized RMSE (B), the left-RMSE (C), the right-RMSE (D), the left-right-RMSE (E), the normalized discounted cumulative gain (NDCG) (F), the Pearson correlation coefficient (PCC) (G) and the Spearman correlation coefficient (SCC) (H). The methods are indexed alphabetically: BMTMKL (1), CaDRReS (2), CDRscan (3), cwKBMF (4), DualNets (5), ENet (6), M-ENet (7), GENet-Lap (8), GENet-NLap (9), KRL (10), KRR (11), MACAU (12), MERGE (13), MVLR (14), pairwiseMKL (15), RF-g (16), RF-gs (17), RWENet-both (18), RWENet-left (19), RWENet-right (20), SRMF (21), TANDEM (22).

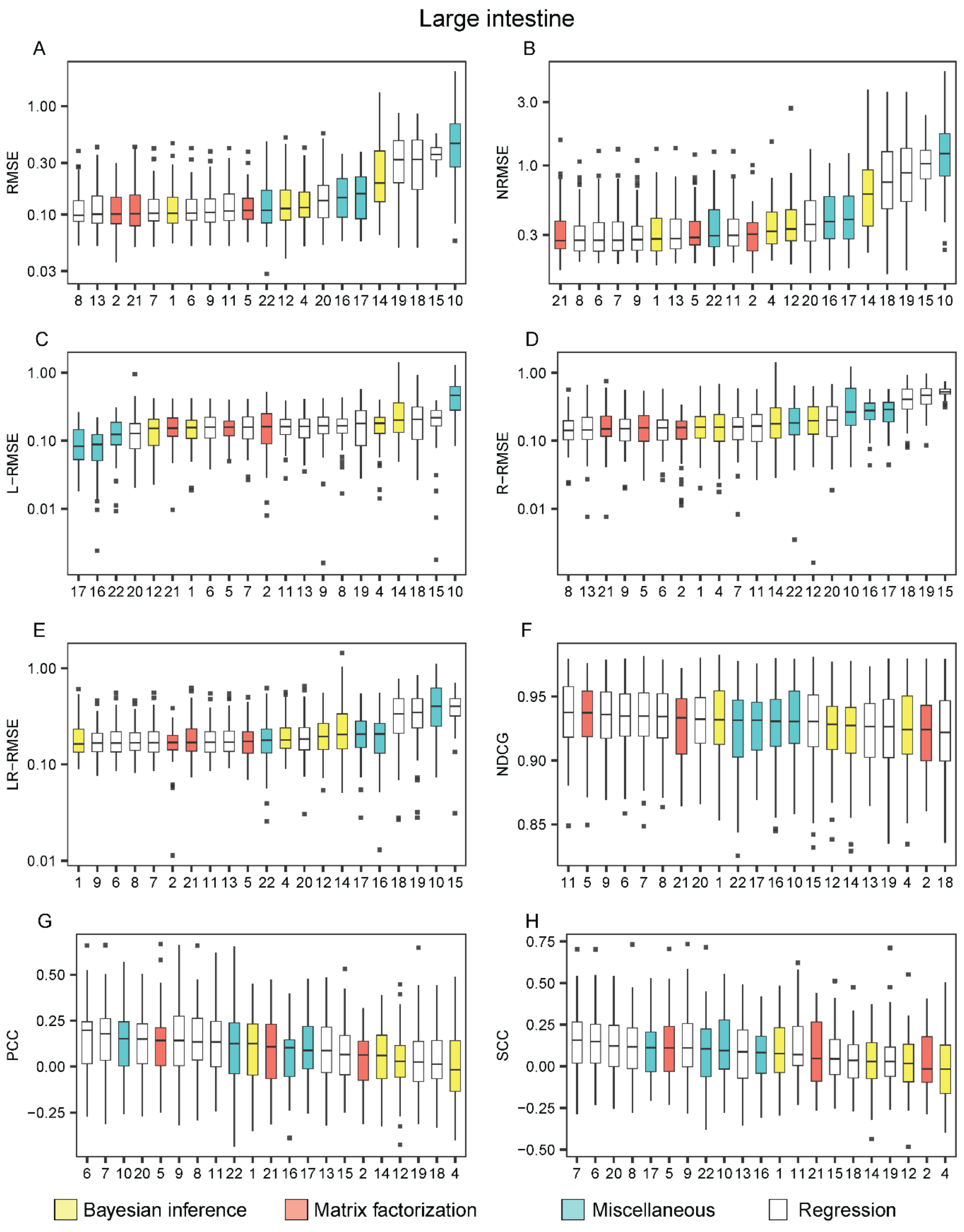


**Figure S56. The performance of the methods on the large intestine cell lines when trained on the GDSC AUC dataset and tested on CTRP AUC dataset**. The methods are listed from the best (leftmost) to the worst in each metric: the root mean square error (RMSE) (A), the normalized RMSE (B), the left-RMSE (C), the right-RMSE (D), the left-right-RMSE (E), the normalized discounted cumulative gain (NDCG), the Pearson correlation coefficient (PCC) (G) and the Spearman correlation coefficient (SCC) (H). The methods are indexed alphabetically: BMTMKL (1), CaDRReS (2), CDRscan (3), cwKBMF (4), DualNets (5), ENet (6), M-ENet (7), GENet-Lap (8), GENet-NLap (9), KRL (10), KRR (11), MACAU (12), MERGE (13), MVLR (14), pairwiseMKL (15), RF-g (16), RF-gs (17), RWENet-both (18), RWENet-left (19), RWENet-right (20), SRMF (21), TANDEM (22).

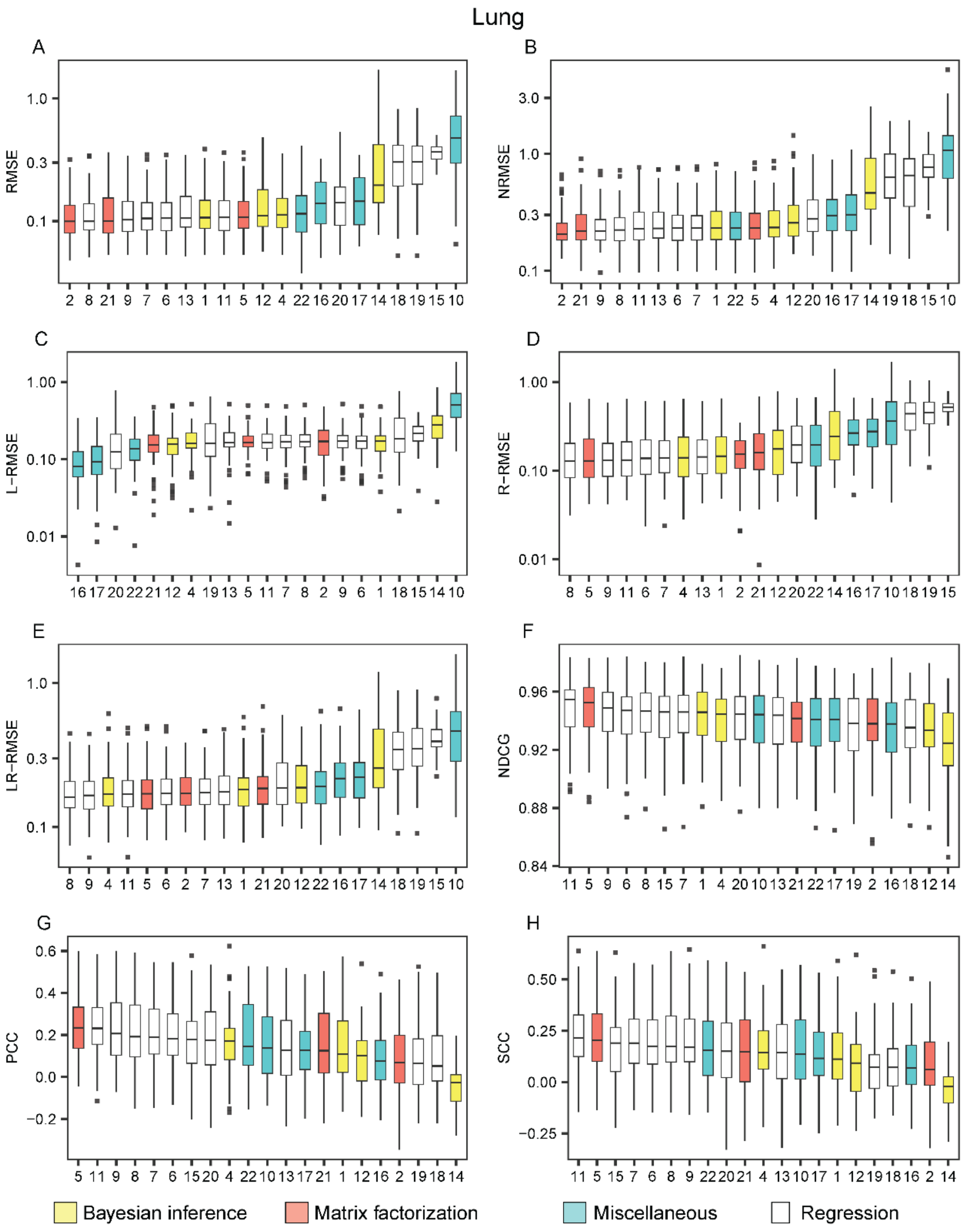


**Figure S57. Performance of the methods on the lung cell lines when trained on the GDSC AUC dataset and tested on CTRP AUC dataset**. The methods are listed from the best (leftmost) to the worst in each metric: the root mean square error (RMSE) (A), the normalized RMSE (B), the left-RMSE (C), the right-RMSE (D), the left-right-RMSE (E), the normalized discounted cumulative gain (NDCG), the Pearson correlation coefficient (PCC) (G) and the Spearman correlation coefficient (SCC) (H). The methods are indexed alphabetically: BMTMKL (1), CaDRReS (2), CDRscan (3), cwKBMF (4), DualNets (5), ENet (6), M-ENet (7), GENet-Lap (8), GENet-NLap (9), KRL (10), KRR (11), MACAU (12), MERGE (13), MVLR (14), pairwiseMKL (15), RF-g (16), RF-gs (17), RWENet-both (18), RWENet-left (19), RWENet-right (20), SRMF (21), TANDEM (22).

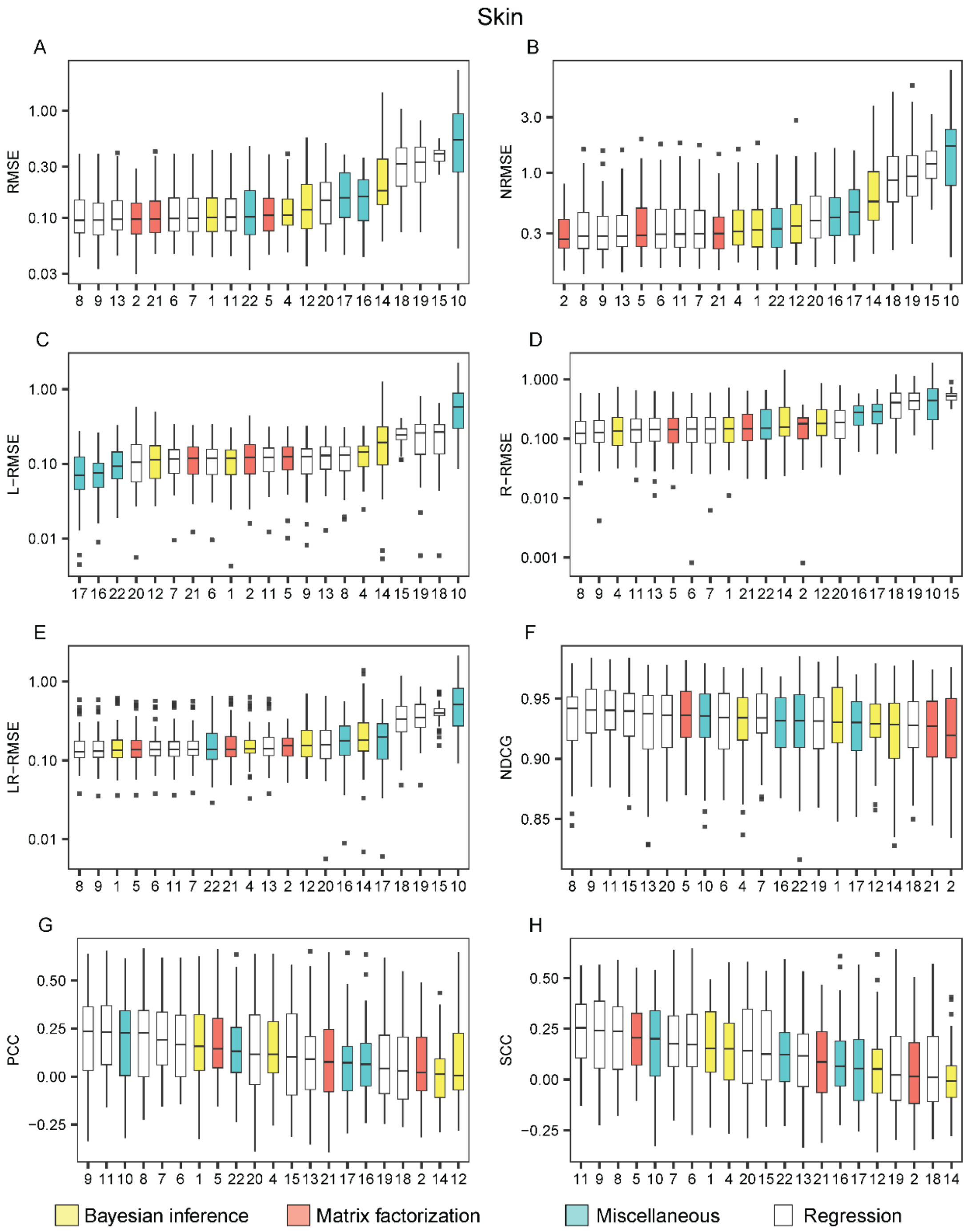


**Figure S58. The performance of the methods on the skin cell lines when trained on the GDSC AUC dataset and tested on CTRP AUC dataset**. The methods are listed from the best (leftmost) to the worst in each metric: the root mean square error (RMSE) (A), the normalized RMSE (B), the left-RMSE (C), the right-RMSE (D), the left-right-RMSE (E), the normalized discounted cumulative gain (NDCG), the Pearson correlation coefficient (PCC) (G) and the Spearman correlation coefficient (SCC) (H). The methods are indexed alphabetically: BMTMKL (1), CaDRReS (2), CDRscan (3), cwKBMF (4), DualNets (5), ENet (6), M-ENet (7), GENet-Lap (8), GENet-NLap (9), KRL (10), KRR (11), MACAU (12), MERGE (13), MVLR (14), pairwiseMKL (15), RF-g (16), RF-gs (17), RWENet-both (18), RWENet-left (19), RWENet-right (20), SRMF (21), TANDEM (22).
